# Large-scale skin metagenomics reveals extensive prevalence, coordination, and functional adaptation of skin microbiome dermotypes across body sites

**DOI:** 10.1101/2025.04.24.650393

**Authors:** Chengchen Li, Aarthi Ravikrishnan, Indrik Wijaya, Ahmad Nazri Mohamed Naim, Jean-Sebastien Gounot, Stephen Wearne, Yue Yuan On, Eliza Xin Pei Ho, Eileen Png, Junmei Samantha Kwah, Shree Harsha Vijaya Chandra, Qi Yan Ang, Minghao Chia, Ng Hui Qi Amanda, Srinivas Ramasamy, Yik Weng Yew, Marie Loh, Ruo Yan Ong, Faith Ng, Sze Han Lee, James Chan, The HELIOS Study team, Viduthalai Rasheedkhan Regina, Thomas Dawson, John Chambers, John Common, Niranjan Nagarajan

## Abstract

While skin microbiome studies have increasingly highlighted its importance in health and disease, our understanding of inter-individual heterogeneity in structure and function remains limited, impacting the ability to develop microbiome-based stratification and therapeutics. Powered by comprehensive skin microbiome characterization in a multi-ethnic population-based cohort (>3,550 shotgun metagenomes across 18 sampling sites), we established significant undescribed inter-individual heterogeneity and the extensive prevalence of distinct microbial configurations (17 species-resolution dermotypes) in seven out of nine body sites. Combining functional *in silico* and *in vitro* studies revealed insights into how these dermotypes assemble as a function of niche-dependent microbial interactions (e.g. hypoxia-dependent inhibition of *S. hominis* by *S. epidermidis*/*M. luteus*) and metabolic resource utilization (e.g. differential galactose and histidine metabolism). Integration of demographic, skin physiological, and behavioral data further identified >30 significant associations with host attributes. Cross-site analysis revealed remarkable coordination across disparate skin regions (predictive AUC-ROC>0.8) and bilateral consistency (Pearson ρ>0.95), emphasizing the role of specific microbial and host factors in shaping dermotypes. Finally, we provide multiple lines of evidence that dermotype states impact the risk for skin discomfort (e.g. irritation, itch) and diseases (e.g. eczema), that when combined with our highly accurate dermotype classifiers (AUC-ROC>0.98), provide a new paradigm for understanding skin microbiome function and stratifying patients in the context of skin and other diseases.

## Introduction

The skin microbiome, a diverse community of bacterial, viral, and eukaryotic microbes, plays a vital role in human health by contributing to skin barrier function, preventing pathogen colonization and infections, and maintaining immune homeostasis^1–3^. Alterations in the skin microbiome are associated with a range of common skin diseases^4–17^ (e.g. eczema, acne, psoriasis) and discomforts^18–23^ (e.g. itch, dryness, body odor), with a significant impact on quality of life^24^. Despite groundbreaking early studies highlighting differences in skin microbiome composition across body sites^25–28^ and long-term temporal stability^29^, our understanding of variability across individuals (i.e. population heterogeneity), and factors defining skin microbial ecology and function remain limited^30–33^. This knowledge is critical for developing targeted immune and microbiome therapies, and patient stratification to define disease risk, treatment response, and prognosis^34–36^.

The subject of population heterogeneity and distinct microbial community types has received notable attention in body sites such as the gut and the vaginal tract. For example, in the gut, a seminal study described three distinct enterotypes—stable microbiome configurations— associated with various host factors and health^37–39^. Recent studies have sought to challenge or refine this concept further with ideas such as entero-signatures and microbial guilds, which describe distinct sets of species that co-occur variably across individuals and define host-microbe interactions^40,41^. Similarly, vaginal microbiome studies have consistently identified five distinct community types and linked them with personal hygiene, lifestyle, and reproductive health^42–46^.

While gut and vaginal microbiome community types have been established as important organizing principles and risk factors relevant to host health, skin microbiome studies have yielded comparatively limited evidence. Early work based on data from the Human Microbiome Project provided initial hints into the presence of skin microbiome community types^47,48^, but it relied on low-resolution bacterial 16S rRNA sequencing and lacked information on associations with microbial functions and skin phenotypes. Recent shotgun metagenomics-based studies have offered more detailed information about microbial species and functions (including key fungal species), but have been limited to single-site, disease/dysbiosis contexts^49–51^ (e.g. eczema or air pollution). Correspondingly, we know little about skin microbiome community types (that we refer to as dermotypes) in the general population, and the roles that microbial and host functions play in shaping them remain unestablished. Furthermore, whether dermotypes are coordinated across the entire skin organ system, and whether dermotype states can influence host phenotypes including skin discomfort and disease, remains unknown.

Here we address these knowledge gaps by comprehensively exploring skin microbiomes in 18 different sampling sites of a multi-ethnic cohort of 200 adults representing the general population across the spectrum of health and disease. We captured a broad range of parameters by combining shotgun metagenomic sequencing data (n=3,600 samples), demographic information (e.g., age, ethnicity, etc.), skin physiological measurements (e.g., skin pH, hydration, etc.), and extensive skin health questionnaire data (e.g., skin care and skin discomfort/disease related questions). Integrated analysis of this data revealed substantial population heterogeneity, and extensive prevalence of dermotypes across body sites (7 out of 9 sites, including five different dermotypes in the axilla). Remarkably, we discovered systemic coordination of dermotypes across physiologically distinct sites (e.g., dry forearm and sebaceous forehead), challenging the independence of site specificity in the skin microbiome and suggestive of a globally coordinated system. We derived mechanistic insights into dermotype emergence and established how microbiome-intrinsic factors, including differences in species co-abundance patterns, interspecies interactions (e.g., hypoxia-dependent inhibition of *S. hominis* by *S. epidermidis* and *M. luteus*), and microbial metabolic pathways (e.g., galactose utilization favoring *S. hominis* over *S. epidermidis*), can shape dermotype structures. Host factors such as demographics were found to be informative for dermotype prediction in only a few body sites (e.g., elbow crease, mean AUC-ROC=0.79), whereas microbial features such as metabolic functions and dermotypes from other body sites typically had greater predictive power. We trained near-perfect classifiers for dermotype labels (AUC-ROC>0.98), with as few as five species in all cases, underscoring the potential for dermotype identification with just a few signature species. Overall, this work establishes the importance of accounting for population heterogeneity in skin microbiome investigations, while highlighting specific microbial and host factors that shape dermotypes, which in turn can impact skin discomforts and diseases. We further provide a clinically actionable framework for assigning skin microbiome states to individuals, facilitating the development of patient stratification strategies and personalized treatments.

## Results

### Analysis of inter-individual heterogeneity in the skin microbiome identifies robust dermotype states across multiple body sites

To capture skin microbiome heterogeneity across individuals and different body sites in the general population, we generated an extensive collection of skin samples (n=3,600) from 18 different sampling sites in 200 adults recruited from the Health For Life In Singapore (HELIOS) precision medicine cohort^52^ (**Figure 1A**, **Supplementary Table 1**; **Methods**). The samples included the left and right sides of nine sites of importance for skin health and disease^17,53,54^, including scalp (Sc), parietal scalp (Ps), forehead (Fo), cheek (Ch), upper back (Ub), axilla (Ax), antecubital fossa/elbow crease (Ac), volar forearm (Vf), and lower leg (Ll; **Figure 1A**). A substantial fraction of the samples (3,586/3,600=99.6%) were successfully and consistently processed for the construction of metagenomic libraries and sequenced on an Illumina HiSeq platform to generate >70 billion reads and >10Tbp of data in total (2×150bp reads, median of 21.6 million reads and 3.3Gbp of data per library). The resulting metagenomic datasets were processed to filter reads for quality, identify the non-human fraction (median of 7.4 million reads and 1.1Gbp per library), and conduct microbial taxonomic profiling. Data from negative controls (n=46; median of 7.3 million non-human reads; **Supplementary File 1**) were integrated to identify and exclude contaminant taxa (**Supplementary File 2**) and remove poor quality profiles (**Supplementary Figure 1**; **Methods**). Separately, we collected extensive participant data to investigate associations between inter-individual skin microbiome variations and host factors, including demographic information (age, gender, ethnicity, etc.), skin physiological measurements (sebum level, pH, etc.), skin-care practice (hygiene, personal-care product usage), and skin health data (skin discomfort and diseases; **Supplementary Table 1**). This collection provides the largest consistently processed skin metagenome dataset for a population-wide cohort to date (n=3,572; **Supplementary File 3**), enabling in-depth comparisons of the skin microbiome across individuals and analysis of its implication for skin health.

**Figure 1.**
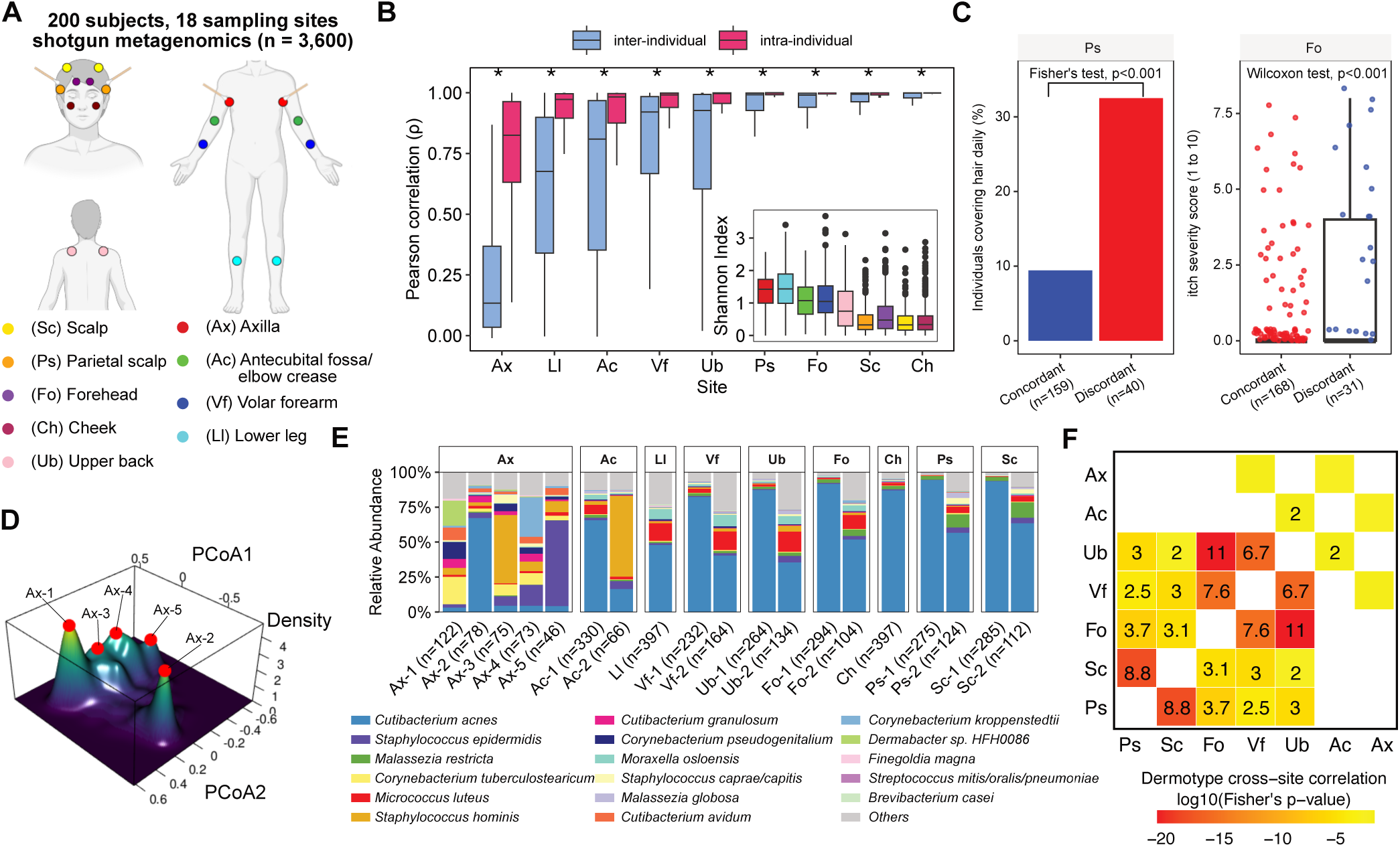
Heterogeneity of site-specific skin microbial community compositions within and across individuals. **A.** Overview of skin sampling strategy, illustrating bilateral sampling at nine sites with swabs for Ps and Ax and tapes for others. **B.** Boxplots quantifying the similarity of the skin microbiome within (left versus right) and between individuals across sites. Sites in the inset figure for alpha diversity are presented in the same order. All p-values were calculated using the one-sided Wilcoxon rank-sum test (* denotes BH-adjusted p-value<0.001). Boxplot whiskers represent 1.5× interquartile range or the maximum/minimum data point within the range. **C.** Left barplot comparing the percentage of subjects with their hair covered daily between those with concordant and discordant parietal scalp microbiomes between left and right body sites; right boxplot showing that individuals with discordant forehead microbiomes have significantly higher itch severity. **D**. Perspective kernel density plot of Ax microbiome PCoA results with the five red dots representing local maxima determined via Mean Shift analysis. **E.** Averaged relative abundance plots of major species ranked by median abundance for each dermotype or site, with the number of stable samples in each site and dermotype shown at the bottom. **F.** Heatmap showing significant correlations for dermotype labels across sites. The color gradient denotes log10-transformed BH-adjusted Fisher’s exact test p-values, and odds ratios are annotated in the boxes where appropriate.

Leveraging data availability from the left and right sides of different body sites (n=9) for many individuals (n=200), we first investigated the extent to which skin microbiomes exhibit bilateral symmetry, beyond observations in a few individuals^26^ and sites^5,49^. Despite factors such as dominant hand use and skin product usage not being symmetric, high bilateral concordance was observed for most individuals across body sites (median Pearson correlation>0.95, except for axilla; **Figure 1B**). At all nine body sites studied, skin microbiomes from the left and right sides of the body were significantly more similar than across individuals (Wilcoxon test p-value<0.001; **Figure 1B**). This pattern is robust and seen with the Yue-Clayton similarity index as well, which factors in shared and non-shared species^55^ (Wilcoxon test p-value<0.001, **Supplementary Figure 2A**), and after accounting for (within-sample) diversity (Shannon index, **Figure 1B**) and *Cutibacterium acnes* dominance^2^ as a potential confounder (Wilcoxon test p-value<0.001, **Supplementary Figure 2B**; **Methods**). In the background of the high general concordance observed, >14% of sites (255/1779) exhibited discordance in 121 individuals across different body sites (**Supplementary Figure 3A**; **Methods**). Intriguingly, bilateral discordance was enriched in some individuals (binomial test BH-adjusted p-value<0.05; **Supplementary Figure 3B**), where multiple body sites exhibited correlated patterns of bilateral discordance (e.g., head sites and upper back; **Supplementary Figure 3C**). Specific associations were also detected between bilateral microbiome discordance and various host attributes, including gender (e.g. cheek discordance in women; **Supplementary Figure 3D**), personal habits (e.g. scalp discordance and daily hair covering; **Figure 1C**), and skin conditions (e.g. forehead discordance and higher itch score; **Figure 1C**). Hence, while the adult skin microbiome is established here to be strongly bilaterally concordant across body sites, coordinated discordance at specific sites may be triggered by host-specific differences in the skin microenvironment arising out of behavioral and/or skin condition changes.

Aggregating skin microbiome taxonomic profiles across body sites revealed the broad influence of skin microenvironment on skin microbiome composition^27,28^, with profiles from the same site largely clustering together, and axilla (Ax) sites with their hypoxic and moist environments being the most distinct (**Supplementary Figure 4A**). While sebaceous head and scalp sites (Sc, Ps, Fo, and Ch) exhibited the highest similarity, even the two most similar scalp sites (Ps, Sc) had statistically significant differences in beta (between-sample) diversity (PERMANOVA adonis p-value<0.05; **Supplementary Figure 4B**). Site-specific differentially abundant species included enrichment in lipophilic *C. acnes* in sebaceous head sites^27^, the fungus *Malassezia globosa* in the elbow crease (Ac) and upper back^56^ (Ub), and *Malassezia restricta* in scalp sites^57,58^, as well as novel observations including *Cutibacterium avidum*, *Cutibacterium granulosum*, and various *Staphylococcus* and *Corynebacterium* species in the axilla (consistent with their preference for moist environments), and *Micrococcus luteus, Moraxella osloensis*, and common oral taxa (*Streptococcus pneumoniae*, *Corynebacterium matruchotii*, and *Rothia mucilaginosa*) in the forearm (Vf) and elbow crease (Ac; **Supplementary File 4**), defining the broad dimensions of skin microbes with preferences for site-specific environmental niches.

Despite these consistent differences across sites, we found that individual skin microbiomes from a given body site often exhibit multimodal distributions across principal axes of variation (e.g., Ac and Ax sites; **Supplementary Figure 4A**), suggesting substantial inter-individual heterogeneity in the population. Exploiting the availability of a large number of consistently processed taxonomic profiles for each site (n>395), we developed a systematic clustering strategy to identify robust stratification in the data, as well as assess stability of cluster membership via bootstrapping (n=100; **Supplementary Figure 5**; **Methods**). Overall, seven out of nine sites (i.e. Ax, Ac, Vf, Ub, Fo, Ps, Sc) showed clear evidence for multiple robust clusters we term dermotypes, with all clusters having a median stability >90%. All identified dermotypes were found to have prevalence in the population >10%, with the two dermotypes identified in the volar forearm both having prevalence >40%. Notably, axilla sites displayed the greatest diversity with five robust dermotypes, each represented by a high-density region of samples (**Figure 1D**). Additionally, we identified two dermotypes at each of the remaining sites except cheek and lower leg sites, where only a single robust cluster was detected (**Supplementary Figure 6A-F**). While *C. acnes*-enriched dermotypes were common (Ax-2, Ac-1, Vf-1, Ub-1, Fo-1, Ps-1, and Sc-1; **Figure 1E**) and with lower diversity (except Ac-1, Wilcoxon test p-value<0.05; **Supplementary Figure 7A**), other species including *Staphylococcus*, *Corynebacterium*, *Dermabacter, Streptococcus*, *Finegoldia*, *Brevibacterium,* and *Rothia* were found to be dominant or differentially abundant across dermotypes, highlighting their contributions to dermotype structure and function (**Supplementary Figure 7B**; **Supplementary File 5**). For example, the four dermotypes in the axilla were dominated either by combinations of *Corynebacterium* and *Dermabacter* species and with higher α-diversity (Ax-1, Ax-4), or two different *Staphylococcal* species (*Staphylococcus hominis* at Ax-3, *Staphylococcus epidermidis* at Ax-5) and with intermediate levels of α-diversity. Similarly, the elbow crease exhibited a *Staphylococcus*-dominant dermotype (Ac-2) seen in >15% of the general population, while scalp sites (Sc, Ps) exhibited an *M. restricta* enriched dermotype in >25% of the population, highlighting the diversity of species contributions (**Supplementary Figure 7B**). Of note, species-level relative abundances enabled the training of near-perfect classifiers for dermotype labels (AUC-ROC>0.98) with a few key signature species, emphasizing the feasibility of dermotype identification in the clinic with targeted assays (**Supplementary Figure 7C**; **Supplementary File 6**).

Building on the observation that skin microbiomes are bilaterally concordant, we next investigated whether dermotypes across different body sites exhibited consistent associations within individuals (**Figure 1F**). Indeed, remarkably strong associations were detected between dermotypes across body sites, between not just physically proximal sites (e.g., scalp and parietal scalp), and those with similar physiological characteristics (e.g., scalp and forehead), but also more distant sites with distinct physiology (e.g., dry volar forearm and sebaceous forehead sites; Fisher’s exact test adjusted p-value<0.05; **Figure 1F**). These associations are not explained by preferences for specific species; for example, *Staphylococcus*-dominant dermotypes in the elbow crease and axilla are negatively associated with each other (**Supplementary Figure 8A**), even though *Staphylococcus* strains are known to be transmissible via skin contact^59–61^. Furthermore, no strong correlations were observed for the abundance of *Staphylococcus* species between these sites (**Supplementary Figure 8B**), indicating that other systemic factors play a role in the observed dermotype associations. Overall, these results emphasize the presence of distinct dermotype configurations across individuals and body sites, with surprisingly high levels of bilateral and cross-site concordance that point to body-wide coordination of host and microbial factors shaping them.

### Dermotype-specific core and ancillary species exhibit coordinated patterns across body sites

To gain deeper insights into dermotype composition, we categorized species into dermotype-specific core (with ≥50% prevalence) and ancillary (<10% prevalence, ≥3 occurrences) species (**Figure 2A**; **Methods**). Across all dermotypes, this identified 15 core bacterial species from the genera *Cutibacterium*, *Staphylococcus*, *Corynebacterium*, *Micrococcus,* and *Moraxella*, and two core fungal species from the genus *Malassezia* (**Figure 2B**). No archaea or viruses were core among the 560 species analyzed, emphasizing their sporadic distribution in the human skin microbiome^29^. Each dermotype contained 2-12 core species, collectively accounting for >70% of community abundance, with *C. acnes* being a universal core species across all dermotypes and sites. However, most taxa were core in only some sites or dermotypes, for instance *Dermabacter sp. HFH0086,* which was categorized as a core species in the most common dermotype in the axilla (Ax-1, prevalence>77%) but rarely seen in the other dermotype (Ax-2, prevalence<2%; **Figure 2B**). Even abundant species such as *S. epidermidis* and *S. hominis* were not universally core, while some taxa with low relative abundances (median abundance<0.5%) were core in some dermotypes (e.g., *Corynebacterium tuberculostearicum*, *Corynebacterium pseudogenitalium*, *Staphylococcus caprae/capitis,* and *M. globosa*). This co-existence of high- and low-abundance core species within communities suggests that their collective maintenance may arise from niche partitioning or complementary functional roles, independent of an abundance effect^62^ (**Figure 2B**). Building on our observation of dermotype correlations between body sites (**Figure 1F**), we analyzed intra-individual correlations in core species abundances across the nine skin sites. Several taxa, such as *M. restricta*, *S. epidermidis,* and *C. acnes*, exhibited modest positive correlations when aggregated across all sites (spanning diverse microenvironments, mean ρ=0.26), and higher correlations when excluding a few sites (e.g. *M. restricta* excluding Ax, mean ρ=0.37; **Supplementary Figure 9A-D**). This suggests that core species shared across skin sites are coordinated body-wide within individuals, potentially mediated by mechanisms such as global sebum secretion favoring lipophiles, antibiotic resistance and strain dispersal^63^, or immune-driven niche filtering^64,65^. In contrast, *S. hominis* showed positive correlations limited to the non-scalp region (**Supplementary Figure 9E**), while *M. globosa* displayed divergent correlation patterns with positive associations among facial sites (ρ=0.28), but negative correlations between the cheek and upper back (ρ=-0.25; **Supplementary Figure 9F**). These heterogeneous patterns imply that systemic host/microbial drivers might interact with localized environmental variations (e.g., pH, humidity gradients) to modulate cross-site microbial assembly patterns.

**Figure 2.**
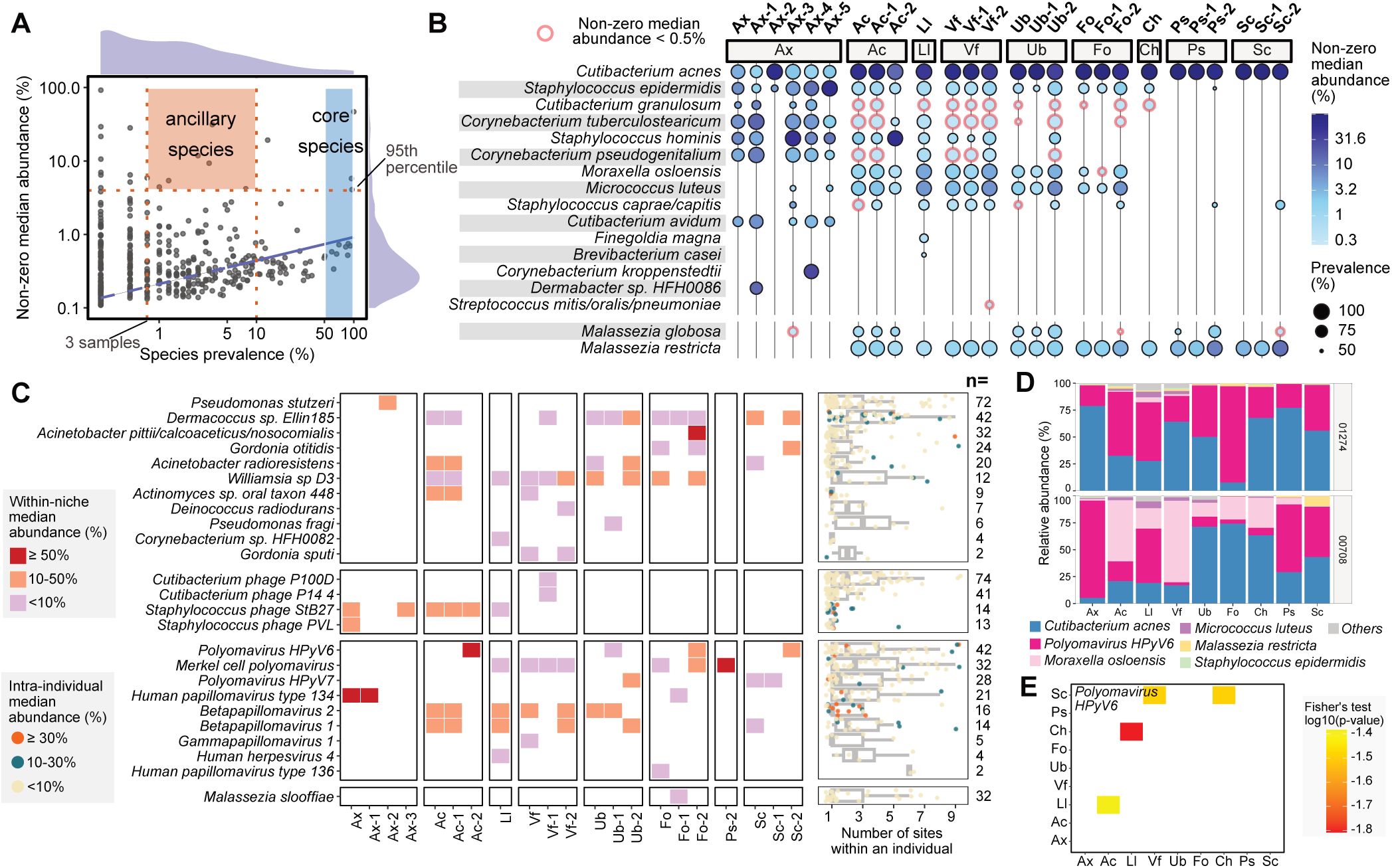
Dermotypes are characterized by distinct core and ancillary species. **A.** Overview of the general trend relating species prevalence and abundances (lower leg microbiomes as an example), and the schema to categorize species into *core species* (≥50% prevalence, in blue) and *ancillary species* (≥3 occurrences, <10% prevalence, ≥95^th^ percentile non-zero median abundance, in orange). **B.** Core species distribution by site and dermotype, with sizes corresponding to prevalences and color intensities corresponding to non-zero median abundance. Circles with red boundaries mark species with <0.5% median abundance. **C.** Left: Ancillary species distribution and non-zero median abundance at each site and dermotype. Right: Boxplots showing the intra-individual occurrence and median abundance of each ancillary species, with the number of individuals indicated on the right (each overlaid point represents one subject). The x-axis shows the number of sites (out of nine) where a given ancillary species is observed, and the color indicates its within-subject median abundance. **D.** Stacked barcharts depicting relative abundance profiles for selected subjects with blooms of *HPyV6* across multiple sites. **E.** Significant inter-site co-abundance patterns of *HPyV6*, with the color gradient denoting log10-transformed BH-adjusted Fisher’s exact test p-values.

The 25 ancillary species we identified were disproportionally abundant at various sites and dermotypes (in some cases >50% relative abundance) despite being, by definition, not overall very prevalent (<10%; **Figure 2C**; **Methods**). This group includes bacteria primarily from the *Pseudomonas*, *Acinetobacter*, and *Gordonia* genera (often environmentally resilient and biofilm forming^66–68^), along with bacteriophages of common skin bacteria (e.g., *Cutibacterium*, *Staphylococcus*), common skin viruses (e.g., polyomavirus and human papillomavirus), and a fungus (*Malassezia slooffiae*; **Figure 2C**). Nearly half of ancillary taxa (12/25=48%) showed specificity to a particular dermotype; for example, *Acinetobacter pittii/calcoaceticus/nosocomialis* was disproportionally abundant (median abundance>50%) only in the less prevalent forehead dermotype Fo-2, indicating that ancillary taxa may be adapted to specific niches available only in some individuals and microbiome configurations. This was further seen in the form of significant enrichment of ancillary species across multiple body sites of many individuals (≥2 sites in 87 individuals, BH-adjusted Poisson binomial p-value<0.05; **Figure 2C**, **Supplementary Figure 10A**). A striking example of this is *polyomavirus HPyV6,* which was abundantly present (>10%) in many skin sites of some individuals (n=3) despite being lowly abundant in the general population (<0.1%; **Figure 2D**). Interestingly, we also detected co-occurrence patterns in these "blooms" of ancillary species across various body sites (**Supplementary 10B**), including statistically significant coordination between head and limb sites for *polyomavirus HPyV6* (cheek and lower leg, scalp and forearm, Fisher’s exact test p-value<0.05; **Figure 2E**). In some cases, more frequent carriage of an ancillary species across body sites was also associated with host attributes including ethnicity (e.g. Chinese and *Malassezia slooffiae*) and skin phenotypes (e.g. itch severity and *Betapapillomavirus 2;* adjusted p-value<0.05; **Supplementary Figure 10C**). These results highlight the dermotype-specific nature of core and ancillary skin microbial species, their coordinated distribution across body sites within individuals, and their distinct associations with host phenotypes.

### Species co-abundance patterns reveal dermotype-dependent microbial interactions

To understand how dermotypes emerge at various skin sites, we analyzed microbial correlated abundance (co-abundance) patterns with a view to gaining insights into resource competition and metabolic cross-feeding interactions (**Methods**). The scale of our dataset enabled robust identification of >100 species-level co-abundance relationships across the nine body sites and 17 distinct dermotypes (**Supplementary File 7**). These include known antagonism between *C. acnes* and *S. epidermidis*^69–71^, and cross-feeding interactions between *Corynebacterium* and *Staphylococcal* species^72^ (|r|≥0.3, CCREPE q-value<0.05; **Figure 3A**; **Supplementary Figure 11**). Systematic comparison of co-abundance networks and hub species—highly connected taxa— across dermotypes from the same site showed that while a core set of relationships are conserved, 8-48% of co-abundance relationships differed significantly between dermotypes, with highest differences in the axilla and lowest in parietal scalp and forearm (Fisher’s Z-test p-value<0.05; **Supplementary Figure 12A**). For example, *C. acnes* and *M. luteus* abundances displayed a strong negative correlation in the elbow crease dermotype Ac-1 but were positively correlated in Ac-2 (**Figure 3A**), emphasizing the critical context that dermotypes provide for conditional interactions between skin microbial species.

**Figure 3.**
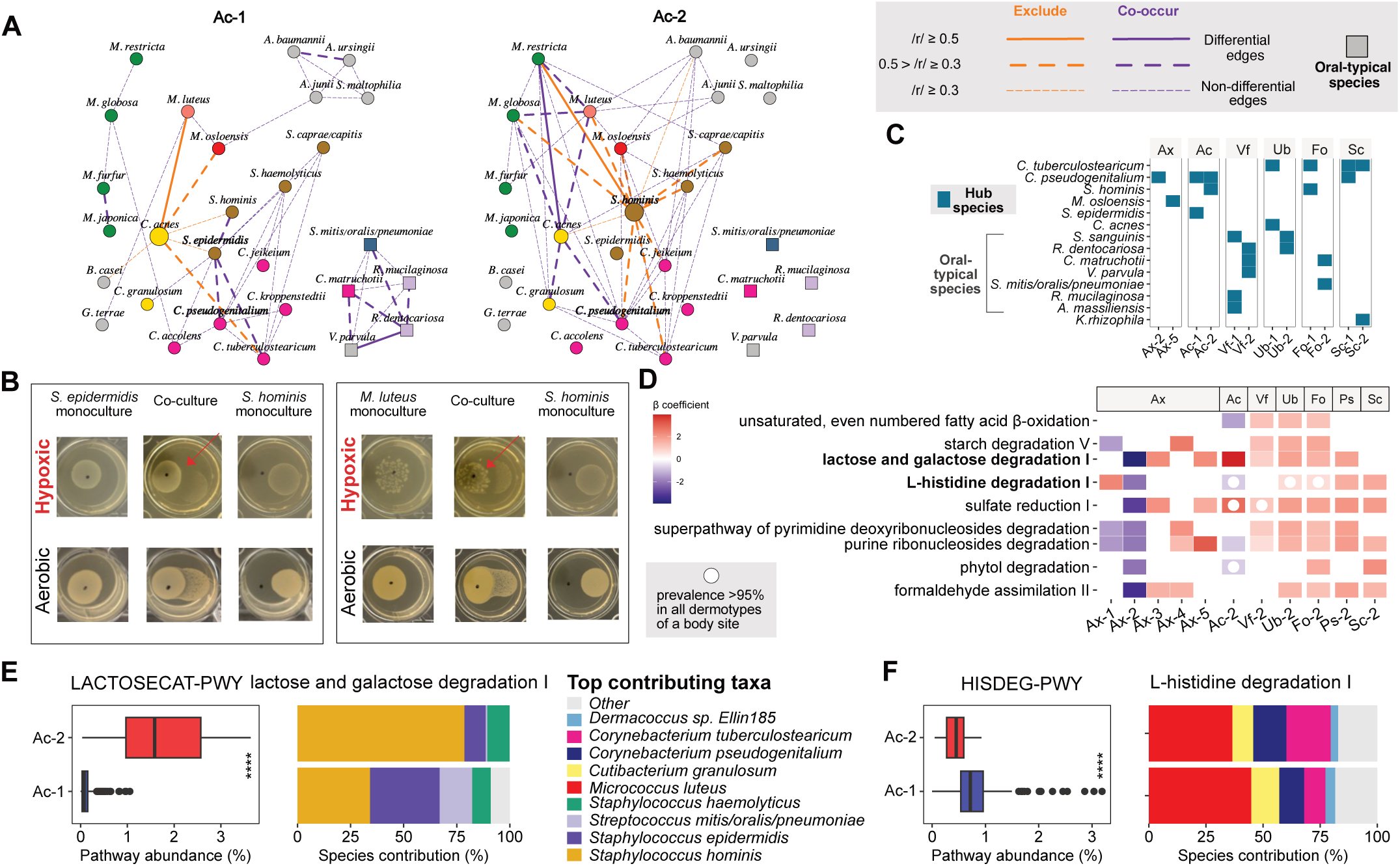
Dermotypes exhibit varied species co-abundance patterns and interactions, and present different metabolic niches. **A.** Species co-abundance networks for dermotypes in the axilla and volar forearm, respectively. Nodes represent microbial species, sized by relative abundance and colored by microbial genera (names are bolded for hub species). Squares denote species typically present in oral cavities. Edges denote significant Spearman correlations (CCREPE |r|≥ 0.3, p-value<0.05): purple for positive and orange for negative correlations. Thin lines indicate shared edges, while thick solid and thick dashed lines denote differential edges with |r|≥0.5 and 0.3≤|r|<0.5, respectively. **B.** Representative agar-based antagonism co-culture assay results showing *S. epidermidis* and *M. luteus* interactions with *S. hominis* under hypoxic and aerobic conditions. **C.** Presence/absence patterns for hub species identified from species co-abundance networks across dermotypes. **D.** Selected degradation-related metabolic pathways that are differentially abundant across dermotypes. **E-F**. Boxplots depicting the variation in community-level relative abundance of, (**E**) Lactose and galactose degradation I pathway, and (**F**) L-histidine degradation I pathway between two elbow crease dermotypes (Ac-1 & Ac-2), with corresponding HUMAnN2-derived species contributions summarized in stacked barcharts. P-values for comparisons across dermotypes were calculated using a one-sided Wilcoxon rank-sum test (∗∗∗∗ denotes p−value<0.0001), and boxplot whiskers represent 1.5× interquartile range or the maximum/minimum data point.

This unexpected pattern extended to core species (significant changes in 10-68% of co-abundance relationships; **Supplementary Figure 12B**), with *C. acnes* in particular frequently exhibiting switched relationships across dermotypes (**Supplementary Figure 12C**). As *C. acnes* is an aerotolerant anaerobe, we hypothesized that oxygen availability strongly impacts *C. acnes* growth and metabolism, and through its interactions with other skin microbes plays a critical role in shaping dermotype structure across body sites. To explore this, we systematically conducted pairwise antagonism assays among a subset of core skin bacteria (*C. acnes*, *S. hominis*, *S. epidermidis* and *M. luteus*) under hypoxic, aerobic and mixed conditions (n=186 experiments, with triplicates; **Supplementary File 8**; **Methods**). Consistent with our co-abundance network analysis, we observed that interspecies inhibitions were often condition-specific (42% of interactions tested). They also revealed several interactions that may have *in vivo* relevance, including partial inhibition of *M. luteus* by *S. epidermidis* and *S. hominis* under aerobic conditions (that mirrored corresponding exclusion edges in *C. acnes*-depleted dermotypes, e.g. Ax-2), and *C. acnes* inhibition by *M. luteus* under mixed (but not aerobic) conditions corresponding to weaker negative correlation edges in *C. acnes*-depleted dermotypes (e.g. Ub-2; **Supplementary File 7-8**). Of particular note, *S. hominis* (the dominant species in the Ac-2 dermotype) was strongly inhibited by *S. epidermidis* and *M. luteus* under hypoxic but not aerobic conditions (**Figure 3B**, **Supplementary Figure 12D-E**). Further testing showed that the agar became alkaline (pH>7) in aerobic conditions but turned acidic under hypoxic conditions (**Supplementary File 8**), indicating that these hypoxia-dependent inhibition patterns could be mediated by acidification or fermentation-derived short-chain fatty acids^74^ that are unfavorable for *S. hominis* growth. Our findings suggest that the common elbow crease dermotype (Ac-1) may represent a hypoxic niche where *C. acnes* flourishes, but *S. hominis* remains suppressed due to inhibition by other skin species. In contrast, the less common elbow crease dermotype (Ac-2) may provide a more aerobic environment enabling *S. hominis* to evade inhibition and dominate, especially with reduced growth of *C. acnes*. Consistent with this hypothesis, we noted that while *S. epidermidis* is a prevalent hub species in the Ac-1 dermotype, it is no longer so in the Ac-2 dermotype and is instead replaced by *S. hominis* as a hub (**Figure 3C**), supporting the notion that *S. hominis* is more well-adapted to the environmental niche of the Ac-2 dermotype.

Further analysis of hub nodes identified key species such as *C. pseudogenitalium* and *C. tuberculostearicum* in 4 out of 12 dermotype-specific networks, highlighting that even low-abundance community members can play a major role in determining community structure (**Figure 3C**). Lastly, we noted that several bacterial species typically found as part of commensal oral flora were network hubs for specific dermotypes in the forearm, upper back, and forehead (**Figure 3C**). Further investigation of their species correlations revealed a guild of co-abundant oral bacteria such as *Veillonella parvula*, *Rothia dentocariosa*, and *Corynebacterium matruchotii*, that were seen in several dermotypes and were strongly co-abundant in some (e.g. Ac-1 & Vf-2; **Figure 3A**, **Supplementary Figure 11**). These species are known to form a "hedgehog" structure in the supragingival biofilm, which regulates pH in response to changes in nutrient availability associated with food intake^75,76^. While the co-abundance patterns in these dermotypes are reminiscent of oral interactions, it remains unknown whether they perform similar pH-stabilizing functions, act as transient co-colonizers, or potentially exacerbate pathogenicity on skin— analogous to the role of oral microbes in gastrointestinal diseases when translocating to the gut^77,78^. Overall, the analysis of species co-abundance networks, supported by the scale of our datasets and experimental data from pairwise antagonism assays, revealed dermotype-specific inter-species interactions and switches in hub species that may be a function of the aerobicity of the environment and likely play an essential role in dermotype assembly.

### Metabolic variations between dermotypes reflect distinct host microenvironments and influence on skin phenotypes

Based on the observation that microbes residing on skin adapt to limited nutrient availability in the form of carbohydrates, lipids, and proteins present in sweat, sebum, and the stratum corneum^2,3^, we hypothesized that differences in metabolic functions between dermotypes of the same body site could provide insights into how they assemble and interact with the host. Systematic comparisons of metabolic pathways present in skin metagenomes (**Supplementary File 9**) highlighted both significant differences in pathway diversity across dermotypes (**Supplementary Figure 13A; Supplementary File 10**), as well as a large fraction of pathways (>35%) being differentially abundant in at least one dermotype of a body site, particularly relating to lipid, carbohydrate, amino acid, nucleotide, and energy metabolism (**Supplementary Figure 13B**). We further observed a suite of metabolic pathways that, while consistently present across distinct dermotypes of a given site, varied significantly in abundance. For example, enrichments in arginine, tryptophan, and branched-chain amino acid biosynthesis pathways (e.g., L-isoleucine, L-valine; **Supplementary Figure 14**), that suggest frequent microbial compensation for amino acid depletion across dermotypes^79–81^.

Focusing on key differences in microbial functions for sugar degradation, amino acid utilization, and fatty acid oxidation, highlighted the strong enrichment of lactose and galactose degradation pathways in the elbow crease dermotype Ac-2 (**Figure 3D**). Galactose is a common component of human sweat^82^ and complex stratum corneum glycoconjugates where it is released by corneodesmosome degradation^83^, and absorbed by skin commensals via diverse phosphotransferase systems^84^. We therefore hypothesized that inter-individual differences in metabolite availability may play an important role in dermotype structure. Indeed, our data indicates that specific *Staphylococcus* species, particularly *S. hominis*, contribute strongly to the enrichment of this pathway (LACTOSECAT-PWY, **Figure 3E**), potentially due to a competitive advantage under these conditions. To test this hypothesis, we compared growth kinetics of *S. hominis* and *S. epidermidis* in mono- and co-culture, under baseline (BHI) and galactose-supplemented conditions (**Supplementary Figure 15A**). While galactose enhanced overall growth in both species, *S. hominis* exhibited disproportionally greater fitness gains in co-culture (>100% relative to *S. epidermidis*; **Supplementary Figure 15B**), suggesting that preferential galactose metabolism by *S. hominis* can provide it a competitive edge and that host-derived metabolites could shape dermotype structure.

Similarly, histidine, a product of epidermal filaggrin breakdown and a common amino acid detected on human skin^85^, can be further degraded by skin microbes^86^. We observed significant variations in the L-histidine degradation pathway across dermotypes (**Supplementary Figure 14**). For example, the *Cutibacterium-*dominant dermotype Ac-1 in the elbow crease exhibited enrichment in the L-histidine degradation pathway, with *M. luteus* being a key contributor (HISDEG-PWY; **Figure 3F**; **Supplementary Figure 16A**). Further analysis of metagenomic and metabolomic data collected at the elbow crease from a separate cohort (n=100; **Methods**), showed that among 20 amino acids surveyed, only histidine derivative urocanic acid (UCA) differed significantly between Ac dermotypes, with Ac-1 subjects enriched for cis-UCA and depleted for trans-UCA (**Supplementary Figure 16B**). This is consistent with the enrichment of *M. luteus* in Ac-1, and in the key enzyme for trans-UCA degradation (urocanate hydrolase; **Supplementary Figure 16A**). While *M. luteus* abundance on skin exhibited a marginal correlation with the levels of histidine and its derivates (ρ=0.21; **Supplementary Figure 16C**), a notable growth advantage was observed *in vitro* with L-histidine supplementation (20% increase in OD; **Supplementary File 12**). Intriguingly, we also observed a negative correlation between lactose metabolism and histidine catabolism pathways specific to Ac-2 subjects, potentially reflecting niche-specific usage of these energy sources (**Supplementary Figure 16D**). Furthermore, the enrichment of the histidine degradation pathway was negatively associated with skin sensitivity (p-value<0.05; **Supplementary Figure 16E**; **Supplementary File 11**), consistent with the known immunosuppressive effects of cis-urocanic acid^87^. Other microbiome functions associated with host phenotypes included depletion of isoleucine biosynthesis pathway (a source of natural moisturizing factor^88,89^) in the volar forearm in subjects with dry skin, as well as an age-associated enrichment in the phosphopantothenate biosynthesis I pathway in the cheek (a precursor for coenzyme A and essential for lipid metabolism^90^), that may compensate for reduced sebum production with age^91^. Together, these findings highlight that dermotypes, beyond representing mere taxonomic differences, are niches with distinct metabolic properties that can be shaped by differences in host-derived nutrients, and in turn can impact host skin conditions.

### Dermotype distribution is associated with host factors and skin discomfort/diseases

Skin microbial composition can be shaped by a combination of endogenous and exogenous host factors (e.g., genetics, skin physiology, lifestyle, and skin conditions), and can in turn influence some of them. We leveraged the scale of our cohort to evaluate associations between dermotypes and host demographics, skin physiological measurements, >15 personal care behaviors, and >25 skin conditions (**Methods**). Focusing on attributes relevant to each body site, we identified >35 significant associations (**Supplementary File 13)**.

Among demographic factors, ethnicity exhibited strong associations with axilla and elbow crease dermotypes. In particular, axilla dermotypes dominated by *C. acnes* (Ax-2), *S. hominis* (Ax-3), and *C. kroppenstedii* (Ax-4) were significantly enriched in ethnic Malay, Chinese, and Indian subjects, respectively (adjusted p-value<0.05; **Figure 4A**). In contrast, the *S. hominis*-enriched elbow crease dermotype (Ac-2) was more frequent in Malay and less common in Chinese subjects (adjusted p-value<0.05; **Figure 4A**). Beyond ethnicity, the *S. hominis*-enriched elbow crease dermotype (Ac-2) and the *Cutibacterium*-depleted upper back dermotype (Ub-2) were significantly associated with female and older subjects (adjusted p-value<0.05). Strikingly, no significant demographic associations were observed for head sites (forehead, scalp, parietal scalp; **Supplementary File 13**), suggesting that other factors such as localized skincare practices or environmental exposures may drive dermotype distributions in these sites.

**Figure 4.**
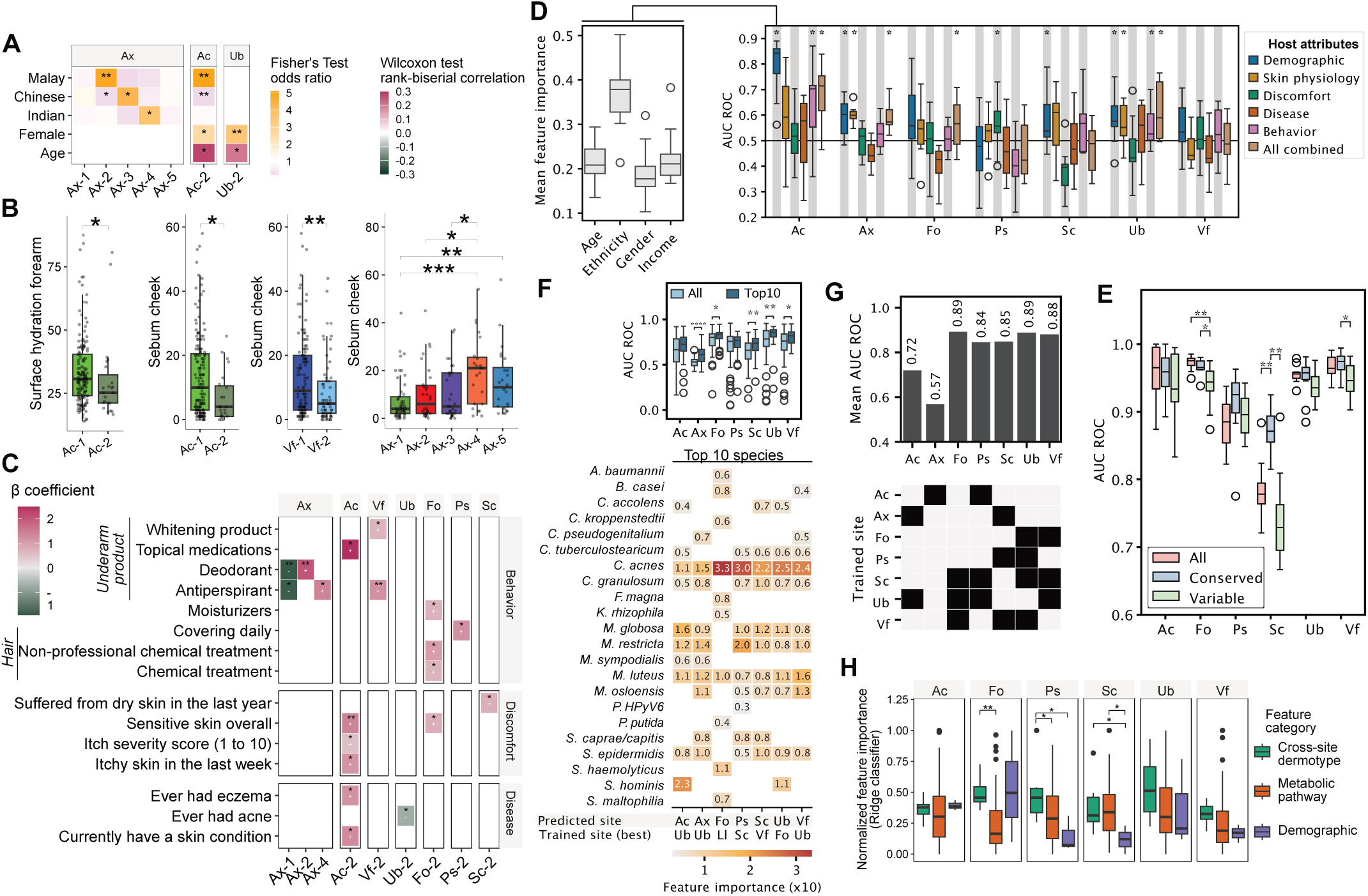
Comprehensive investigation of host attributes, same-site and cross-site intrinsic-microbiome factors for their ability to predict dermotype labels. **A.** Heatmap depicting associations between dermotypes and host demographic factors. Effect sizes are represented as odds ratios (gender, ethnicity; Fisher’s exact test) or rank-biserial correlations (age; Wilcoxon test). **B.** Boxplots displaying variations in skin physiological measurements across dermotypes. **C.** Heatmap of beta coefficients from one-versus-rest logistic regression models, with corresponding p-value labeled on each tile, showing associations between dermotypes and selected personal care behaviors and skin conditions. **D.** AUC-ROC values for dermotype classifiers using random forest models trained on multi-domain host attributes. The left panel compares feature importance of various demographic factors for the Ac dermotype classifier. **E.** Boxplots depicting AUC-ROC values for ridge classifiers trained on different categories of metabolic pathways. **F.** Top boxplots comparing AUC-ROC values for cross-site random forest classifiers trained on the full taxonomic profile versus top 10 abundant species from a single site, with the bottom heatmap showing feature importance for the top 10 species. **G.** Top barplot showing AUC-ROC values for the best-performing cross-site dermotype classifiers trained on dermotype labels from other skin sites, with the heatmap displaying combinations of dermotype labels used for training. **H.** Feature importance comparison in an inclusive ridge classifier trained on host demographic features, cross-site dermotype labels, and conserved metabolic pathway profiles. For inter-group comparisons of continuous variables (e.g., skin physiology, AUC-ROC, feature importance), p-values were calculated using a Wilcoxon rank-sum test, and boxplot whiskers represent 1.5× interquartile range or the maximum/minimum data point. For all subfigures ∗, ∗∗, ∗∗∗ and ∗∗∗∗ denote p−value<0.05, 0.01, 0.001 and 0.0001, respectively.

We next assessed if skin physiological measurements revealed distinct trends across dermotypes. Of note, individuals with an *S. hominis*-enriched elbow crease dermotype (Ac-2) exhibited lower forearm surface hydration (p-value<0.05; **Figure 4B**), indicating that Ac-2 may arise from compromised skin barrier function. Additionally, elevated cheek sebum levels (as a proxy for overall sebum production in the body) were observed in individuals with *C. acnes*-enriched dermotypes at the elbow crease and the volar forearm (i.e., Ac-1, Vf-1), consistent with the robust growth of *C. acnes* in sebum-rich environments. In contrast, higher cheek sebum levels were associated with the *C. kroppenstedii*-enriched dermotype in the axilla (i.e., Ax-4; p-value<0.05; **Figure 4B**). While both species are lipophilic^92^, this divergence likely reflects differences in local lipid composition and species-specific substrate preferences. More broadly, to explore whether site-specific skin physiological measurements reflect body-wide intrinsic states, we conducted further site-agnostic analyses. Intriguingly, we observed strong associations between lower trans-epidermal water loss (TEWL) levels on the cheek and *C. acnes*-depleted dermotypes in the upper back (Ub-2) and forehead (Fo-2; p-value<0.05; **Supplementary Figure 17A**). Similarly, lower TEWL on the forearm associated with the *C. acnes*-depleted parietal scalp dermotype (Ps-2; p-value<0.05; **Supplementary Figure 17A**), highlighting the systemic connections between skin barrier functionality and skin microbial community structure.

Leveraging the availability of extensive skincare routine information, we next assessed their potential impact on dermotypes after accounting for relevant demographic factors (**Methods**). Intriguingly, routine moisturizer use and hair treatments correlated with the *C. acnes*-depleted forehead dermotype (Fo-2), while daily hair covering was associated with the *C. acnes*-depleted parietal scalp dermotype (Ps-2). Similarly, routine deodorant use was positively associated with the axilla dermotype Ax-2 (>55% prevalence, compared to baseline of <20%), while antiperspirant use was linked to Ax-4 (>25% prevalence, compared to baseline of <15%); conversely, both product uses were negatively correlated with the most common axilla dermotype Ax-1, where deodorant and antiperspirant use was notably lower (11% and 5%, respectively; p-value<0.05; **Figure 4C**). Consistent with the antimicrobial effects of these products, Ax-2 and Ax-4 exhibited reduced alpha diversity compared to Ax-1 (**Supplementary Figure 7A**). No significant associations were identified for the last time of washing or antibiotic usage in the last month for any of the dermotypes across the seven skin sites (**Supplementary File 13**). These observations support the notion that while specific skincare practices may partially contribute to dermotype prevalence in the population for some body sites, they do not define dermotype distribution across individuals.

Our data also provides multiple lines of evidence that suggest that dermotype states may serve as biomarkers or influence risk and severity for skin discomfort and diseases (after accounting for demographic factors; **Methods**). For example, individuals with the *Cutibacterium*-depleted scalp dermotype (Sc-2) were significantly more likely to have suffered from dry skin in the last year (>70% enriched, p-value<0.05; **Figure 4C**). Similarly, individuals with the less common upper back dermotype (Ub-2) were less likely to have had acne (>40% depleted, p-value<0.05), while Fo-2 individuals were more likely to have sensitive skin (>60% enriched, p-value<0.05). Of note, individuals with a *Staphylococcus*-dominant elbow crease dermotype (Ac-2) were significantly associated with skin sensitivity, itch severity, recent itchy skin (in the last week), as well as eczema history. The association with skin sensitivity and itch was found to be consistent in Ac-2 subjects even among individuals without eczema history (**Supplementary Figure 17B-C**). To explore if there could be a causal link between dermotype state and risk/severity of itch, we treated human keratinocytes with supernatants of various skin derived bacterial isolates (**Methods**) and assessed pro-IL-33 levels as an important cytokine signal in itch^93,94^. Strikingly, several commensal strains of *S. hominis* (5/8) and *S. epidermidis* (9/12) elicited a significantly elevated pro-IL-33 response compared to controls, while others such as *M. luteus* did not trigger a similar response (**Supplementary Figure 17D**). These observations support the hypothesis that the Ac-2 dermotype may predispose individuals to heightened skin irritation, itch, and associated diseases through the enrichment of specific *Staphylococcus* species and strains.

Finally, we applied machine learning models to test whether the information in host attributes such as demographic data, skin physiological measurements, and questionnaire responses could predict dermotype labels across body sites (**Supplementary Figure 18A**; **Methods**). Machine learning models that were significantly better than random were obtained in all sites except for scalp sites, with the highest classification performance observed in the elbow crease (Ac random forest AUC-ROC=0.8; **Figure 4D**). Among various host attributes, demographic factors emerged as the most predictive in four out of seven sites analyzed (axilla, scalp, upper back, elbow crease), with ethnicity frequently being among the most important features (e.g. for elbow crease; **Figure 4D**, **Supplementary Figure 18B**). In summary, our findings underscore the intricate interplay between host demographics (particularly ethnicity), skin physiology, skincare practices, and skin dermotypes, while highlighting the utility of dermotypes as biomarkers for skin health and disease risk.

### Microbial metabolic function and cross-site dermotypes labels are more strongly predictive of dermotypes

Having shown that certain host attributes can predict dermotype occurrence, we wondered if microbial metabolic functions could serve this purpose. To ensure predictions were not driven by species-specific pathway signatures, we excluded pathways highly correlated with taxa abundances (Spearman ρ≥0.9, p-value<0.05) and trained multiple classifiers (e.g. ridge regression, random forests etc.) to explore key metabolic functions that could jointly predict dermotype structure (**Methods**). Strikingly, very high predictive performance (AUC-ROC>0.95, ridge regression) was obtained with microbial metabolic features across most body sites (**Figure 4E**). Restricting this analysis to conserved pathways (prevalence≥95% across all dermotypes; **Supplementary Figure 19A**), still retained strong predictive ability (**Figure 4E**). A small number of highly informative pathway features were identified (**Supplementary Figure 19B**), and their aggregation into metabolic superclasses highlighted the role of carboxylate degradation, amino-acid degradation, carbohydrate biosynthesis, fermentation, and secondary metabolite biosynthesis in being predictive of dermotype labels (**Supplementary Figure 19C**). It is noteworthy that a small number of microbial metabolic functions were found to be more predictive of dermotype identity than all the host attributes analyzed, emphasizing the role of microbial function and ecology in determining dermotypes.

As dermotypes were strongly correlated across body sites (**Figure 1F**), we further investigated if dermotype labels could be predicted based on taxonomic profiles from other sites. Remarkably, high predictive performance was obtained in many sites (AUC-ROC>0.8, random forests; **Supplementary Figure 20A**), even when only data from a single site (**Supplementary Figure 20B-C)**, or a small number of taxa were used (best AUC-ROC=0.89; **Supplementary Figure 20D**). Intriguingly, data from the lower leg (Ll) consistently ranked highest in feature importance among top-performing classifiers, despite no robust dermotypes being discovered at this site. Several species including *C. acnes*, *S. epidermidis, M. restricta*, *M. globosa,* and *M. luteus,* consistently ranked high in terms of feature importance for dermotype classification across sites (**Figure 4F**), reflecting their body-wide importance for defining dermotype structure. Restricting to just dermotype labels, we again noted that cross-site labels were strongly predictive, and in many cases more so than host attributes (AUC-ROC>0.84, except for Ax, Ac; **Figure 4G**).

Ridge classifiers that integrated cross-site dermotype labels, metabolic pathway profiles, and demographic factors were used to evaluate their relative information content for dermotype prediction (**Figure 4H**), obtaining uniformly high performance across all sites (AUC-ROC>0.9). Notably, cross-site dermotype labels ranked highest by feature importance at the parietal scalp (Ps), upper back (Ub), and volar forearm (Vf; **Figure 4H**), highlighting the importance of systemic microbial assembly patterns underlying dermotype structure across body sites. Although metabolic pathways were generally secondary to cross-site dermotype labels, they provided complementary information and even ranked highest in terms of feature importance for the scalp (Sc), suggesting that microbial metabolic capabilities may have greater influence on dermotype assembly for some sites. Similarly, while demographic factors had lesser feature importance in general, they had significant predictive value in select sites (e.g., elbow crease [Ac] and forehead [Fo]; **Figure 4H**). Overall, our findings emphasize the unexpected strength and predictive power of dermotype coordination across body sites, and the critical role of specific microbial metabolic functions and host demographic factors in shaping dermotype structure and function on skin.

## Discussion

Despite growing efforts to understand the role of the skin microbiome in skin health, population-based investigations defining baseline variations and features have been limited to a few early studies with small cohort sizes^27,29^ (n<20). Here we present the largest-to-date metagenomic characterization of skin microbiomes from multiple body sites (n=18) in a cohort study that (i) is an order of magnitude larger (n=200), (ii) captures broad Asian ethnic representations of Chinese, Malay, and Indian individuals, and (iii) was conducted in a geographically localized setting with a tropical climate (consistent humidity levels and temperature year-round), minimizing confounding effects from geography and seasonable variability^95–97^. This provides a well-powered, rich dataset with consistent protocols and comprehensive mapping of skin microbiomes (n=3,572 metagenomes), that is complemented by detailed phenotypic information, creating a valuable resource for studying skin microbial ecology and function, and its relationship with host attributes and skin health.

Leveraging this dataset, we uncovered substantial population heterogeneity in the skin microbiome by comprehensively mapping robust skin microbiome dermotypes in multiple body sites (7 out of 9; **Figure 1D-E**), thus establishing dermotypes as an important paradigm for population stratification in skin health studies. Our work transcends prior studies that provided limited evidence for, (i) context-specific community types (e.g., eczema-associated elbow crease^49^, air pollution-exposed face^51^), or (ii) site-agnostic patterns that overlook key site-specific differences^50^. Critically, our data supports the idea that the dermotypes identified here represent distinct microbiome states in regions with higher density of support (**Figure 1D**, **Supplementary Figure 6**), unlike the continuum described in some gut enterotype studies^98,99^. Furthermore, these dermotypes exhibit differential abundances for many species rather than a single signature species (**Supplementary File 6**), and yet near-perfect stratification is possible with a few marker species, enabling cost-effective strategies for clinical translation (e.g. using qPCR panels). This opens up many avenues for studies based on dermotype stratification. For example, the axilla demonstrates striking heterogeneity that has never been described before, with evidence for at least five robust dermotypes dominated by multiple *Staphylococcus* and *Corynebacterium* species. While these show some associations with ethnicity, sebum levels, and underarm product usage^100^, they are not defined by them, and further studies are needed to understand how these dermotypes emerge and influence host health (e.g. body odor^22^). Similarly, the *Staphylococcus*-dominant elbow crease dermotype (Ac-2) identified here supersedes our previous results specific to atopic dermatitis patients^49^, defining a group of individuals that may be at a greater risk for skin sensitivity, itch, and eczema, and could benefit more from microbially-targeted interventions. Our additional observations of *M. luteus*-enriched dermotypes in dry sites, and *M. restricta*-enriched dermotypes in sebaceous scalp regions, provide the basis for further investigation into dermotype function in skin conditions such as dryness and dandruff, potentially guiding the development of new pre-, pro- and postbiotic treatment strategies for these large consumer care market segments.

Extending beyond the long-standing view of site specificity in skin microbiomes^27,28^, our large multi-site study revealed remarkable cross-site correlations in dermotype patterns, even between physically and physiologically distinct skin sites (e.g. dry forearm and sebaceous forehead sites; **Figure 1F**). This coordination is further evident from the ability to classify dermotypes at one site using information from as few as 10 species or dermotype labels from another site (**Figure 4F**), a striking observation that has not been reported before in human microbiome studies^48,65,101^. This cross-body site correlation of dermotypes, combined with the high bilateral concordance in skin microbiomes across all 9 sites and most individuals as we establish here (**Figure 1B**, **Supplementary Figure 2**), indicates a deeper systemic coordination of microbiome states across the body within an individual. The observation that loss of bilateral concordance can also be correlated across body sites further supports the idea that dermotype assembly is shaped by coordinated influences across the body (**Supplementary Figure 3C**). There are several potential explanations for dermotype coordination, including strain sharing, host factors, and microbial functions. For example, correlated abundances of a few core species shared across multiple sites (e.g., *C. acnes*, *S. epidermidis*, and *M. restricta*; **Figure 2B**, **Supplementary Figure 9**), perhaps due to strain-specific ecological functions^102–104^, may partially explain our observations. However, the preference for specific species/strains alone does not fully explain the patterns observed, as some taxa show no significant correlations between specific sites (e.g., *Staphylococcus* species in axilla and elbow crease; **Supplementary Figure 9**). Similarly, while shared host demographic, physiological or behavioral factors may partially explain dermotype assembly and thus coordination (**Figure 4A-C**), our predictive models suggest that they may not be the dominant explanatory factors, and that key microbial metabolic functions likely play a greater role in many sites (**Figure 4H)**. Taken together, our findings establish the coordination of dermotypes across body sites as an important phenomenon that needs to be considered in any model for dermotype assembly, and the potential to predict dermotypes across body sites based on information for a few body sites.

To further understand the role of microbial functions in dermotype assembly, we systematically investigated interspecies interactions based on mining of dermotype-dependent co-abundance patterns in our large datasets, and through pairwise antagonism assays between core species. These experiments identified varying oxygen availability (shaped by skin physiology^105,106^) as likely a key factor impacting interspecies interactions and dermotype assembly (e.g. hypoxia-dependent *S. hominis* inhibition by *S. epidermidis* and *M. luteus*; **Figure 3B**). Such oxygen-dependent antagonism may arise from differential toxin production (e.g. *S. epidermidis* phenol-soluble modulins produced under hypoxia^107^) or redox-sensitive metabolic cross-feeding^108^. Similar modulation was also seen in a recent report of multi-species interaction dynamics in healthy and dandruff scalp microbiomes^109^. In addition to oxygen gradients, nutrient availability differences (e.g. lipids, carbohydrates and amino acids) may also explain the large number of dermotype-specific co-abundance patterns seen in our data (**Supplementary Figure 12A-B**). For example, in sebaceous environments *C. acnes* may grow well by hydrolyzing sebum triglycerides into free fatty acids and thus acidifying the skin surface. This acidification, combined with lipid reduction and potential antimicrobial activities, likely contributes to the unfavorable conditions seen for *M. luteus* and other skin bacteria in our data^110^. Conversely, under lipid-depleted conditions, the competitive dominance of *C. acnes* may diminish (**Supplementary File 8**), while species such as *S. hominis* and *M. luteus* may leverage that availability of other nutrients, such as galactose and histidine as shown here (**Figure 3E-F**), to flourish and define dermotype structure based on conditional interactions. Overall, the prominence of shared microbial pathways such as amino acid, carbohydrate, and nucleotide biosynthesis in our predictive models (**Supplementary Figure 19**) emphasizes the role of metabolic function rather than taxonomic identity in driving niche specialization, dermotype-specific microbial interactions and ecosystem assembly.

The combination of extensive host demographic, skin physiological, behavioral, and discomfort/disease data, together with skin microbiome characterization across multiple body sites allowed us to not only investigate the many dimensions through which specific host attributes may influence dermotype prevalence, but also the potential for dermotypes to impact host phenotypes and thus serve as functional biomarkers for skin health and disease. In particular, several of our observations in the *S. hominis*-dominated elbow crease dermotype (Ac-2) including associations with older age, reduced surface hydration, skin sensitivity, and itch, demonstrate a bidirectional interplay between microbial ecology, metabolic activities, and barrier dysfunction (**Supplementary Figure 21**). Compromised skin barrier, whether due to dryness, inflammation, or mechanical damage from scratching, may enhance oxygen penetration, creating aerobic microenvironments where *S. hominis* escapes inhibition by *S. epidermidis* and *M. luteus*. Notably, *S. hominis* may further circumvent *S. epidermidis* antagonism via galactose metabolism, which disproportionately enhances its growth relative to *S. epidermidis* (**Supplementary Figure 15**). Thriving under these niche conditions in the skin, *S. hominis* can exacerbate itch via IL-33 induction in keratinocytes, perpetuating an itch-scratch cycle that further compromises barrier integrity and reinforces the aerobic conditions that favor its dominance (**Supplementary Figure 17**). In contrast, enhanced *M. luteus*-mediated histidine degradation in Ac-1 dermotypes can suppress inflammation and contribute to reduced skin sensitivity in Ac-1 individuals through production of immunosuppressive cis-urocanic acid. Such metabolic cross-talk between the host and microbes, alongside those between microbes, underscores the complex interactions that may underlie dermotype associations with skin conditions (**Supplementary Figure 21**). Future work in 3D skin and animal models could help disentangle these intricate relationships and enhance our understanding of dermotype contributions to various skin discomforts (e.g. skin irritation, itch, malodor etc.) and skin diseases (e.g. eczema, psoriasis, acne etc.). The ability to stratify subjects based on skin microbiome dermotype status could offer a convenient (non-invasive) and cost-effective (based on targeted panels) approach for predicting disease risk and severity, monitoring treatment response, and identifying patients who would respond to specific interventions. Our finding and dermotype prediction models thus redefine the roadmap for future research into therapeutic strategies for skin conditions based on microbiome-informed stratification and microbiome-targeted interventions.

## Methods

### Subject recruitment and microbiome sampling

Subjects were recruited as part of the Asian Skin Microbiome Programme (ASMP) sub-study for the Health For Life In Singapore (HELIOS) study (https://www.healthforlife.sg/), a large population-based cohort that is central to the National Precision Medicine program (NPM) in Singapore. The HELIOS cohort study specifically recruits subjects to ensure that their demographics accurately match that of Singapore’s population, considering factors such as ethnicity, age, gender and socio-economic status. Informed consent was obtained from all subjects, and associated protocols for the study were approved by the NTU Institutional Review Board (IRB reference number: IRB-2016-11-030). Subjects were excluded from the study if they were currently pregnant, breastfeeding, had any ongoing acute illness, had recent major surgery, were under the age of 30, or had any mental incapacities that would prevent them from understanding and giving informed consent. Upon arrival to the HELIOS clinic, subjects were given time to acclimatize to the temperature of the clinic. Following this, subjects were asked to complete a skin health questionnaire. This extensive questionnaire has questions related to several topics including overall skin health, skin-related medical treatments, personal care routines, product usage, lifestyle habits, and finally a series of questions relating to menopause for all female subjects (**Supplementary File 14**). Subjects were then moved to a climate-controlled room (temperature: 20-22°C; relative humidity: 40-60%) for sampling by a trained research coordinator/clinician. To minimize external influences, participants were instructed to avoid application of topical products to sampling and measurement areas for >12 hours prior to the measurement; to avoid washing with tap water, synthetic detergents, and alkaline soaps for >2, >5, and >10 hours, respectively; and to refrain from ingestion of caffeinated beverages for >3 hours before and during the study period.

A diverse set of body sites were chosen for skin surface sampling, based on their relevance for skin health and diseases^53,17,54^, importance for consumer care studies, and to capture the breadth of the human skin microbiome. These consisted of nine different body sites, sampled on both the left and right sides (**Figure 1A**), including: scalp (Sc), forehead (Fo), cheek (Ch), upper back, (Ub), antecubital fossa (elbow crease, Ac), volar forearm (Vf), and front of lower leg (Ll), for which an adhesive tape disc was used for sampling (D-Squame Standard Sampling Discs, D100, CuDerm). The tape was repeatedly placed on the same location with pressure and peeled 25 times to saturate discs with skin surface material. For the remaining two sites, i.e. the parietal scalp (5cm above the ear, Ps) and axilla (armpit, Ax), sterile swabs (FLOQSwabs Flocked Swabs, 552C, Copan) were used to sample the skin microbiome by soaking in 0.9% saline and rubbing back and forth on the skin for 30 seconds. The swab-based method was selected for these sites due to its greater suitability in areas with dense hair, and the established very high concordance of metagenomic profiles obtained relative to tape-based sampling^5^. One tape disc or swab was collected from each side (left and right) of the skin sites, giving 14 tape discs and 4 swabs per subject. All collected samples were stored at −80°C before sample processing. Negative control samples were collected in the form of blank tape discs (n=34) and swabs (n=12) that were processed in the same way as skin samples.

### Skin physiological measurements

All measurements were conducted in a controlled temperature (20-22°C) and humidity (relative humidity 40-60%) environment after at least 20 minutes of acclimatization. Trans-epidermal water loss (TEWL) was measured using a VapoMeter (Delfin) on the forearm (left or right) and on the cheek (just below the left eye). pH was measured using a pH meter (Delfin) on the forearm (left or right). Stratum corneum surface hydration was measured using a MoistureMeter (Delfin) on the forearm (left or right). Skin sebum secretion was measured using a SebumScale (Delfin) on the cheek (just below the left eye). Three consistent consecutive readings were taken for all measurements and averaged.

### DNA extraction

Genomic DNA was extracted from skin tapes and swabs using a previously established protocol^5,49^, with modifications to improve yield. Briefly, samples were transferred into Lysing Matrix E tubes containing 500 µL of Buffer ATL (Qiagen) and subjected to bead-beating at 4 m/s for 30 seconds, repeated twice, using a FastPrep-24 Automated Homogenizer (MP Biomedicals). Following homogenization, tubes were centrifuged at maximum speed for 5 minutes, and 200 µL of supernatant was transferred to 2 mL EZ1 sample tubes (Qiagen). Proteinase K (12 µL) was added, and samples were vortexed and incubated at 56 °C for 15 minutes. DNA was then purified using the EZ1 Advanced XL Instrument (Qiagen) with the EZ1 DNA Tissue Kit and eluted in 50 µL of EB buffer.

### Construction and sequencing of metagenomic libraries

Purified genomic DNA underwent NGS library construction steps using NEBNext Ultra II FS DNA Library Prep Kit according to manufacturer’s instructions. 26 µl of DNA added to fragmentation reagents were mixed well and placed in a thermocycler for 10 mins at 37°C, followed by 30 mins at 65°C to complete enzymatic fragmentation. Post fragmentation, adaptor-ligation was performed using Illumina-compatible adaptors, diluted 10-fold as per kit’s recommendations prior to use. Post-ligation, size selection or cleanup of adaptor-ligated DNA was performed using Ampure XP beads in a 7:10 beads-to-sample volume ratio. Unique barcode indexes were added to each sample and amplified for 12 cycles under recommended kit conditions to achieve multiplexing within a batch of samples. Finally, each library sample was assessed for quality based on fragment size and concentration using the Agilent D1000 ScreenTape system. Samples that passed the quality-checks were adjusted to identical concentrations employing dilution and volume-adjusted pooling. The multiplexed sample pool was paired-end (2×151bp) sequenced on an Illumina HiSeq X Ten system. All libraries were sequenced at the Novogene sequencing facility in the Genome Institute of Singapore following standard Illumina sequencing protocols.

### Taxonomic and functional profiling

Raw Illumina sequencing reads were processed using an in-house Nextflow pipeline (https://github.com/CSB5/shotgunmetagenomics-nf). Quality control and adapter trimming were performed with FastQC v0.20.0 (default parameters). Human reads were removed by mapping to the hg19 human reference genome using BWA-MEM (v0.7.17-r1188, default parameters), and filtering with SAMtools (v1.7). Taxonomic profiling of the remaining reads was conducted with MetaPhlAn2 (v2.7.7, default parameters). During sequencing data generation (2020 January), MetaPhlAn3 (v3.0, 2020 April release) was tested but frequently omitted eukaryotic taxa, including fungal species commonly observed and cultured from human skin; additionally, MetaPhlAn4 (v4.0, 2022 April release) outputs viral profiles separately, without integration into the non-viral taxonomic profile. Due to these limitations, we opted to use MetaPhlAn2 consistently in this long-running project. To address the limited representation of *Malassezia* species in MetaPhlAn2 (a commonly found fungal member of the human skin microbiome^58,111^), we integrated results from PathoScope^112^ (v2.0) for *Malassezia* species. Briefly, after human read removal, we aligned the remaining reads to 14 *Malassezia* genomes (*M. furfur, M. obtusa, M. yamatoensis, M. japonica, M. sympodialis, M. pachydermatis, M. restricta, M. globosa, M. slooffiae, M. caprae, M. cuniculi, M. dermatis, M. equina,* and *M. nana*) downloaded from NCBI using bpipe (v0.9.8.6_rc), and normalized mapped read counts by the total number of reads to estimate the relative abundance of *Malassezia* species. Lastly, MetaPhlAn2 taxonomic profiles (excluding *Malassezia*) were rescaled by multiplying by the proportion of reads that were not mapped to *Malassezia* (calculated from PathoScope results). The rescaled MetaPhlAn2 profile and PathoScope-derived *Malassezia* abundances were merged and renormalized using Total Sum Scaling (TSS) to generate final taxonomic profiles (n=3,586) for downstream analyses. In addition, we reannotated *Enyhydrobacter aerosaccus* as *Moraxella osloensis*, consistent with a previous study showing that 99.4% of sequences labeled as *E. aerosaccus* in the MetaPhlAn2 database were identified as *M. osloensis* using BLASTN against the NCBI nt database^50^.

Sequencing data from negative control samples (n=46; **Supplementary File 1**) and information from existing databases (PubMed, DISBIOME^113^, MicroPhenoDB^114^) were used to identify potential artifacts in the taxonomic profiles, particularly *kitome* species derived from DNA extraction kits and sequencing reagents (**Supplementary Figure 1**). In the first tier, species commonly found in negative controls (median abundance≥0.1%) were evaluated for negative correlation with library concentration, a signature of a contaminant species^115^ (negative Spearman correlation, p-value<0.05). These kitome candidates were then used to identify additional taxa highly correlated with them, potentially also arising from contamination (Spearman ρ≥0.7, p-value<0.05). The second tier refines this initial selection by restricting to genera not previously reported on human skin or associated with skin disease in three different databases (PubMed, DISBIOME^113^, MicroPhenoDB^114^). Poor-quality profiles were removed (n=14, 0.4%; kitome total abundance>50%). Kitome taxa (**Supplementary File 2**) and extremely low-abundance species (relative abundance<0.1%) were excluded from taxonomic profiles (set to 0), and profiles were TSS-renormalized (n=3,572; **Supplementary File 3**).

Functional profiles and detailed species-specific reconstructions of microbial metabolic pathways (MetaCyc) were generated with HUMAnN2 (v2.8.1, default parameters). Pathways that were unannotated, or highly correlated with kitome species (Spearman ρ≥0.7) were filtered out. TSS-renormalized pathway abundance profiles were used for subsequent analyses (n=3,550; **Supplementary File 9**). Downstream data analysis was performed in R v4.4.0 unless stated otherwise.

### Identification of dermotypes

Dermotypes were identified via a three-step protocol **(Supplementary Figure 6)**. Firstly, Partitioning Around Medoid (PAM) clustering was performed using Bray-Curtis dissimilarity for each site. The optimal number of clusters was determined using key indices (prediction strength, silhouette index, and Calinski-Harabasz index). In cases of ambiguity (i.e., disagreement across various indices), we employed Mean-Shift clustering (bandwidths determined by cross-validation) to search the local maxima of kernel density estimated in PCoA space (PCoA 1-3). Next, we assessed sample stability and cluster robustness via bootstrapping (n=100). Clusters were deemed robust if over half of the samples were consistently grouped into the same cluster in over 90% of iterations (median stability≥0.9). Only sites with at least 2 robust clusters detected were defined as having distinct dermotypes for subsequent analyses. Finally, we relied on PAM to infer cluster memberships for robust dermotypes, with samples with <90% stability excluded from further analyses.

### Diversity and differential abundance analysis

Metrics for α-diversity (Shannon index) and β-diversity (Bray-Curtis dissimilarity) were computed using the vegan package (v2.6.4). Permutational multivariate analysis of variance (PERMANOVA) was performed with Bray-Curtis dissimilarity values to quantify differences in microbial compositions using the adonis2 function from vegan^2^ (v2.6.4) with 1,000 permutations and restrictive permutations within blocks of SubjectID. Multivariable analysis estimating which microbiome attributes (species, pathways) differ significantly across comparison groups was carried out using MaAsLin2 (linear model, "LOG" transformation, prevalence≥10%, BH-adjusted p-value<0.05), including demographic covariates (age, gender, ethnicity) as fixed effects and subject ID as a random effect when applicable.

### Construction of species co-abundance networks

Species co-abundance networks were constructed with taxonomic profiles with high cluster membership stability (≥90%) in each dermotype using the "netConstruct" function in NetCoMi^116^ (v1.1.0), and visualized using igraph^117^. Prevalence filtering was applied to focus on prevalent taxa (≥30% for Vf, Ac, Ub, and Fo dermotypes, and ≥10% for Sc, Ps, and Ax dermotypes); this filtering ensures that each network construction included at least 15 taxa, minimizing spurious findings and allowing for biologically meaningful insights. Compositionally corrected Spearman ρ for each species pair and corresponding p-values were obtained from CCREPE^118^ (v1.34.0), and downstream analysis were restricted to species pairs with strong co-abundant or co-exclusion patterns in each dermotype (|ρ|>0.3, BH-adjusted p-value<0.05; **Supplementary Figure 12**). Hubs were defined as species with correlation strength-weighted degree centrality above the 95^th^ percentile in each network. Differential edges between dermotypes for each skin site were identified using Fisher’s z-test^119^ for correlation values.

### Antagonism co-culture assays

The bacterial strains studied (*Cutibacterium acnes* 6919, *Micrococcus luteus* 10240, *Staphylococcus epidermidis* 12228, and *Staphylococcus hominis* 27844) were obtained from ATCC. *C. acnes* was cultured in half-strength BHI broth in an anaerobic chamber at 37°C overnight with agitation, and other species were cultured under the same condition aerobically. After measuring the optical density at 600 nm (OD600), each culture was diluted to OD600=0.1 with fresh broth (5mL total) and incubated at 37°C with agitation (220 rpm) until growing to an OD that corresponds to the mid-log phase of the bacteria (i.e., *C. acnes* OD=0.6; *M. luteus* OD=1; *S. epidermidis* OD=2; *S. hominis* OD=2).

Assay plates were prepared by dispensing 2mL of molten half-strength BHI agar into each well of a 12-well plate and allowing it to solidify and dry thoroughly. Each bacterial pair was assayed in triplicates, with one well per replicate. For hypoxic conditions, plates were placed in compact plastic pouches with anaerobic sachets (Thermo Fisher Scientific Oxoid AnaeroGen, Cat. No. AN0020D). A total of 40μL of test bacterial culture (bacteria being tested for inhibitory activity; 10μL per droplet, sequentially spotted four times) was applied to each well, allowing each droplet to dry before applying the next. The following day, cultures of target bacteria (bacteria to be assessed for inhibition) were diluted 100× in 1× PBS (10× for *C. acnes*). A 10μL aliquot of the diluted target culture was then spotted adjacent to the test bacteria in each well. The plates were incubated at 37°C overnight, or two nights if the target bacteria was *C. acnes*, and clearing zones were assessed the following day.

### Effect of galactose on *Staphylococcus* species growth

The effect of galactose on the growth of *Staphylococcus* species was assessed by performing monoculture and co-culture experiments with *Staphylococcus epidermidis* 12228 (SE) and *Staphylococcus hominis* 27844 (SH) in Brain heart infusion (BHI) broth with and without 5% galactose supplementation. Briefly, six different experimental conditions were designed: 1) BHI + SE 2) BHI + SH 3) BHI + Galactose 5% + SE 4) BHI + Galactose 5% + SH 5) BHI + SE + SH 6) BHI + Galactose 5% + SE + SH, with two separate negative controls including, BHI and BHI + Galactose 5%. Growth curve experiments were performed at 37°C for 48h with high amplitude, fast speed shaking in a 100-well Honeycomb plate (Growth Curves USA) with a working volume of 200µl in each well. All the experiments were performed in quadruplicates. Optical Density (OD) at 600nm measured using an automated Bioscreen C MBR (Growth Curves USA) reader at 30-minute intervals was plotted to obtain growth curves.

Cell pellet was used for DNA extraction for qPCR analysis. Each sample was treated with 500µl Buffer ATL (Qiagen) in Lysing Matrix E tubes and homogenized twice at 4 m/s for 30 seconds using the FastPrep-24 (MP Biomedicals). After centrifugation at maximum speed for 5 minutes, 200µl of supernatant was transferred into 2ml EZ1 tubes (Qiagen), mixed with 12µl Proteinase K, vortexed, and incubated at 56°C for 15 min. DNA was extracted in EZ1 Advanced XL Instrument (Qiagen) using EZ1 DNA Tissue Kit (Qiagen) and eluted in 50µl EB buffer. DNA concentration was determined using a Qubit 4 Fluorometer with the Qubit dsDNA BR Assay Kit (Invitrogen). qPCR reaction mixture was prepared following the instructions in the PowerSYBR Green MasterMix protocol (Applied Biosystems, Cat. No. 4368577) and 1μl of the extracted DNA was used as the DNA template. Reactions were run in triplicates on a 96-well plate in ViiA 7 Real-Time PCR System (Applied Biosystems) with the following cycling conditions: 95°C for 2 min; 40 cycles of 95°C for 15 s, 60°C for 1 min; followed by a dissociation step. Standard curves were generated using gBlocks (IDT) diluted to 10⁸–10³ copies/µl. Primer and gBlock sequences used for the qPCR are listed in **Supplementary File 17**.

### Effect of histidine on *Micrococcus luteus* growth

The effect of histidine on *Micrococcus luteus* was assessed by culturing *M. luteus* in Brain heart infusion (BHI) broth with and without L-histidine supplementation (50µg/ml concentration), with BHI used as a negative control. Growth curve experiments were performed at 37°C for 6 days at high amplitude, normal speed shaking in a 100-well Honeycomb plate (Growth Curves USA) with a working volume of 200µl in each well. All the experiments were performed in triplicates. Optical Density (OD) at 600nm measured using an automated Bioscreen C MBR (Growth Curves USA) reader at 30-minute intervals was plotted to obtain growth curves.

### Metabolomics sample collection and processing

An additional cohort of 100 subjects was recruited and skin samples collected in a separate sub-study of the HELIOS study for skin metagenomic and targeted metabolomic analysis of 20 amino acids (Ac, elbow crease; manuscript in preparation). For metabolomics, D-Squame (D100, Clinical and Derm) skin tapes were collected in the same manner as for metagenomics, except that they were pushed deep into their collection tube and processed by adding 1 ml of extraction buffer per tube, consisting of 0.1% formic acid in water, and 0.1μg/ml heavy isotope labelled standards. The tubes were then water bath sonicated (Elma) for 15 minutes at 4°C. Following sonication, 120 μl from each sample was loaded onto 96 well chimney plates (Greiner Bio-One) prior to acquisition by liquid chromatography mass spectrometry (LCMS). An additional 150μl was used to determine the protein concentration with the Micro BCA kit (Thermo Scientific). 96 well plates holding the samples were loaded into the Vanquish LC system (Thermo Scientific), utilizing the Atlantis Premier BEH C18 AX 1.7 µm, 2.1×100 mm column (Waters) before being acquired on the TSQ Quantis triple quadrupole MS machine (Thermo Scientific). All data files were analyzed via the software Skyline. No heavy isotope labelled standard were available for trans-urocanic acid (trans-UCA), so the cis-UCA heavy isotope labelled standard was used as a proxy. Output values from each sample were then normalized against their respective Micro BCA derived protein concentrations. Quantified values for histidine and its key derivatives (cis- and trans-UCA) are available as supporting data on FigShare (https://doi.org/10.6084/m9.figshare.28850783.v1).

### Dermotype associations with host attributes

Samples with unstable dermotype assignments (stability<90%) were excluded for each skin site; additionally, subjects with highly stable yet discordant dermotypes at the same site were excluded, to enhance consistency. Associations between dermotypes and demographic or physiological data was assessed in two stages. In the first stage, categorical variables were analyzed using Fisher’s exact test, while continuous variables were evaluated using Wilcoxon tests. Only attributes that demonstrated overall statistical significance were carried forward to the second stage. In the second stage, categorical variables were analyzed using Fisher’s exact test in a one-versus-the-rest framework, with odds ratios recorded as effect sizes; for continuous variables, pairwise Wilcoxon tests were performed, and rank-biserial correlations were reported as effect sizes.

Associations between dermotypes and questionnaire responses were assessed using logistic regression models with a one-versus-rest approach to accommodate multi-label cases (e.g., with five axilla dermotypes), with Beta coefficients recorded as effect sizes. When dermotypes exhibited significant associations with demographic factors in prior analyses (**Figure 4A**), these factors were included as covariates in the regression model. We prioritized responses relevant to the body site (e.g., underarm product usage for axilla dermotypes), and included response groups with at least five observations to maintain statistical power. Questions related to specific product name were excluded.

### Machine learning-based dermotype classification

To ensure comparability across questionnaire responses, all continuous variables (e.g., itch severity) were z-standardized, and categorical variables were one-hot encoded. Binary-class machine learning classifiers were trained to predict the dermotype of the sample based on its own metagenomic profile. A one-vs-rest strategy was employed to accommodate multi-label classification (i.e., axilla dermotype classification). To address class imbalance, we employed the Synthetic Minority Over-sampling Technique (SMOTE) to up-sample minority class instances^120^. Samples with unstable clustering membership (stability<90%) were excluded from each skin site.

We selected six machine learning methods based on their distinct properties, such as linear and non-linear capabilities. These included decision trees, logistic regression, ridge regression, random forests, support vector classifiers^121^ (linear and gaussian kernel) and XGBoost^122^. Model training and performance reporting utilized a nested cross-validation approach to provide an unbiased performance estimate by accounting for hyperparameter selection—this involved 4-fold training with 3-fold inner cross-validation for hyperparameter tuning. Tuned models were evaluated on a withheld 1-fold test, and the process was repeated five times with different data splits. Mean accuracy and AUC-ROC scores were reported (one-vs-rest, macro average). Using the same framework as described above, we trained additional dermotype classifiers with alternative input features, including: 1) the sample’s metabolic pathway profile, 2) metagenomic profiles from other skin sites within the same host. Further analyses were based on the best-performing methods: random forests for models using metagenomic profiles, and ridge regression for models using metabolic pathway profiles.

Random forest classifiers were further trained to assess associations between host factors and dermotypes using host attributes (i.e., demographics, skin physiological parameters, and questionnaire data), as they provided the best performance. Only subjects with concordant dermotype labels between bilateral sites were included. Lastly, to determine the relative informativeness of different feature sets for dermotype classification at a given skin site, we trained an inclusive ridge classifier using three broad categories of features: demographic factors, conversed metabolic pathway profiles, and dermotype labels from other skin sites. Categorical values were one-hot encoded and continuous values z-standardized.

### Commensal *Staphylococcus* strain isolation

Commensal *Staphylococcus* strains were isolated from skin swabs for 4 healthy donors across 4 separate body sites: antecubital fossa, axilla, cheek and scalp. Swabs were immersed in 2.5mL BHI broth (Oxoid), incubated at 37°C at 210rpm for 2 hours before being spread onto TSA-SB (Thermo Fisher Scientific) and Baird Parker (Oxoid) plates and incubated at 37°C for 24 hours. Colonies were then picked and grown in 4ml BHI broth for 16 hours overnight at 230rpm and 37°C. 200μL of the cultures were spun down at 5000rpm for 5 minutes. To extract gDNA the pellet was resuspended in 50μL QuickExtract (Lucigen). Bacterial suspensions were then heated to 65°C for 6 minutes, briefly vortexed, then heated to 98°C for 4 minutes and briefly vortexed again. PCR reactions were setup with 10μL KAPA SYBR FAST (Merck), 6.4μL nuclease free water (Promega), 3μL extracted gDNA and 0.6μL primer mix. Primer mixes were specific for 4 specific *Staphylococcal* species (**Supplementary File 15**). PCR was run on an AriaMx Real Time PCR System (Agilent Technologies). PCR products were sent for Sanger sequencing to confirm product sequences. The identified *Staphylococcal* colonies were inoculated into 4ml BHI broth and grown overnight for 16 hours at 230rpm and 37°C. Optical density (OD) was measured with a SpectraMax M5 Microplate Reader (Molecular Devices). Overnight cultures were then grown from OD 0.1 in fresh BHI for 26 hours at 230rpm and 37°C. Final OD measurements for all strains after 26 hours was normalized to OD 2.75. Bacterial cultures were spun down at 5000rpm for 5 minutes, pellets were discarded and supernatants stored at 4°C.

### Human keratinocyte cell culture and Nano-Glo HiBiT

A genetically modified N/TERT keratinocyte cell-line containing a 33 nucleotide HiBiT tag directly adjacent to the IL33 gene, was utilized for Nano-Glo HiBiT experiments. Keratinocytes were cultured in keratinocyte serum free medium (KSFM; Gibco) supplemented with bovine pituitary extract (BPE) at a final concentration of 20μg/mL, epidermal growth factor (EGF) at a final concentration of 0.2ng/mL, calcium chloride at a final concentration of 300μM and 1:1000 penicillin-streptomycin (pen-strep; Gibco). Keratinocytes were seeded onto 96-well cell culture plates (Greiner) at a density of 20,000 cells per well and grown for 24 hours in 100μL KSFM.

After 24 hours, KSFM was removed and the keratinocytes were cultured in 100μL KSFM with either 0.2% TritonX (Merck), 5% BHI, or 5% bacterial supernatants. Keratinocytes were cultured overnight for 16 hours in treatment conditions. After 16 hours of treatment on the HiBiT N/TERT keratinocytes, 50μL of the KSFM was transferred to a white-bottom 96-well plate (Sigma-Aldrich) and mixed with 50μL Nano-Glo HiBiT extracellular reagent. The remaining 50μL of KSFM was retained on the cell culture plate with the cells and mixed with 50μL CellTitre-Glo reagent. Both plates were mixed on an orbital shaker for 1 minute, and incubated for 10 minutes at room temperature before luminescence was measured with a SpectraMax M5 Microplate Reader.

## Data availability

Shotgun metagenomic sequencing data is available from the European Nucleotide Archive (ENA, https://www.ebi.ac.uk/ena/browser/home) under project accession number PRJEB87291. Demographic data and skin physiological measurements can be found in **Supplementary File 18**. Large datasets and metabolomics data are available on Figshare at https://figshare.com/projects/ASMP_Dermotype/243932. The authors of this study do not own the rights to the HELIOS dataset, and this dataset is under controlled access to ensure good data governance, responsible data use, and that the dataset is only used for the intended research purposes in compliance with HELIOS study cohort IRB and ethics approval. Access to individual level data for study participants should be submitted to the HELIOS Data Access Committee.

## Code availability

Source code for scripts used to analyze the data are available in a GitHub project at https://github.com/CSB5/ASMP_Dermotype.

## Acknowledgments

The work is supported by funding for the Asian Skin Microbiome Programme (Industry Alignment Fund Pre-Positioning – IAF-PP; H18/01/a0/16) and Asian Skin Microbiome Programme 2.0 (IAF-PP H22/J1/a0/040) from the National Research Foundation, Singapore. Additional support is provided by HELIOS study (IRB No. 2016-11-030), which is supported by Singapore Ministry of Health’s (MOH) National Medical Research Council (NMRC) under its OF-LCG funding scheme (MOH-000271-00), Singapore Translational Research (STaR) funding scheme (NMRC/StaR/0028/2017), the National Research Foundation (Singapore) through the Singapore MOH NMRC and the Precision Health Research (PRECISE) under the National Precision Medicine programme (NMRC/PRECISE/2020), National Cohorts Office through the Singapore MOH NMRC (P2022-01-03 and P2022-02-03) and intramural funding from Nanyang Technological University, Lee Kong Chian School of Medicine and the National Healthcare Group. The HELIOS study is also supported by an outstanding team of administrative and operational staff (**Supplementary File 16**). This research was supported by a National Research Foundation Investigatorship grant (NRFI09-0015) to NN. The computational work for this article was partially performed on resources of the National Supercomputing Centre, Singapore (https://www.nscc.sg) and the A*STAR Computational Resource Center. We thank all investigators, staff members and study participants who made HELIOS and ASMP studies possible.

## Competing interests

The authors declare no competing interests.

## Notes

### Competing Interest Statement

The authors have declared no competing interest.

### Summary of Updates

Abstract, introduction, and discussion sections updated for clarify.

