## Supplementary File 8 for "Large-scale skin metagenomics reveals extensive prevalence, coordination, and functional adaptation of skin microbiome dermotypes across body sites"

| Aerobic conditions | | | | |
|---|---|---|---|---|
| | SE | SH | ML | CA |
| SE inhibits | * | | * | No growth |
| SH inhibits | | * | * | |
| ML inhibits | | | | |
| CA inhibits | | | | |

| Hypoxic conditions | | | | |
|---|---|---|---|---|
| | SE | SH | ML | CA |
| SE inhibits | | # | * | |
| SH inhibits | | | * | |
| ML inhibits | | # | | |
| CA inhibits | | | | * |

| Respective favorable conditions | | | | |
|---|---|---|---|---|
| | SE | SH | ML | CA |
| SE inhibits | | | | * |
| SH inhibits | | | | * |
| ML inhibits | | | | # |
| CA inhibits | | | | |

SE: *Staphylococcus epidermidis*

SH: *Staphylococcus hominis*

ML: *Micrococcus luteus*

CA: *Cutibacterium acnes*

### zone of complete inhibition

* zone of partial growth inhibition

cultured in aerobic conditions

cultured in hypoxic conditions

# SE & SH

Aerobic Aerobic

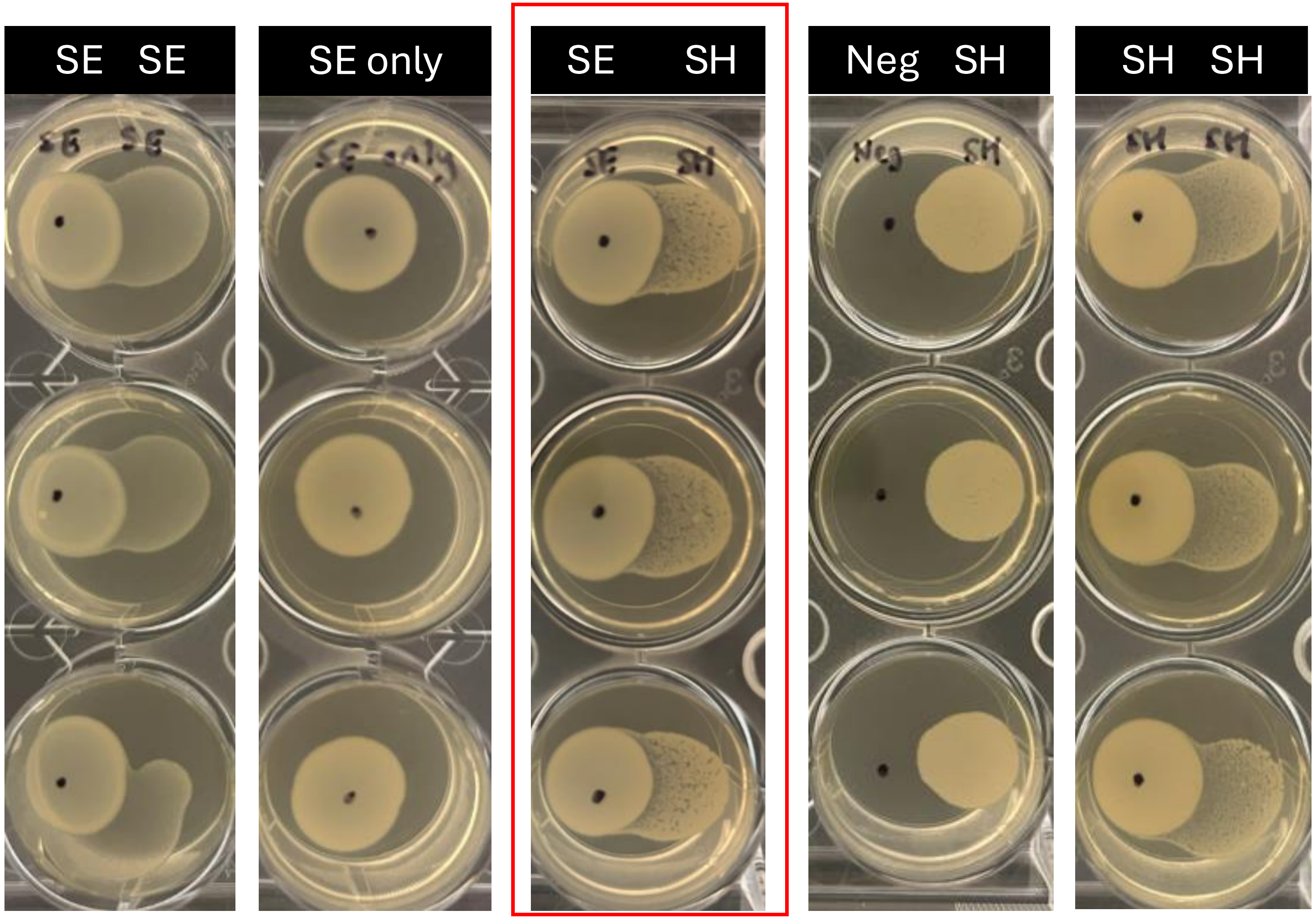

# SE & ML

Aerobic

Aerobic

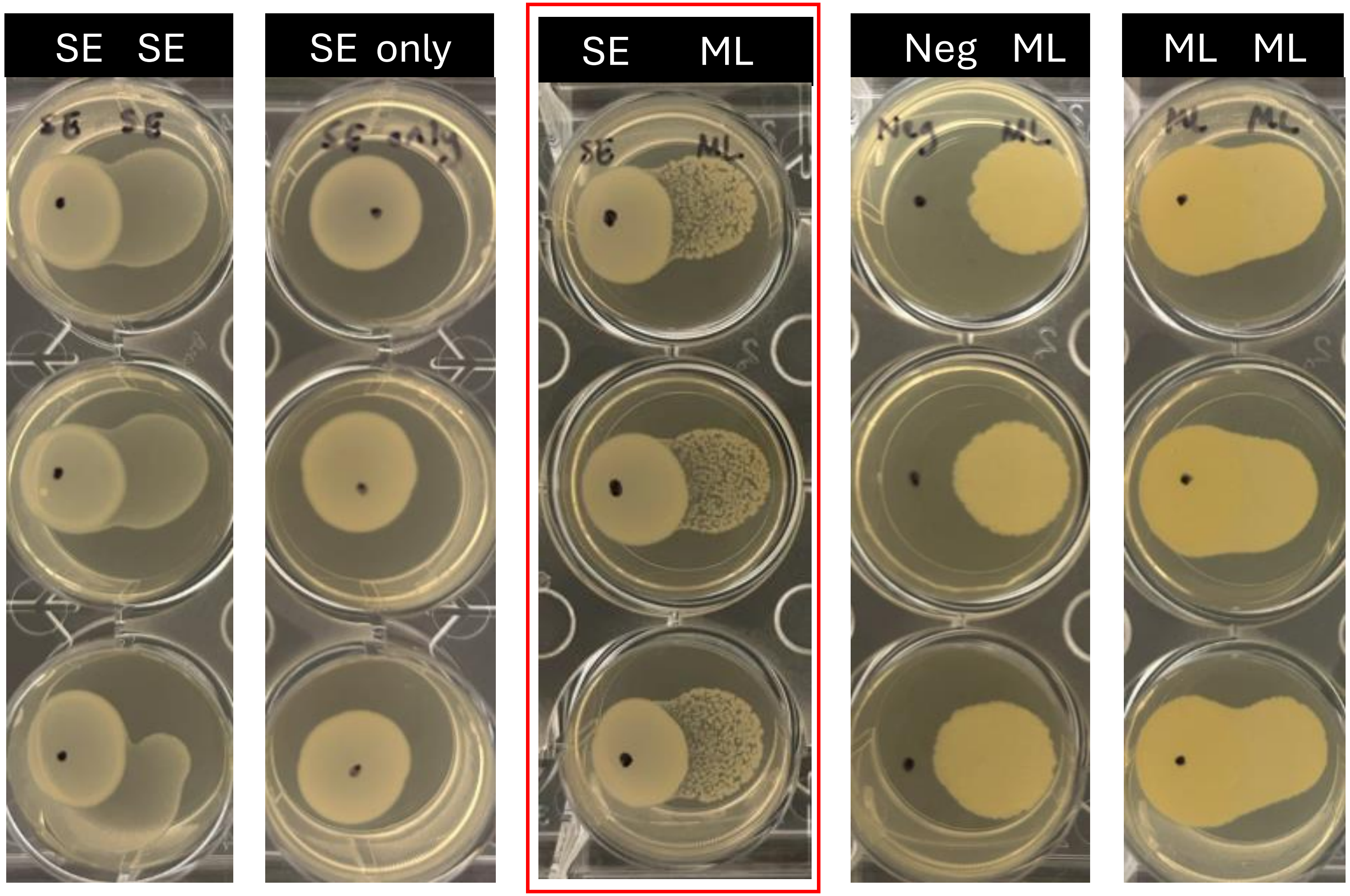

# SH & SE

Aerobic Aerobic

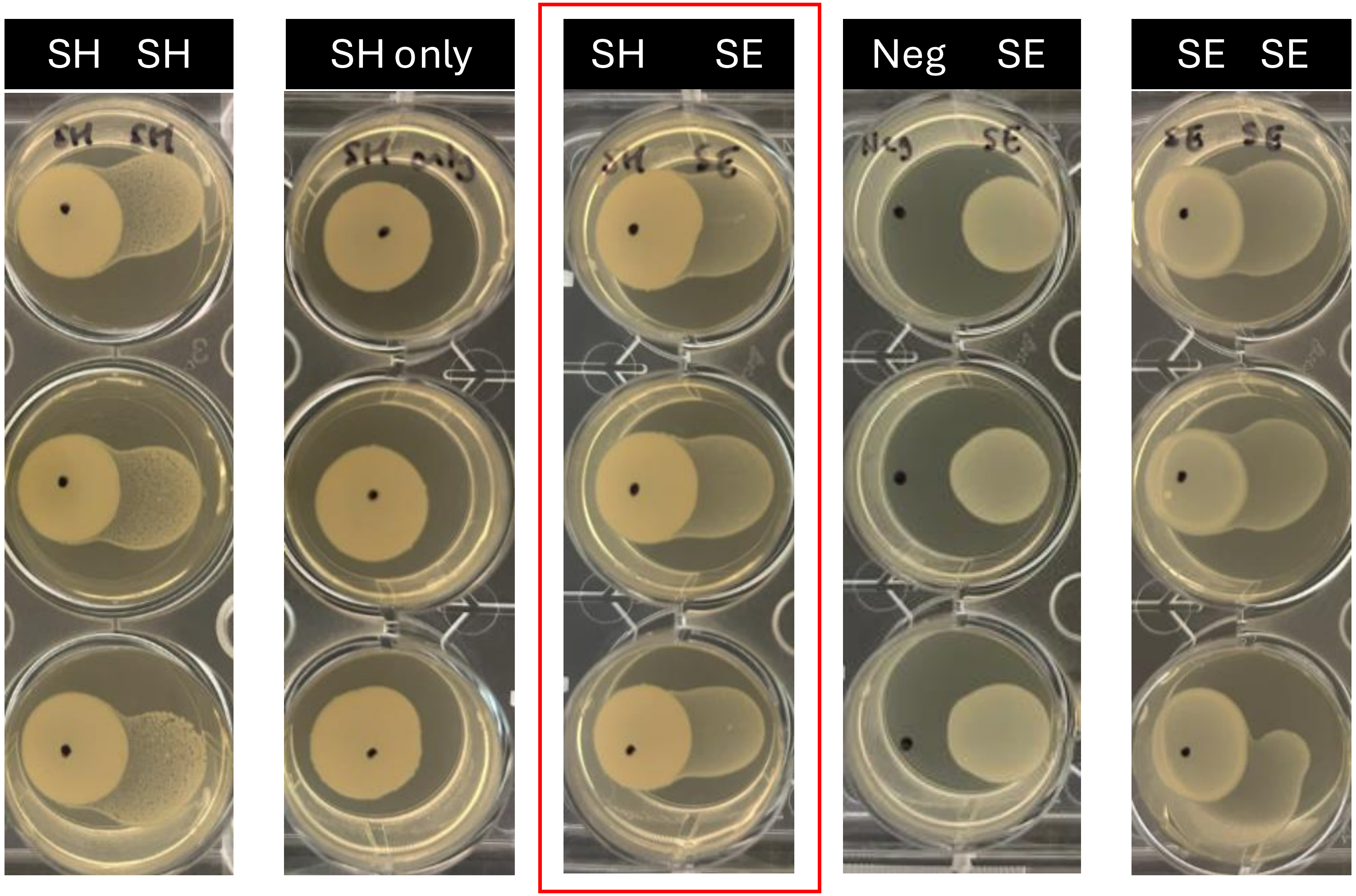

# SH & ML

Aerobic Aerobic

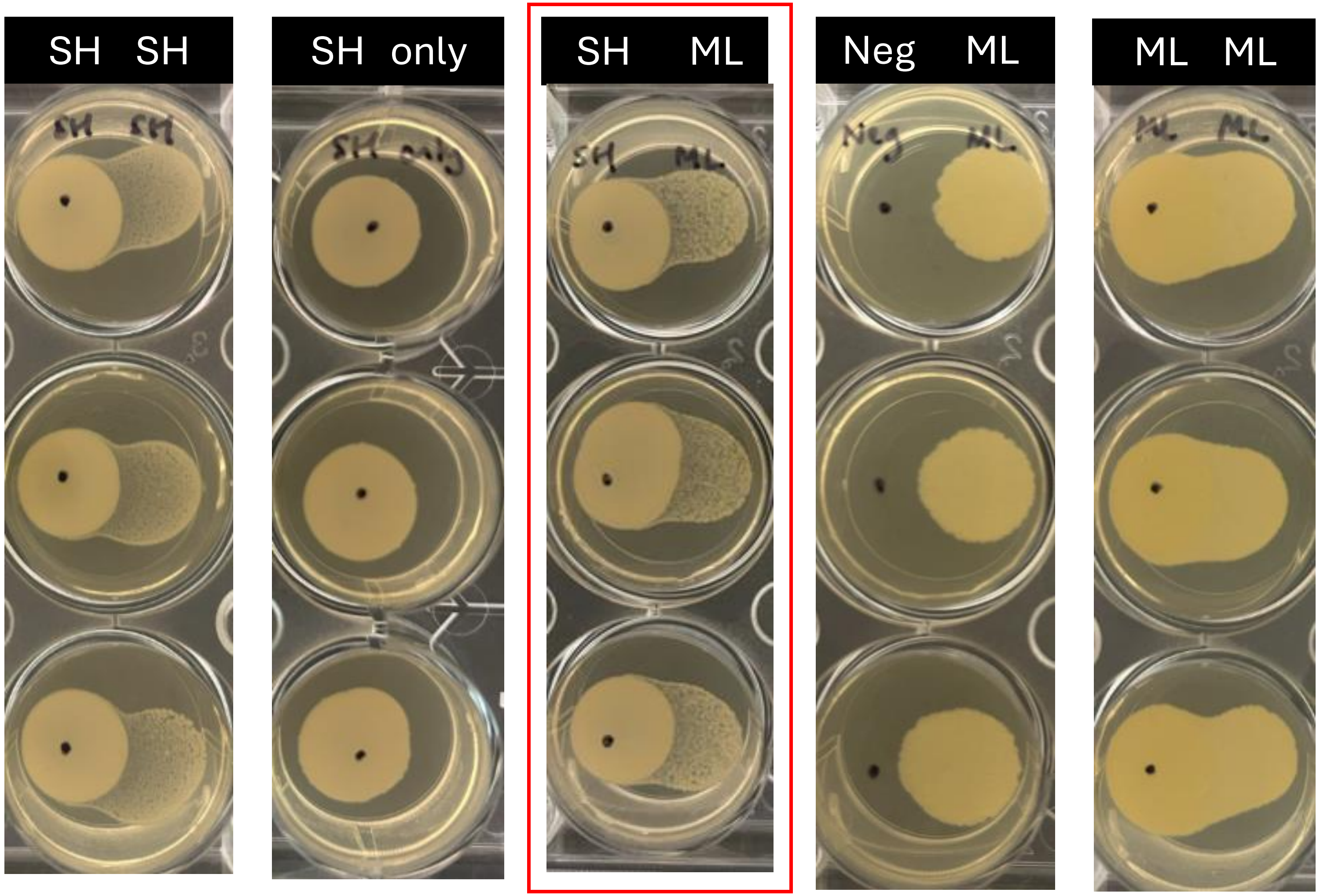

# ML & SE

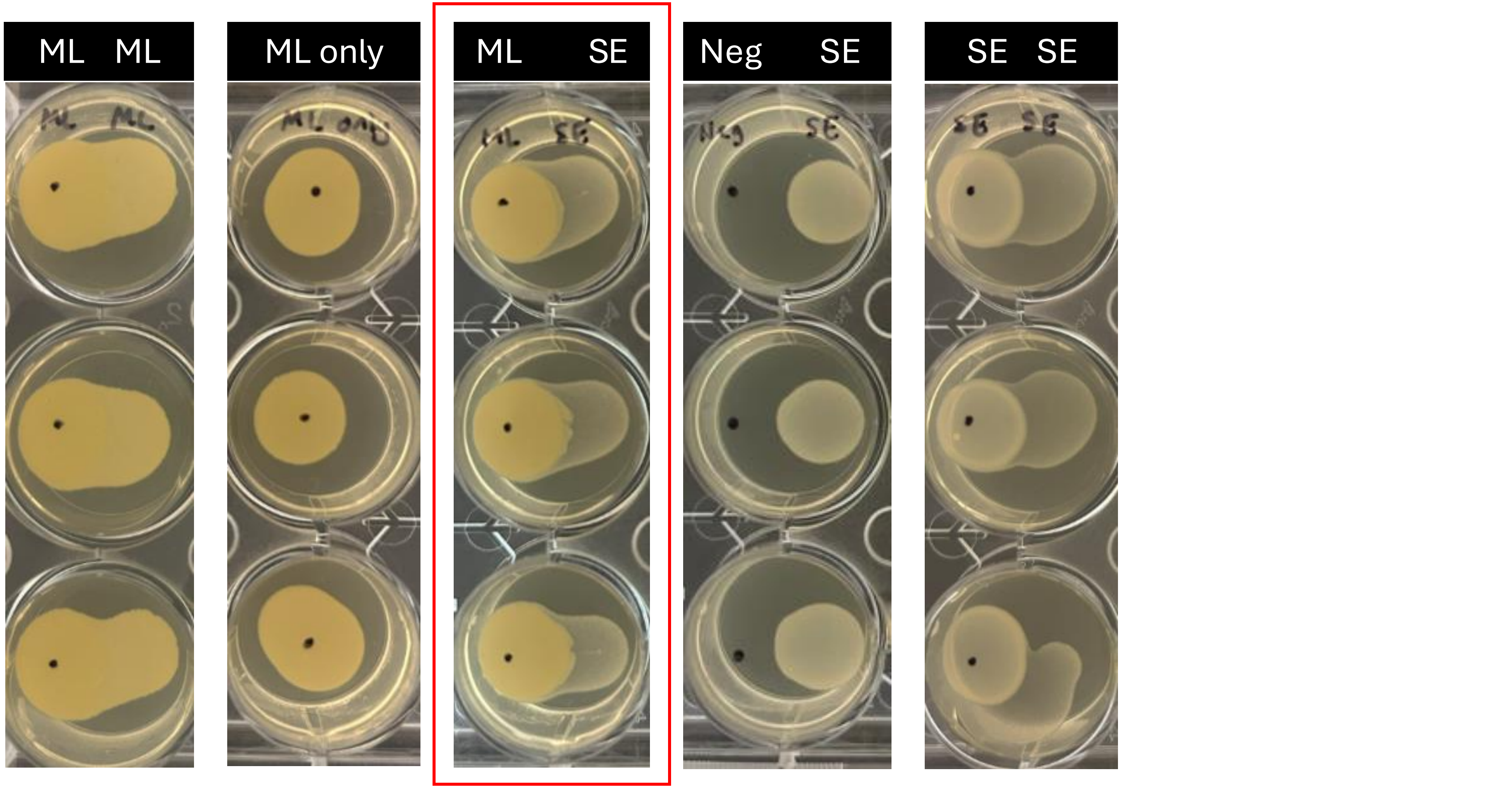

# ML & SH

Aerobic

Aerobic

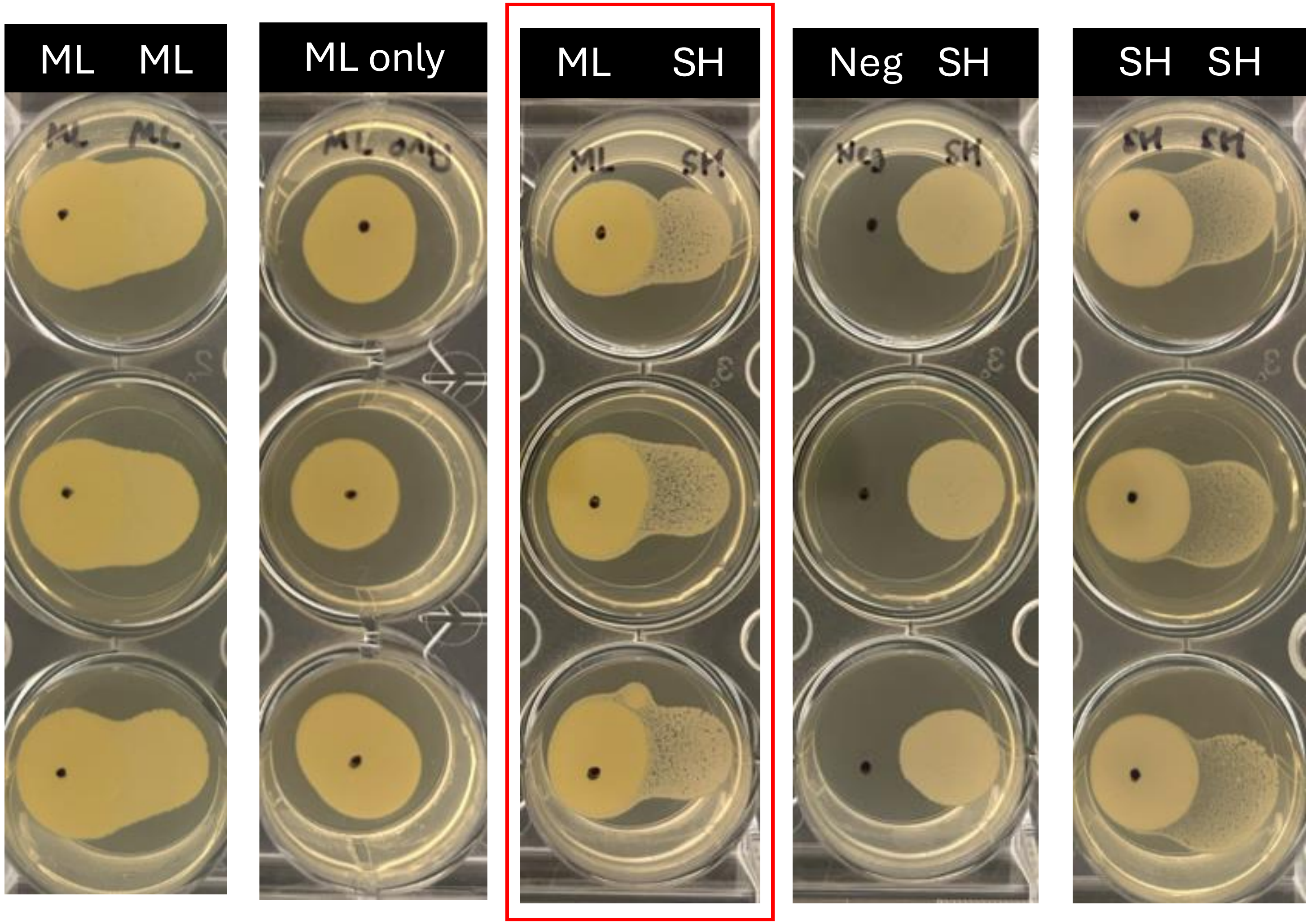

# CA & SE

Aerobic

Aerobic

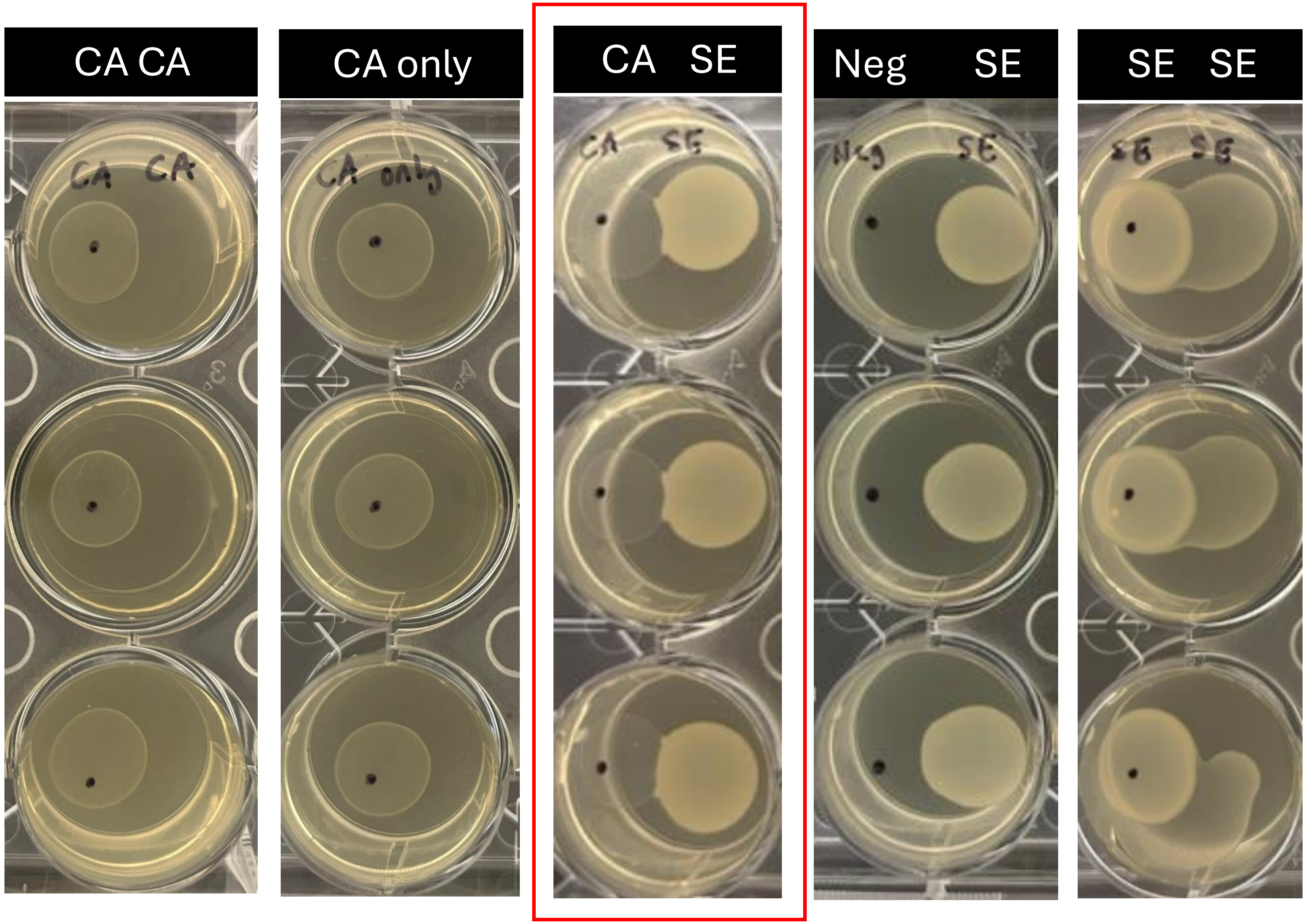

# CA & SH

Aerobic

Aerobic

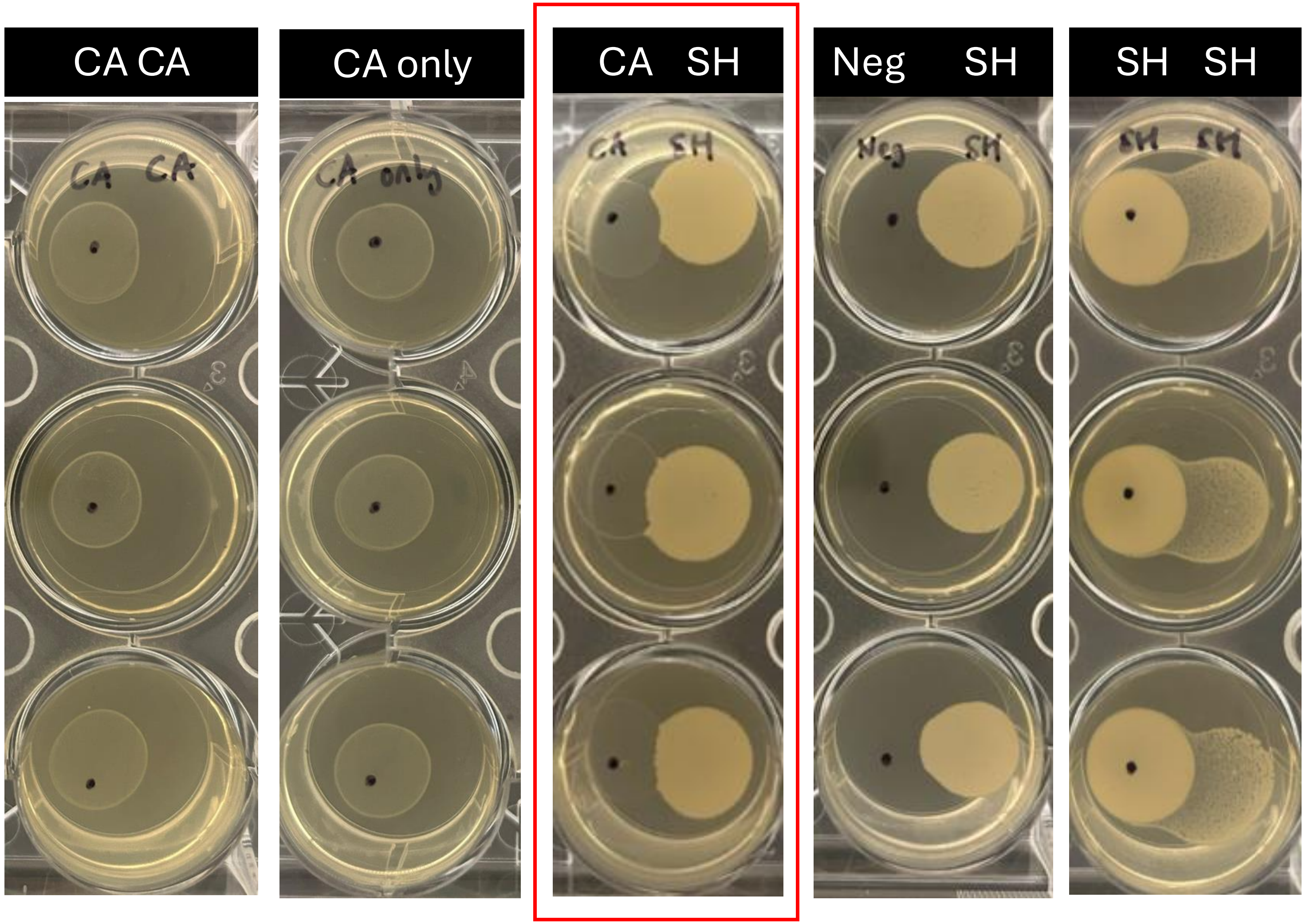

# CA & ML

Aerobic

Aerobic

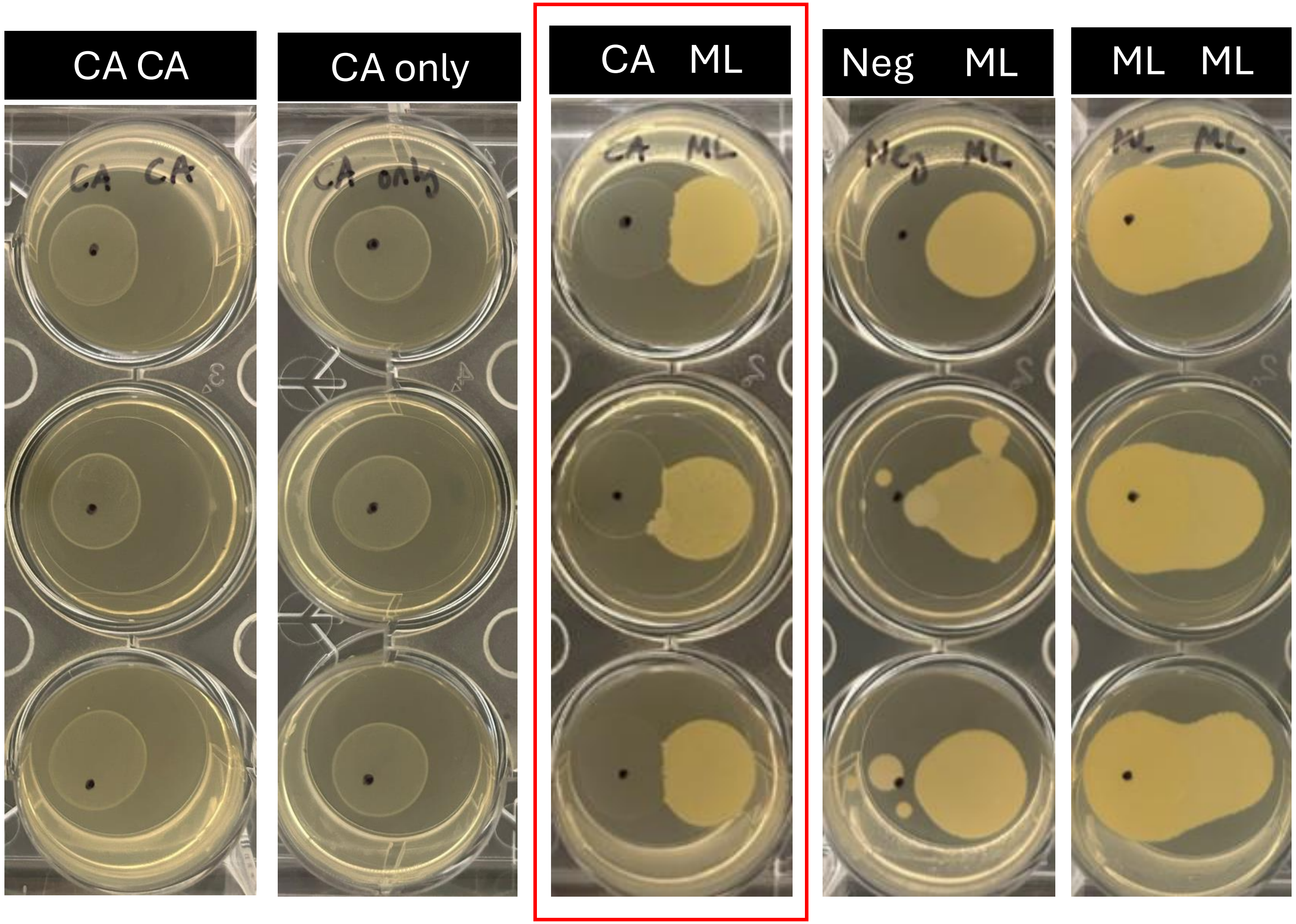

# SE & SH

Hypoxic Hypoxic

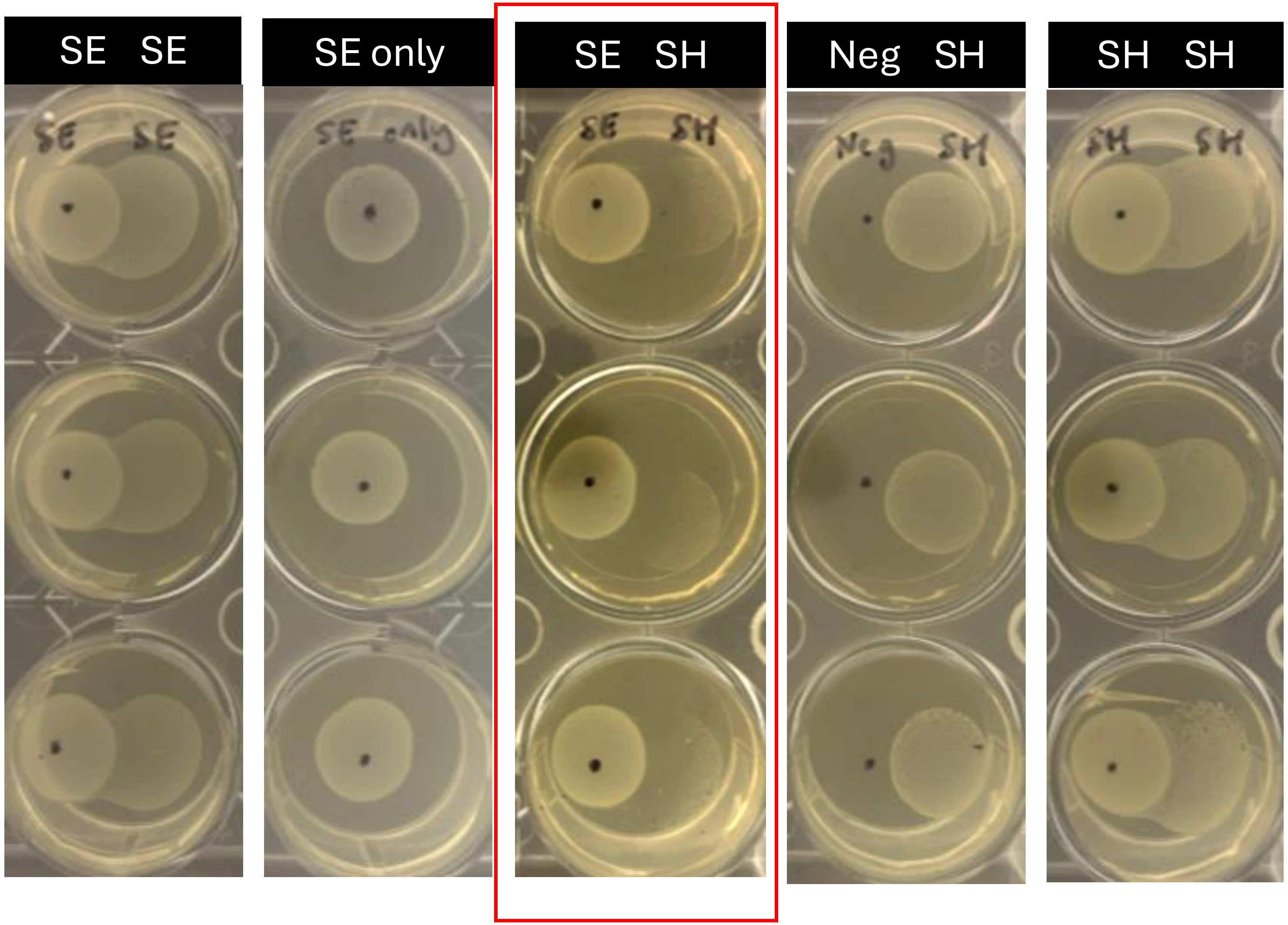

# SE & ML

Hypoxic Hypoxic

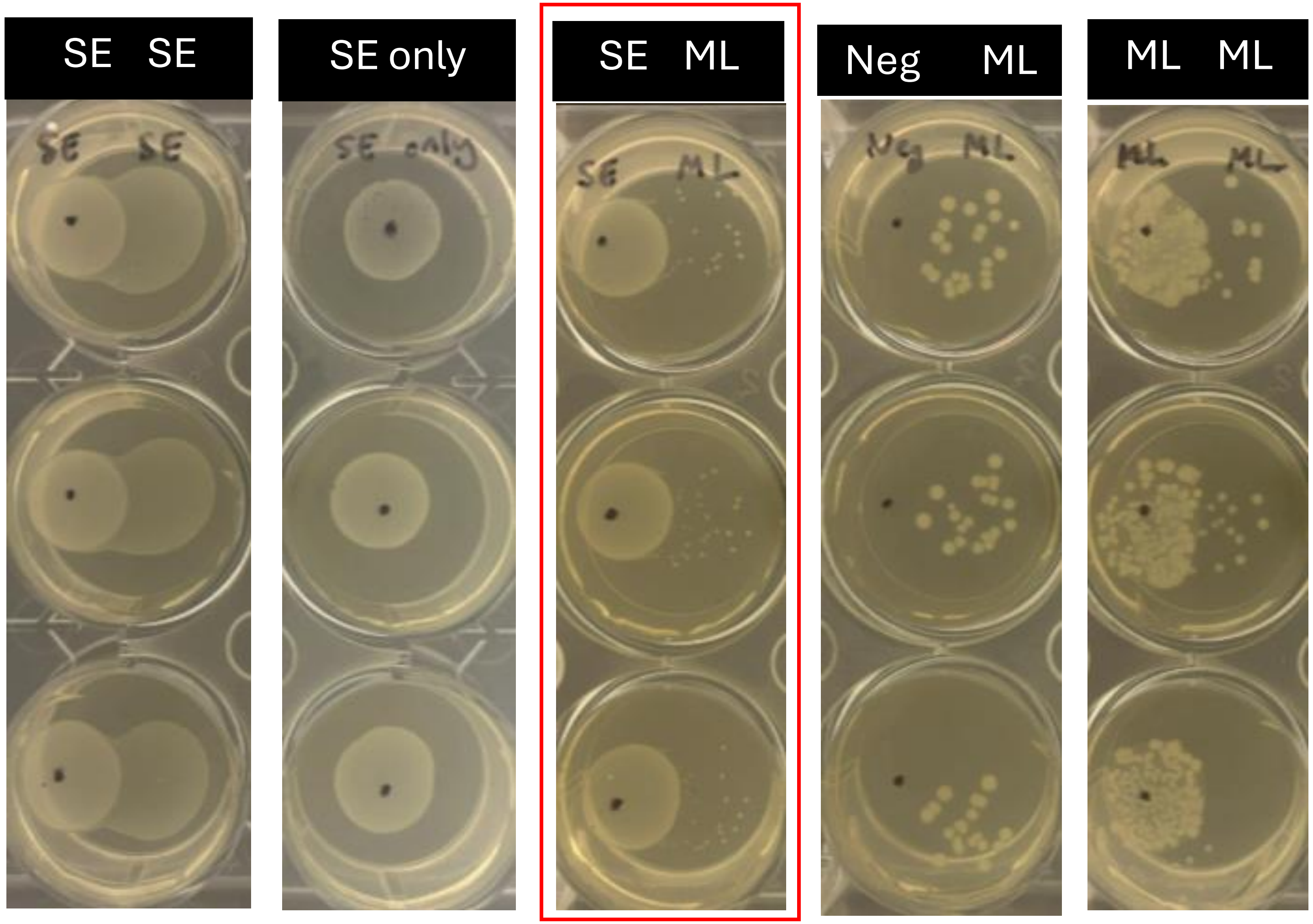

# SE & CA

Hypoxic Hypoxic

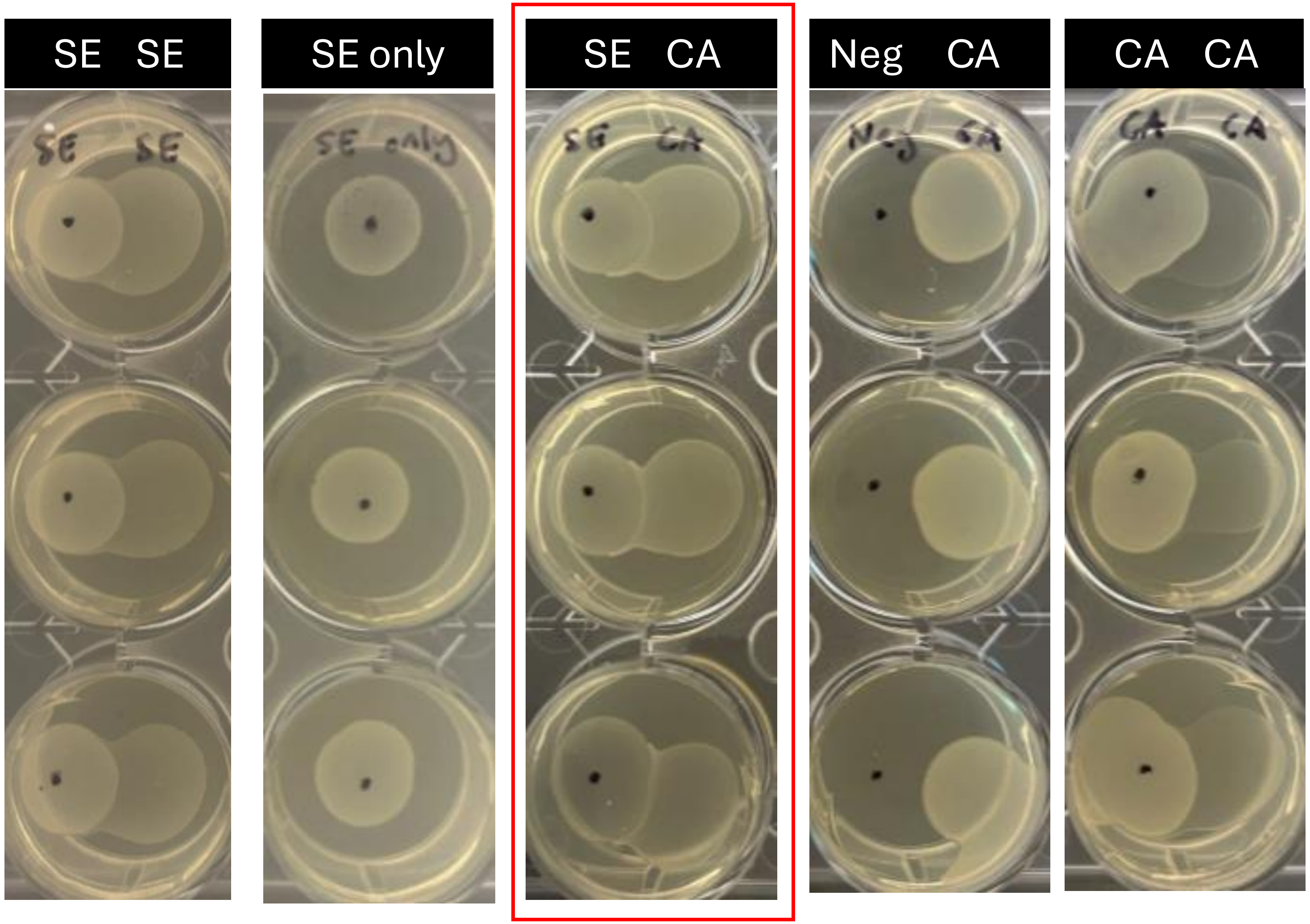

# SH & SE

**Hypoxic** **Hypoxic**

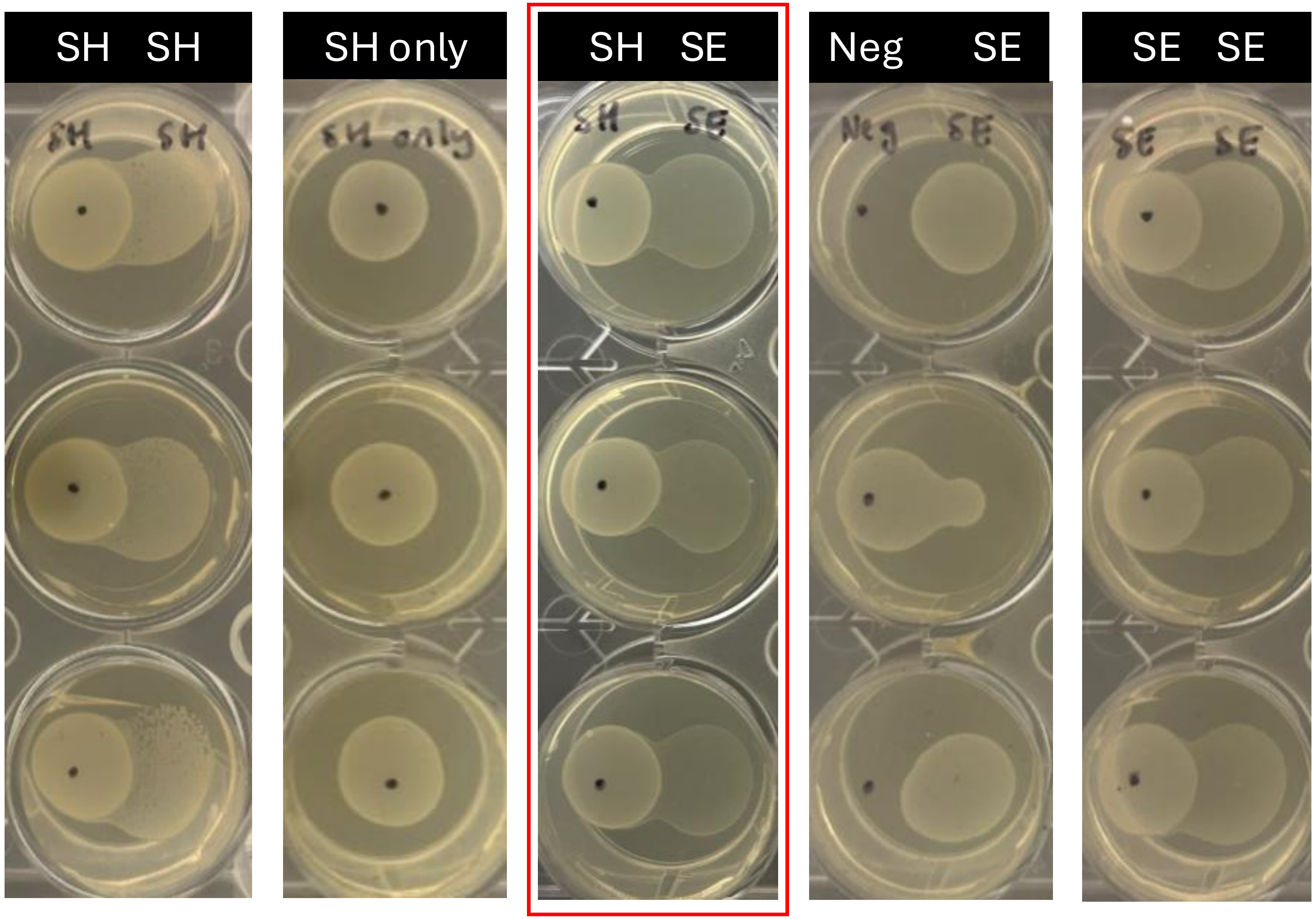

# SH & ML

Hypoxic Hypoxic

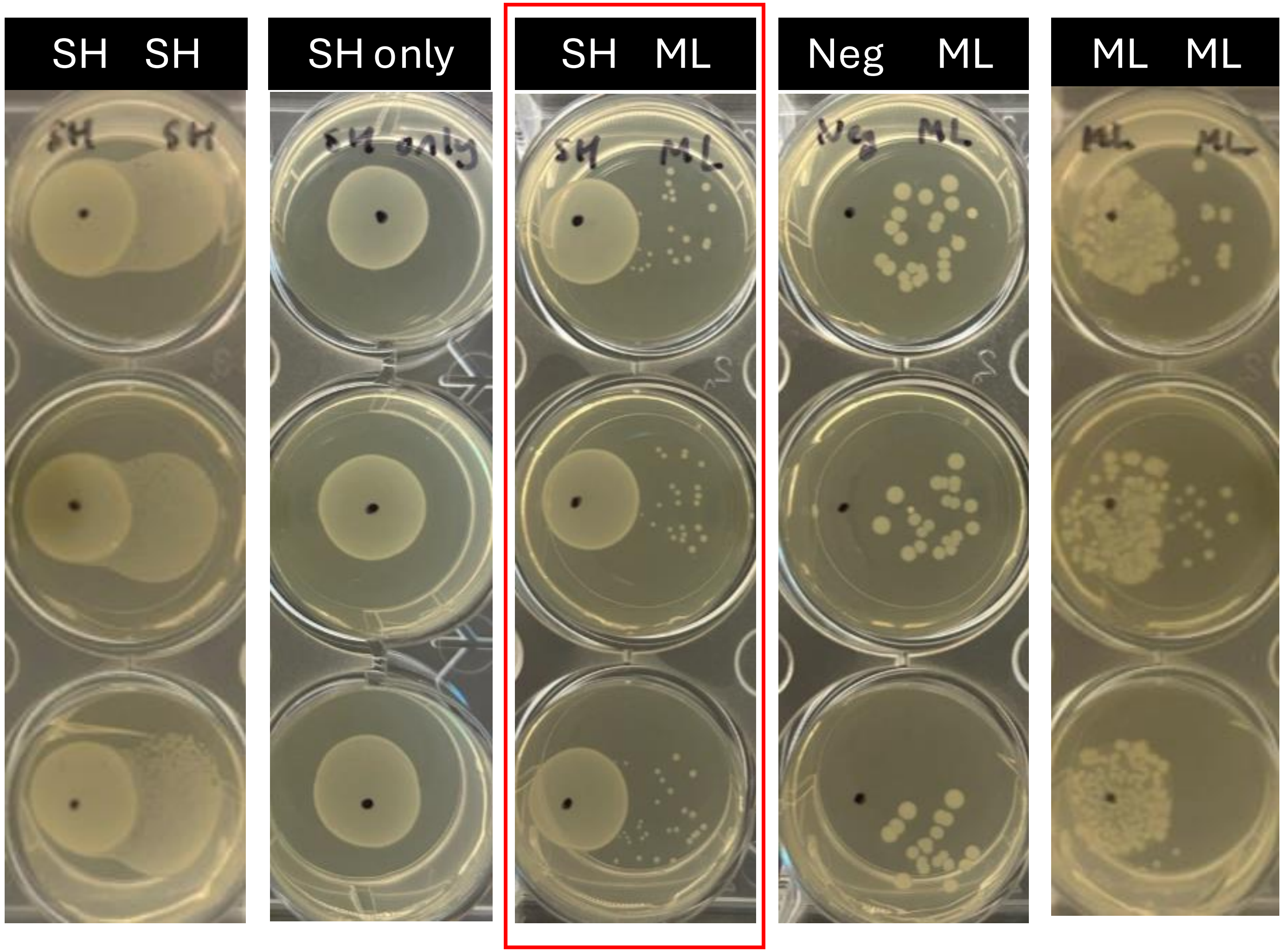

# SH & CA

**Hypoxic** **Hypoxic**

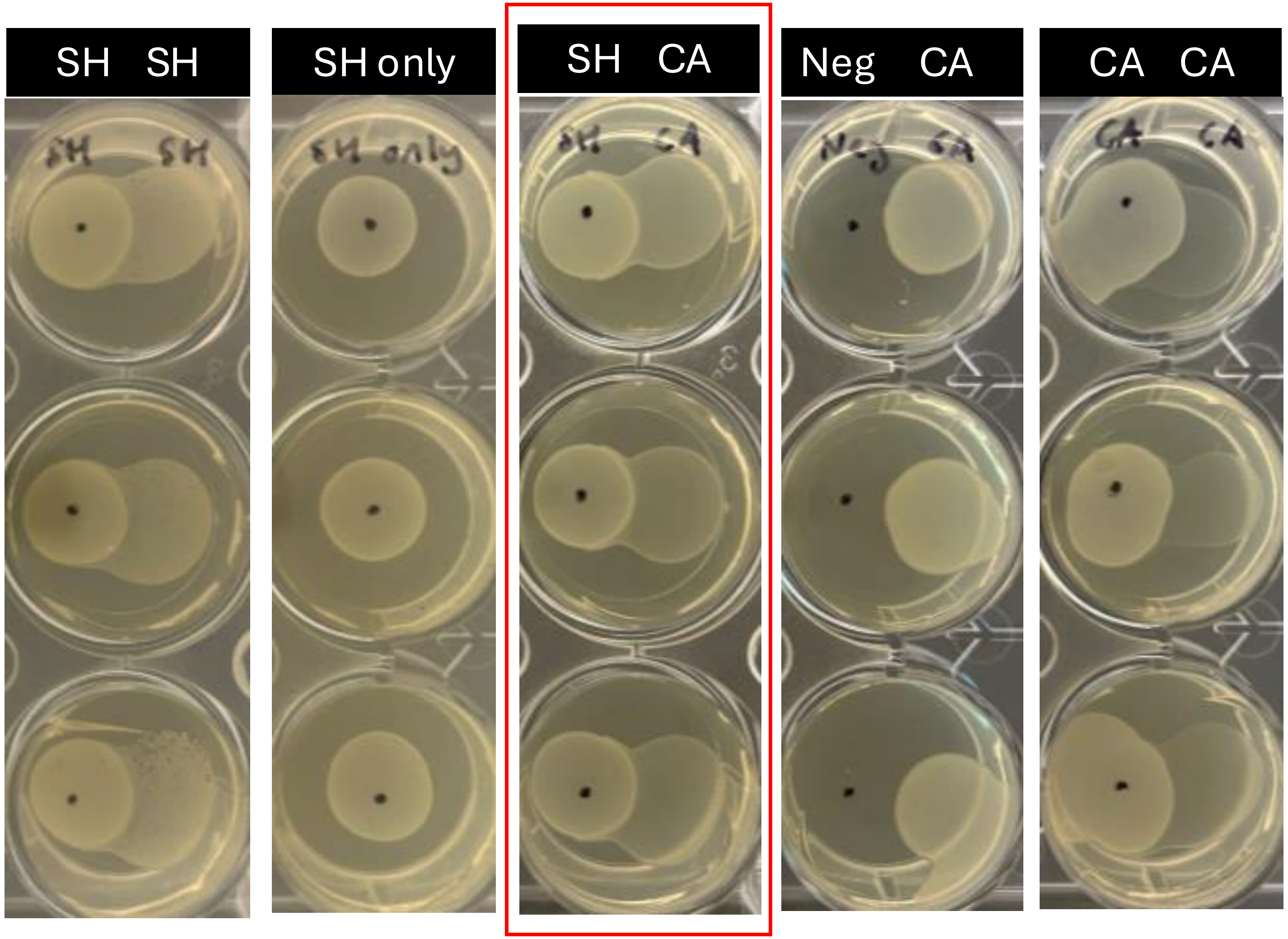

# ML & SE

Hypoxic Hypoxic

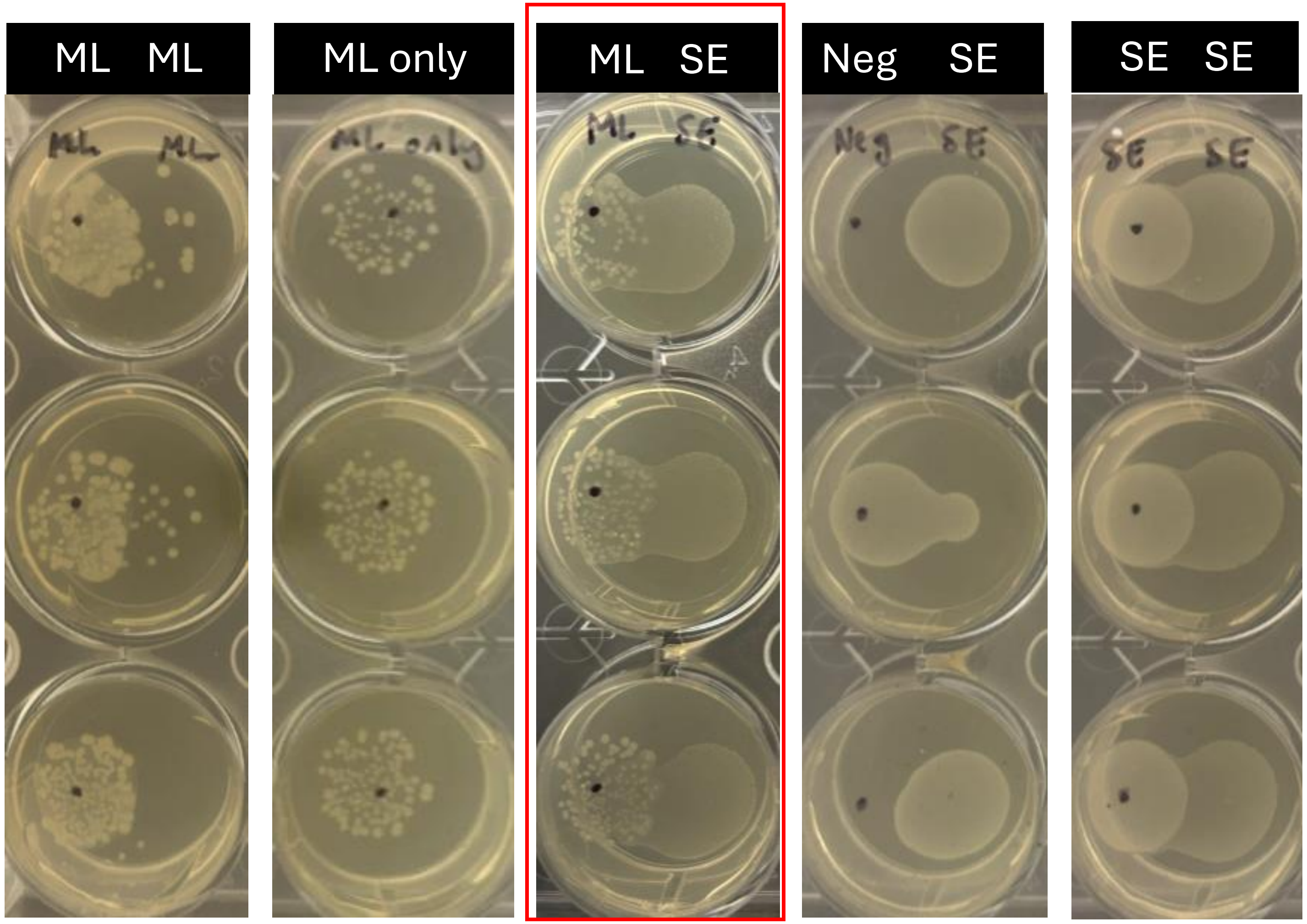

# ML & SH

**Hypoxic** **Hypoxic**

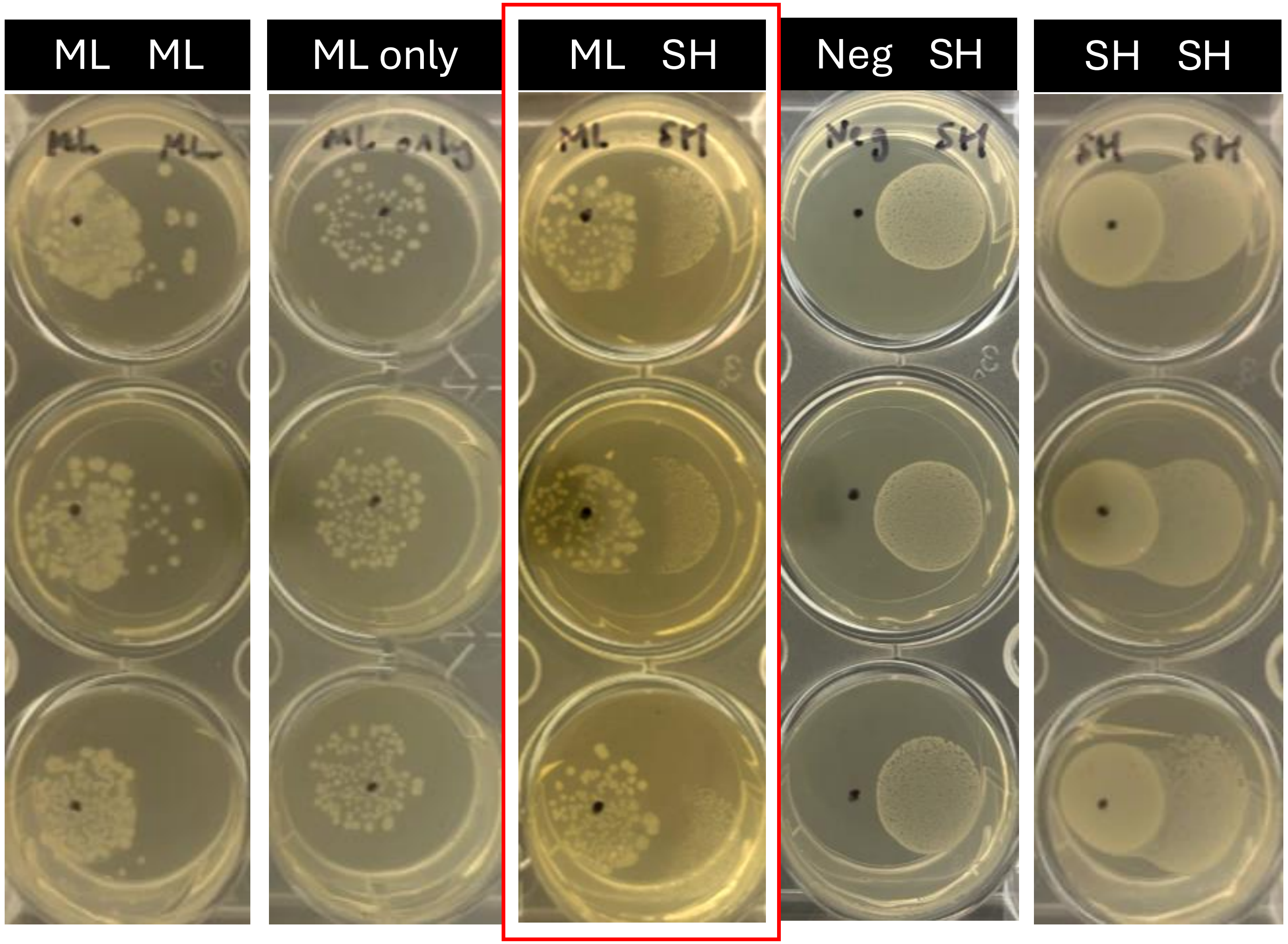

# ML & CA

**Hypoxic**

**Hypoxic**

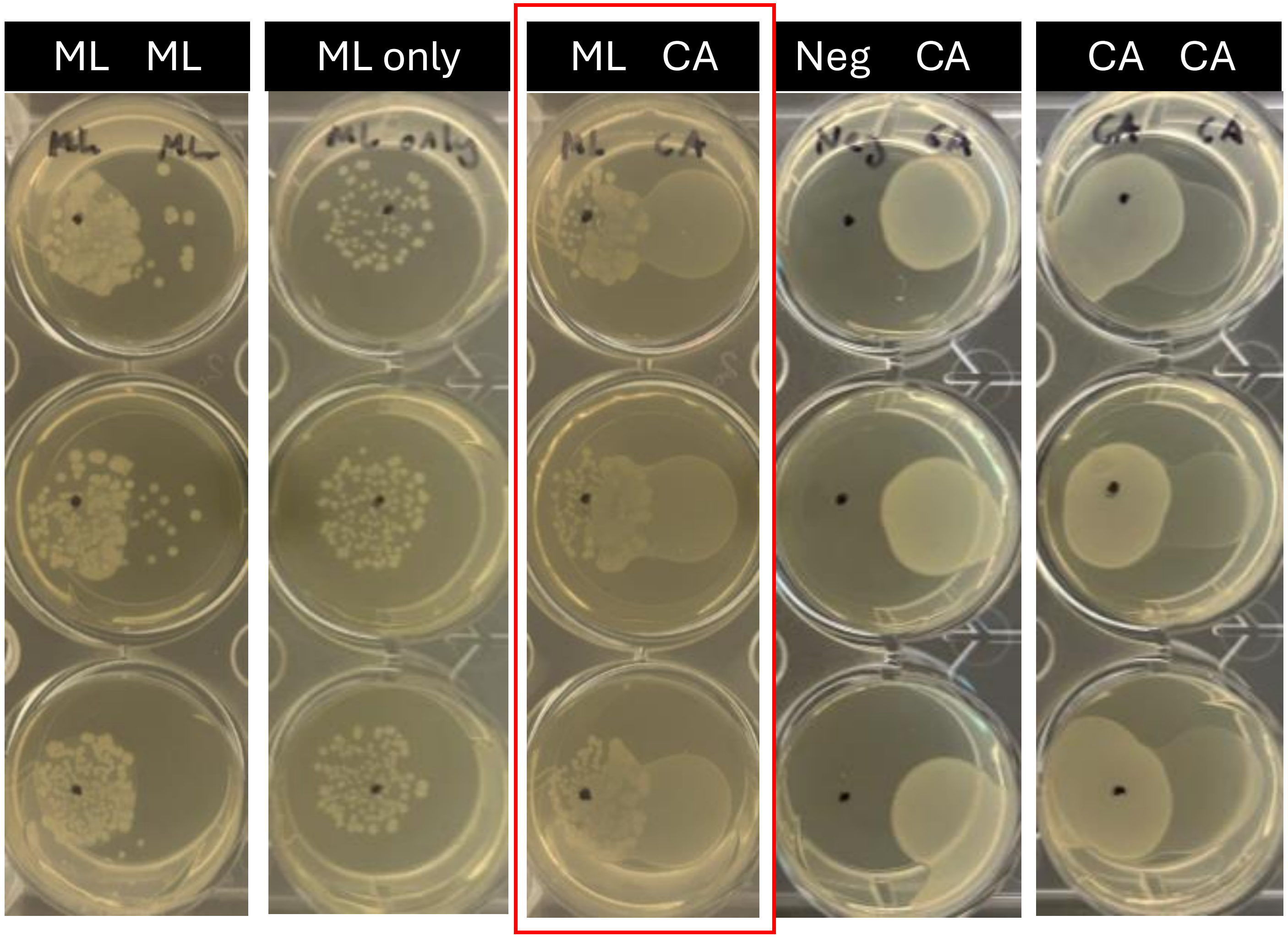

# CA & SE

**Hypoxic** **Hypoxic**

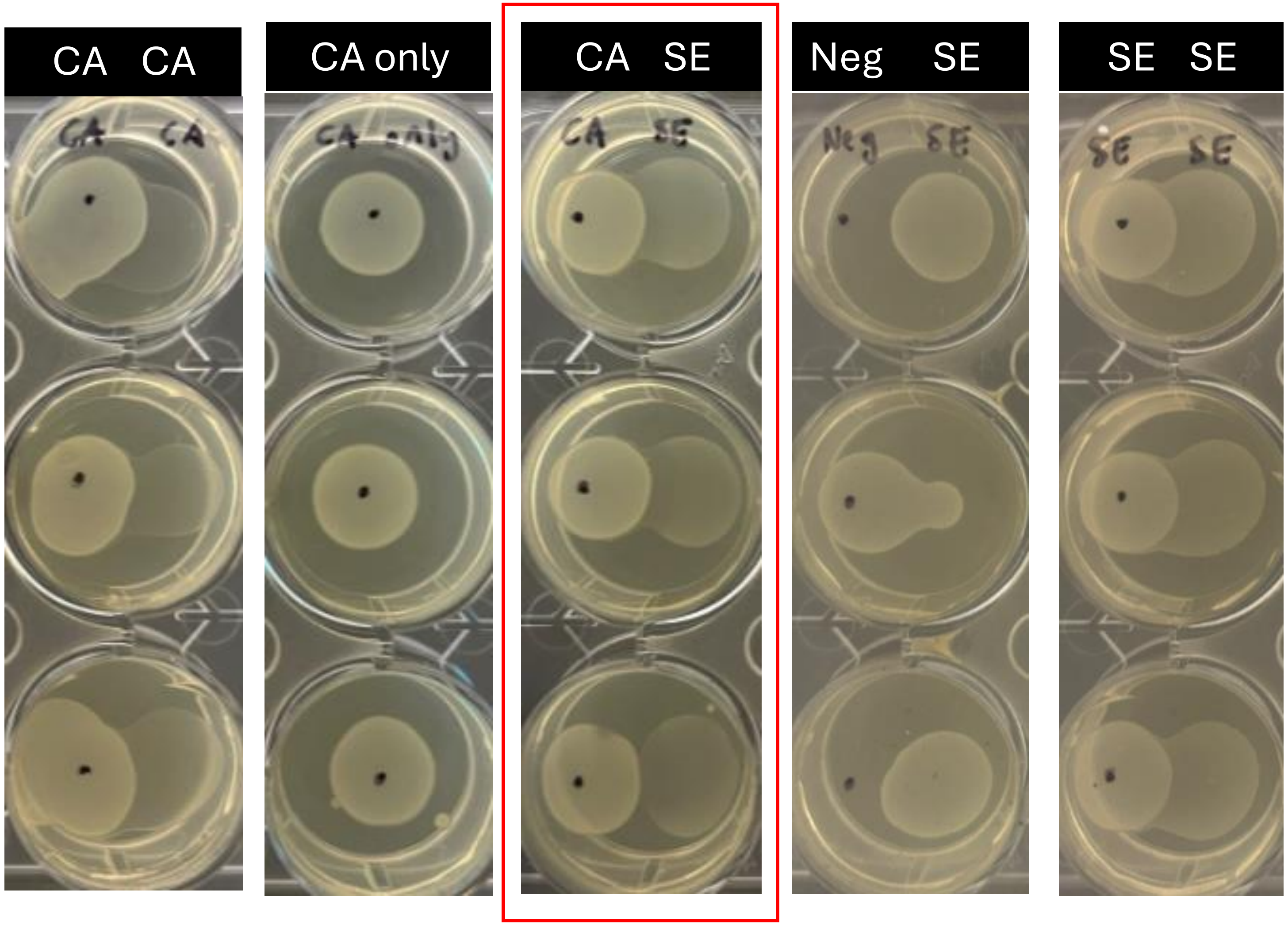

# CA & SH

Hypoxic Hypoxic

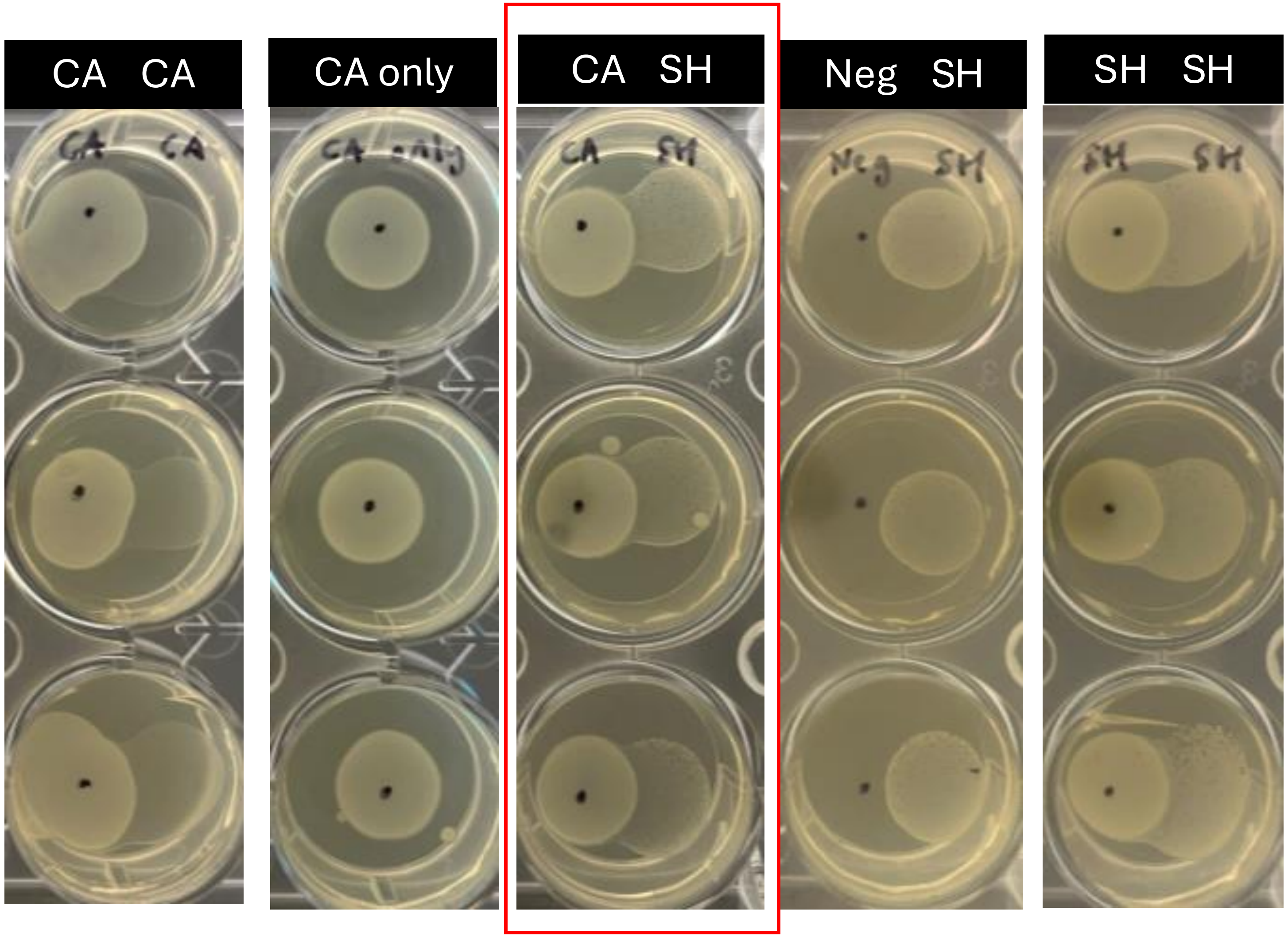

# CA & ML

Hypoxic Hypoxic

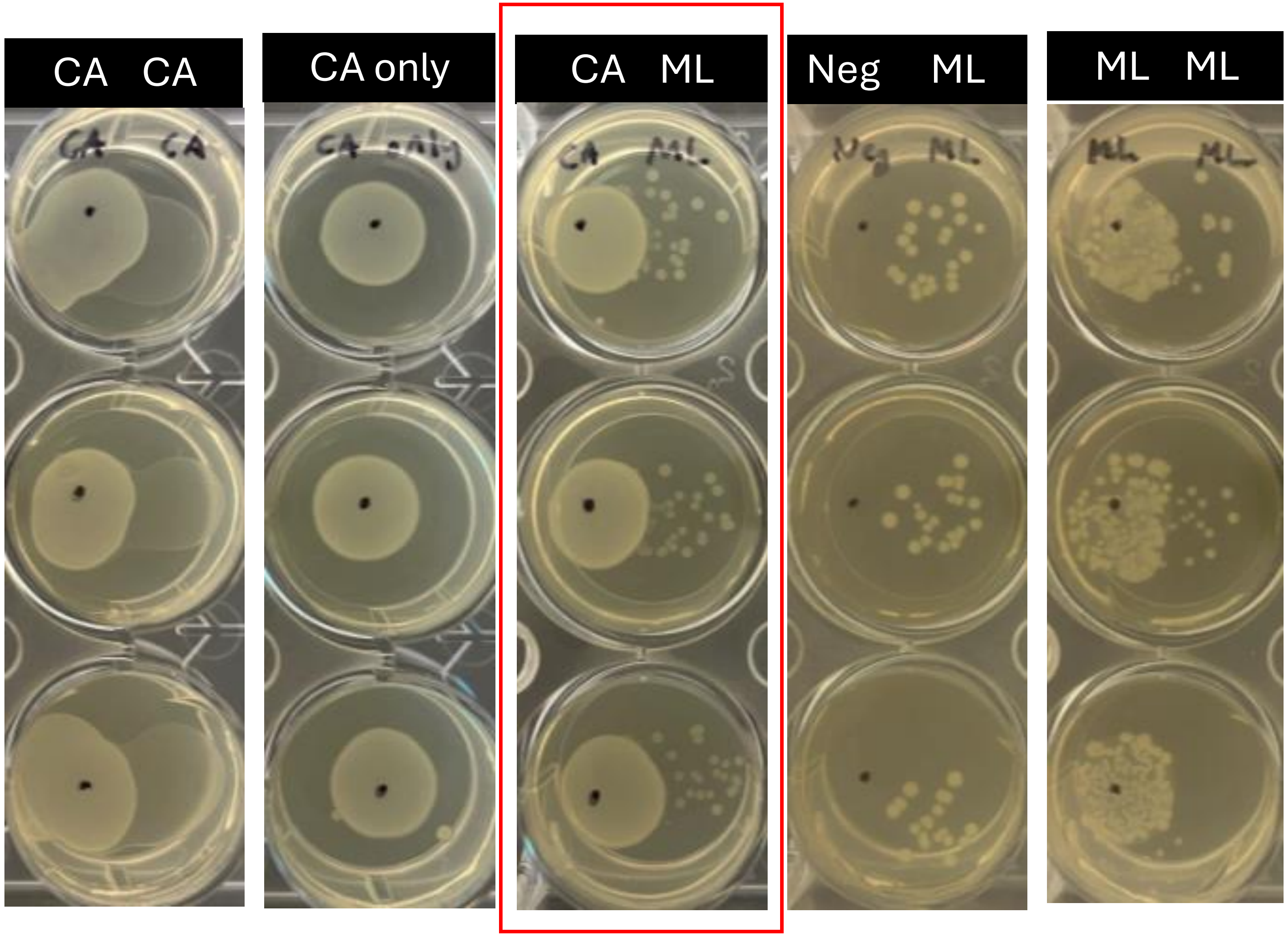

# SE & CA

Aerobic Hypoxic

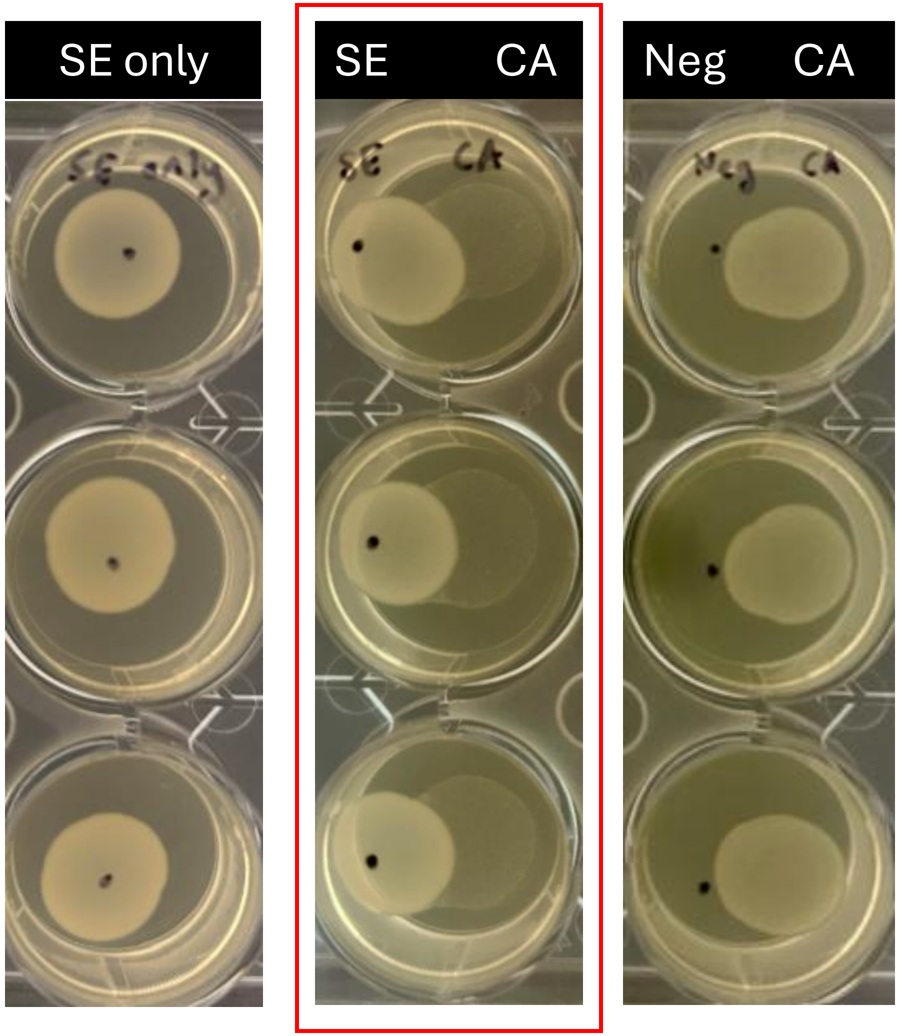

# SH & CA

Aerobic

Hypoxic

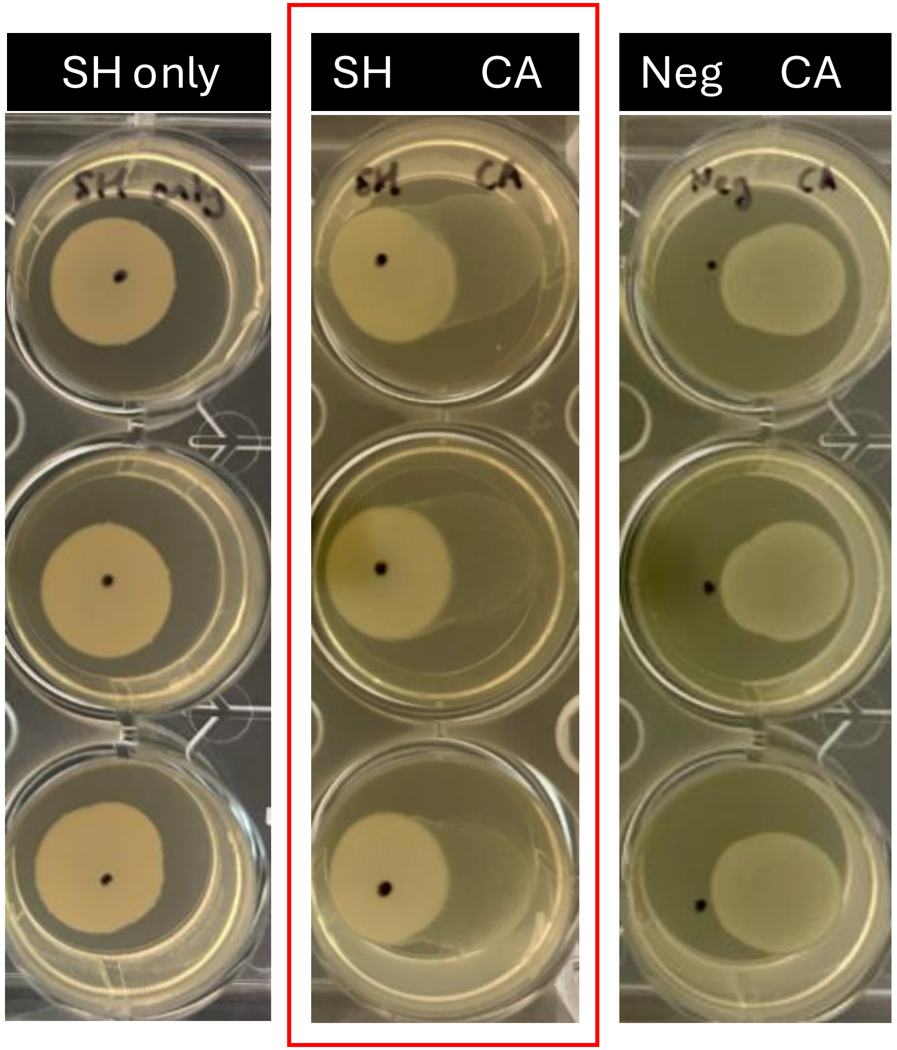

# ML & CA

Aerobic

Hypoxic

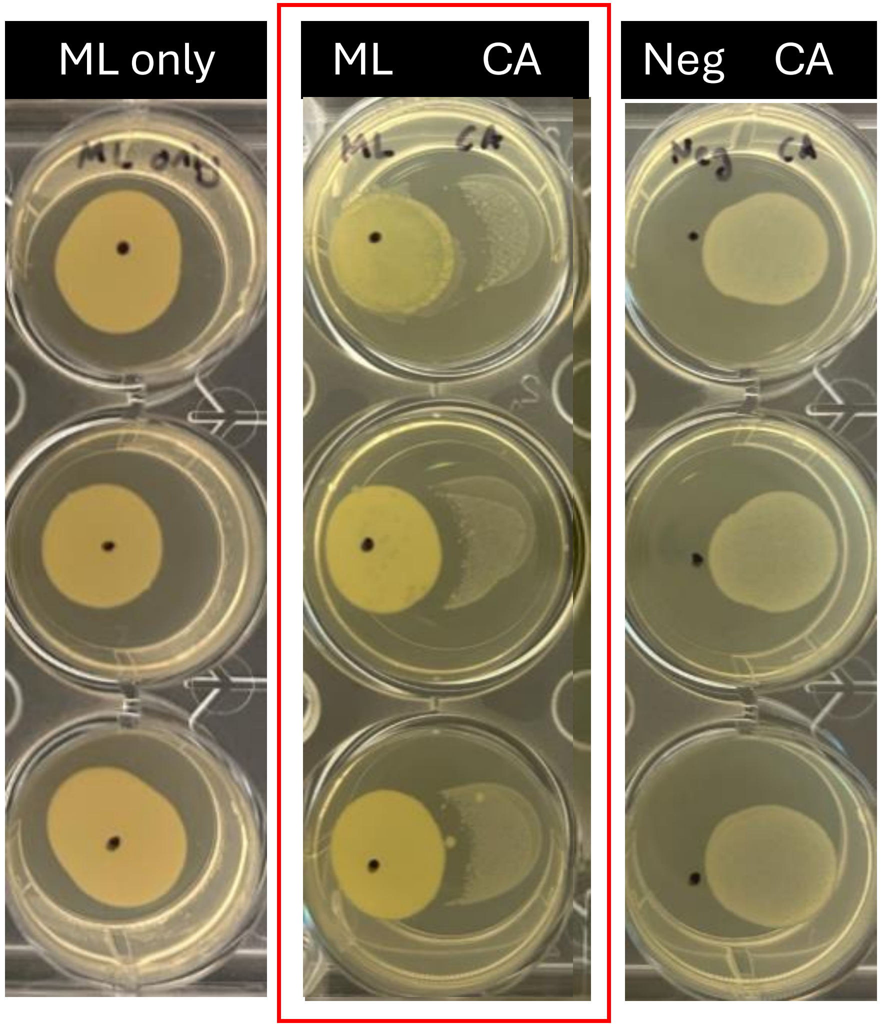

# CA & SE

Hypoxic

Aerobic

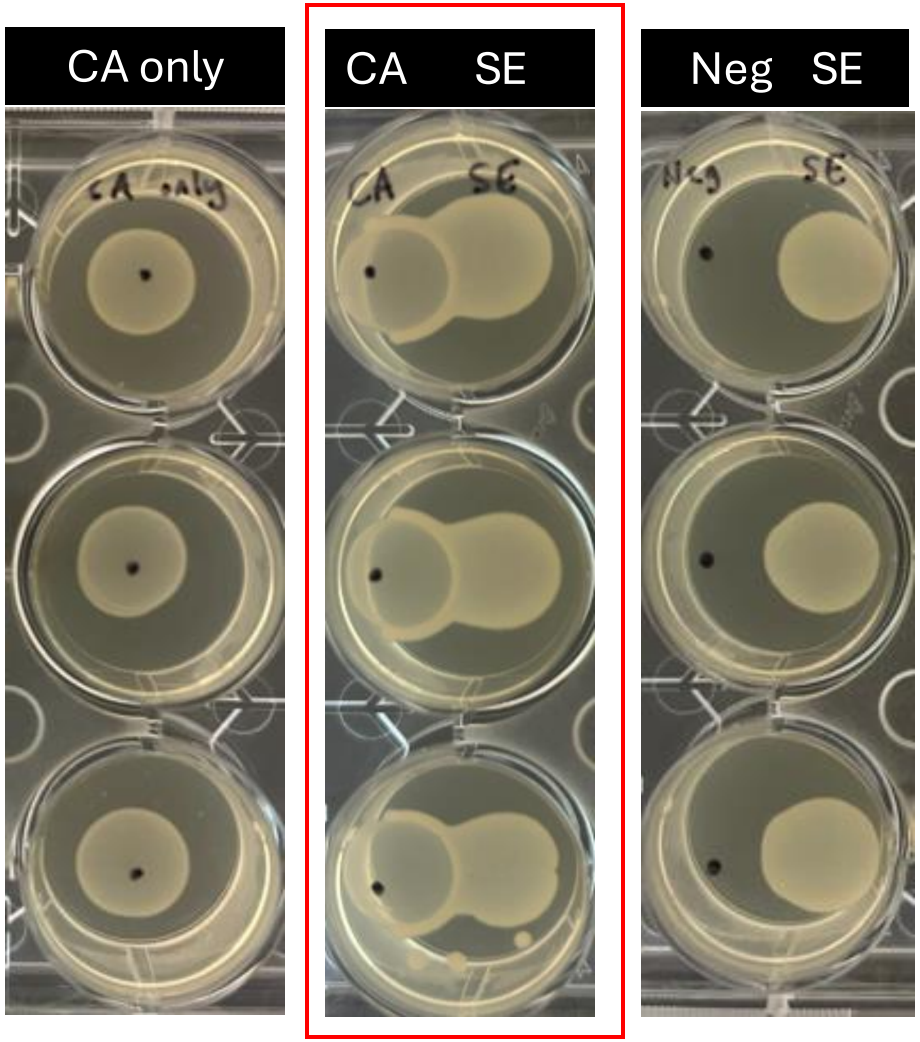

# CA & SH

**Hypoxic** **Aerobic**

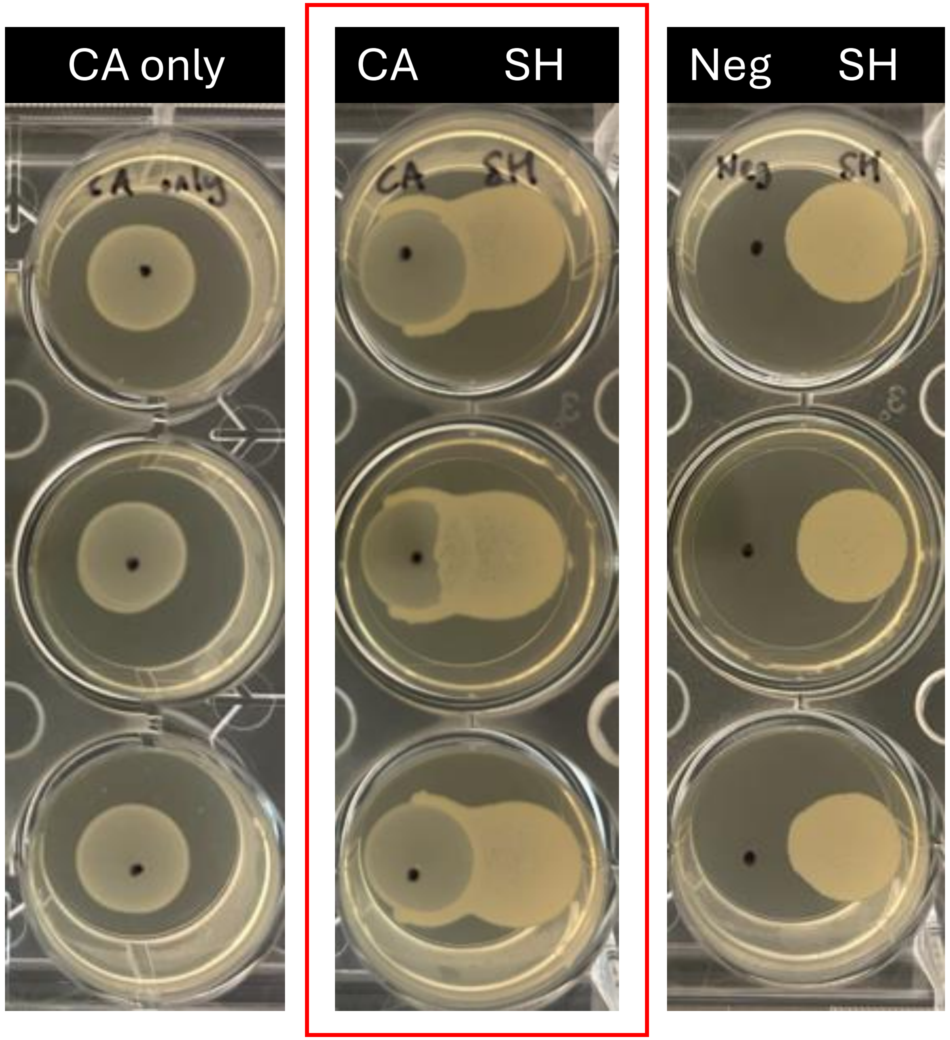

# CA & ML

Hypoxic

Aerobic

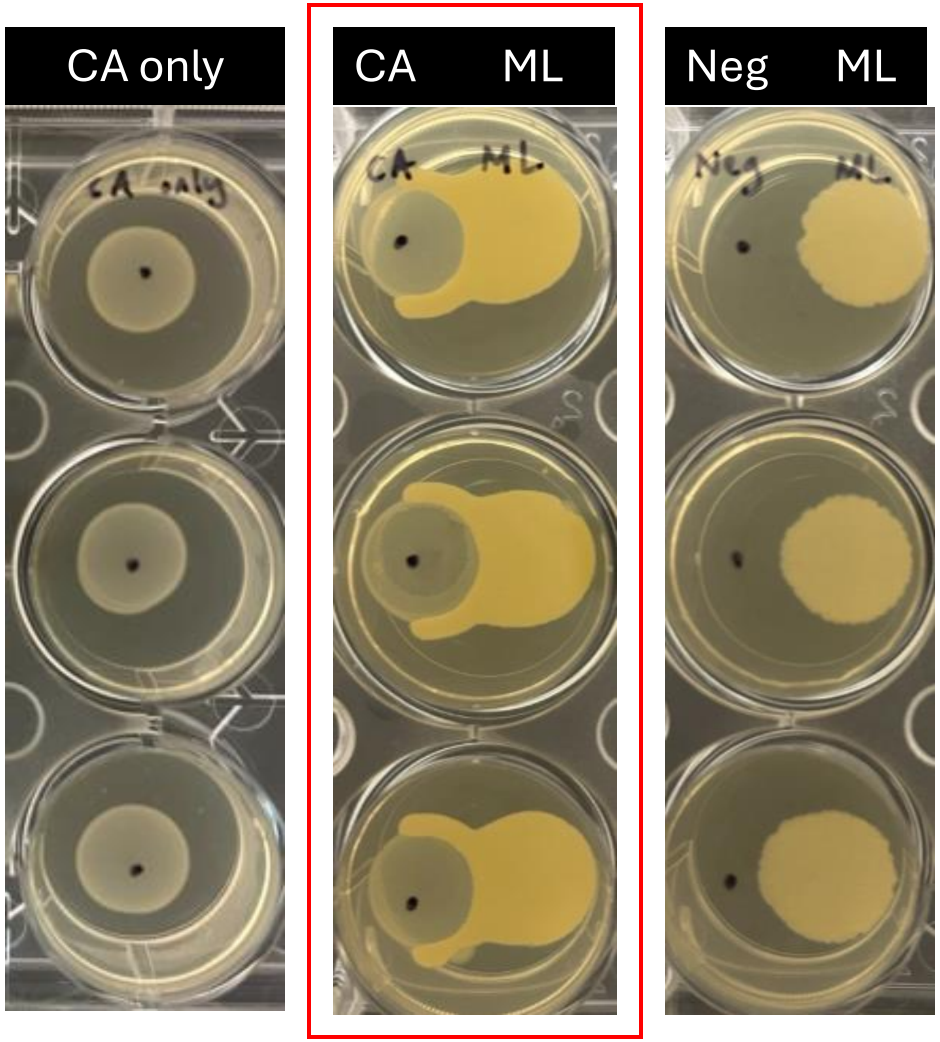
