## Supplementary File 14 for "Large-scale skin metagenomics reveals extensive prevalence, coordination, and functional adaptation of skin microbiome dermotypes across body sites"

| Date (DDMMYYYY): | |  | | | Start Time: |
| --- | --- | --- | --- | --- | --- |
| Remarks: | | | | | |
| Staff Name: |  | | Signature: |  | End Time: |

The HELIOS Study Asian Skin Microbiome Project Questionnaire

A. **Skin Health Questionnaire**
1. Do you have itchy skin? Yes, or No (if no, skip to question 3)
2. If yes, please circle or specify body site(s).
   a. Scalp
   b. Face
   c. Arms
   d. Arm pits
   e. Trunk (body)
   f. Legs
   g. Feet
   h. Others	________

3. Do you feel you have sensitive skin? Yes, or No (if no, skip to question 6)
4. If yes, please circle or specify body site(s).
   a. Scalp
   b. Face
   c. Arm
   d. Arm pits
   e. Trunk (body)
   f. Legs
   g. Feet
   h. Others	________
5. If Yes, what is your degree of overall skin irritation during the past 3 days? Using an X, indicate your degree of skin irritation felt during the past three days ticking on the square boxes below (0 = absence of irritation, 10 = intolerable irritation)

Skin scale	0--------------------------------10

Min	Max

Please indicate by cross or tick below the intensity of each of the following during the past days.

0 = none, 10 = intolerable:

Skin condition felt:

0	1	2	3	4	5	6	7	8	9	10

Tingling											

Burning											

Heat											

Tautness											

Itching											

Pain											

General											

Hot flashes											

6. Are you undergoing any current medical treatment for a skin problem? Yes, or No
7. Do you have history of medical treatment for skin? Yes, or No
8. Have you ever had Acne? Yes, or No (If no, skip to question 11)
9. Do you currently have Acne? Yes, or No
10. If yes: Mild or Severe (circle based on subject’s past or current appearance)
11. Have you been prescribed an oral or topical antibiotic in the past one month? Yes, or No
12. Have you used any medicated topical antimicrobial in the last month? Yes, or No
13. If yes to 11 or 12, what type or brand? (Choose from placard P1)

Letter of chosen product(s) _

If not shown, please write down product if you can	__________________

14. Do you think you have body odor? Yes, or No (if no, skip to question 16)
15. If yes, please specify body site(s)
    a. Scalp
    b. Face
    c. Arms
    d. Arm pits
    e. Trunk (body)
    f. Legs
    g. Feet
    h. Others	_____
16. Do you routinely (> 3 times per week) use cosmetics on your face? Yes, or No
17. Do you think your scalp is healthy? Yes, or No
18. How long is your hair (Tick one)?
    a. Bald	_______
    b. Shaved	_______
    c. <5cm	_______
    d. >5cm	_______

19. Are you currently using a hair loss treatment product? Yes, or No (if no, skip to question 21)
20. If yes, what type or brand? (Choose from placard P2)

Letter of chosen product(s) _

If not shown, please write down product if you can ____________

21. Do you wear a veil, tudung, head scarf or hat?

Never

Occasionally

Daily (any)

Daily (> 2 hours/day)

22. Do you chemically dye, bleach, straighten, curl or otherwise treat your hair? Yes, or No (if no, skip to question 25)
23. If yes, when was the last time

Less than one week ago

More than one week ago

24. If yes, was it

At a salon

At home	(tick “at home” if done non-professionally, like with a friend)

25. When was the last time you washed your whole body or took a shower or bath? This morning

Yesterday evening

Yesterday Morning

More than 2 days ago

26. What did you use for cleaning when you last washed your body? (Placard P3) Water only

Soap and water

Letter from placard

If not shown, please write down the product name if you can ________

27. When was the last time you washed your face?

This morning

Yesterday evening

Yesterday Morning

More than 2 days ago

28. What product did you use for cleaning when you last washed your face? (Placard P4)

Water only

Soap and water

Letter from placard

If not shown, please write down the product name if you can ________

29. When was the last time you washed you scalp/hair? This morning

Yesterday evening

Yesterday morning

More than 2 days ago

30. What product did you use when you last washed your hair? (Placard P5) Water only

Soap and water

Letter from placard

If not shown, please write down the product name if you can ________

31. Do you routinely (>once per week) use any of the below (tick all that apply) Underarm antiperspirant

Underarm deodorant

Topical medications

Whitening or lightening product

32. Do you routinely (>once per week) use moisturizers or emollients (e.g. aqueous creams, urea creams, paraffin) Yes or No (if no, skip to question 34)
33. If yes, which products do you use (Placard P6)

Letter from placard

If not shown, please write down the product name if you can ________

34. Are you a regular swimmer?

Yes, or No (if no, skip to question 37)

35. If yes, do you swim (tick one) Once a month

Once a week	More than once a week	Almost every day

36. What best describes where you swim and the kind of water (tick one) Swimming pool – Chemically treated water

Swimming pool – Saltwater pool

In the ocean – Seawater

Freshwater – pond or lake

37. Do you routinely remove any body hair? Yes, or No (if no, stop here)
38. If yes, do you shave your underarms? Yes, or No (if no, skip to question 40)
39. If yes, when was the last time you shaved your underarm? Today

Yesterday

Less than one week ago

More than one week ago

40. Do you wax your underarms? Yes, or No (if no, skip to question 42)
41. If yes, when was the last time you waxed your underarm? Today

Yesterday

Less than one week ago

More than one week ago

42. Do you shave your bikini line? Yes, or No (if no, skip to question 44)
43. If yes, when was the last time you shaved your bikini line? Today

Yesterday

Less than one week ago

More than one week ago

44. Do you wax your bikini line? Yes, or No (if no, stop here)
45. If yes, when was the last time you waxed your bikini line? Today

Yesterday

Less than one week ago

More than one week ago

**B.) MENOPUSE QUESTIONNAIRE (For Female participants only)**

The menopause is a physiologic event, a transition in life that occurs in all women who reach midlife. These questions relate to menopause and the time period prior to menopause (known as peri-menopause). We define menopause as beginning after you have had no menstrual cycles for one year. Peri-menopause is recognized as the several years prior to menopause and generally lasts from 2-6 years. Most women recognize peri-menopause as the time at which they begin to have irregular periods. This questionnaire is intended to help you inform about your menopause experience and your general health. If you feel uncomfortable to answer menopause related questions in this form, you may seek assistance from the staff.

If you answer YES to any of these questions you may be a candidate for peri-menopause, menopause and further exploration of assessment through detailed questionnaire will be useful to inform about your experience about peri-menopause, menopause and post menopause.

1.) Do you have any irregularities in your periods (Menstrual irregularities)?

 YES	 NO (If ‘YES’, please skip Q2)

2.) At present, which statement best describes your menstrual cycle? (tick all that apply)

⧠ I have regular periods.

⧠ I have regular periods and am on oral (or prescription chemical) birth control.

⧠ I have regular periods because I am taking supplemental hormones.

3.) What was the approximate date of your last menstrual period?

---/----/------ (DD/MM/YYYY)

4.) Do you have any hot flashes, episodes of sweating?

 YES	 NO

5.) Have you experienced any of these below?

 Hot flashes, episodes of sweating

 Vaginal dryness/itching

 Vaginal Pain/Infections

 Pain during sexual intercourse

 Bladder issues

 Urinary incontinence symptoms

 Disturbed sleep

 Weight gain

 Anxiety or Irritability

 Recurrent depression (Any Mood swings)

 Migraine/headaches

 Skin crawling or Itching

 Uncontrollable stool or gas tendency

 Any pain in the joints or muscles

 General Pain attacks

 Hair loss

 Others___________________

If you answer YES or check any boxes to questions above, please proceed to answer further questions

If you answer NO or did not check any boxes to the questions above, the questionnaire ends here

6.) At what age did your menstrual cycles first become irregular? (approximate age) ­­­­­­­­­________
7.) At present which statement best describes your menstrual cycle? (tick all that apply)

(Use Placard P7 for stating medication use)

⧠	I’ve had an operation (surgery) which stopped my periods

⧠	I’ve taken medication which stopped my periods

⧠	If your periods stopped because of medication, which medication were you taking

Medication name:

Please use alphabet from Placard P7 or write your brand name of choice if not found

⧠	I’ve had chemotherapy which stopped my periods

⧠	I’ve had radiation therapy which stopped my periods

⧠	Others__________________________________________________

8.) If you no longer have menstrual periods, how old were you when your menstrual periods stopped. (Please provide us with the age at which your menstrual periods stopped regardless of why – naturally, due to surgery, medication, chemotherapy, or radiation therapy).

⧠	Younger than 20	⧠	20-29 years	⧠30-39 years

⧠	40-44 years	⧠45-49 years	⧠50-55 years

⧠	55 years and above

9.) If you are undergoing menopause or any treatments, please choose the relevant options below

⧠	Hormone therapy

⧠	Vaginal estrogen

⧠	Low-dose antidepressants

⧠	Using medications such as Clonidine

⧠	Medications to prevent or treat osteoporosis

⧠	Others ___________________________________

10.) Have you received any medical treatment, such as a hysterectomy or chemotherapy that caused or precipitated menopause? ⧠ YES	⧠ NO
11.) If YES, what treatment did you receive?

**Placards**:

P1	Q12/13: medicated topical antimicrobials

P2	Q19/20: hair loss treatment products

P3	Q26: body wash products

P4	Q28: facial wash products

P5	Q30: hair wash products

P6	Q33: emollients or moisturizers

P7	Section B: Q5: Menstrual Medications

**Definitions for skin sensitivity conditions:**

All these conditions are felt on the skin and not internal in your body

**Tingling**: Tingling are unusual prickling sensations that can happen in any part of your body.

**Burning:** A burning sensation on the skin is an irritation feeling likely the result of having come into contact with an allergen or an irritant, such as poison ivy

**Heat:** A heat sensation on skin is occasional when your body temperature increases due to environment

**Tautness:** A physical condition of skin being stretched or strained

**Itching:** Itchy skin is an uncomfortable, irritating sensation that makes you want to scratch

**Pain:** A discomfort or pain felt on skin during occasional times

**General:** A general condition in skin not feeling any skin discomfort

**Hot flashes:** Hot flashes are sudden feelings of warmth, which are usually most intense over the skin

**Instructions for CRA/Staff administration of questionnaire:**

a. Please use measuring tool as and when required to measure (Eg: Question 18 for measuring hair)
b. The subject is reminded that he or she should try to answer "yes" or "no" to each of the questions where these are the given answers.
c. Please circle your choice for Yes or No or write details as and where ever required.
d. If an answer of yes or no is required and the subject does not understand the question even when repeated, the answer is coded as "no".
e. For some of the questions an explanation may be given to the subject and instructions for these questions are provided below.
f. For questions with a photo-placard with products, please record the identifying alphabet by the photo selected or write your brand name you are using now or have used within 6 months and if the subject does not know answer is coded as No.
g. For the “Skin Sensitivity”, “on a scale of 1 to 10, how much is the sensation of (burning, tingling, tautness, etc.), then tick the box 1-10.
