## Supplementary Figure for "Large-scale skin metagenomics reveals extensive prevalence, coordination, and functional adaptation of skin microbiome dermotypes across body sites"

| | |
|---|---|
| **Demographics** | |
| Age (years) (median [IQR]) | 49.00 [40.00, 57.00] |
| Gender Male (%) | 96 (48.0) |
| Ethnicity (%) | |
| Chinese | 120 (60.0) |
| Indian | 40 (20.0) |
| Malay | 40 (20.0) |
| **Skin physiological measurements** | |
| Forearm | |
| Surface hydration (median [IQR]) | 30.02 [23.18, 39.00] |
| pH (median [IQR]) | 4.67 [4.38, 5.14] |
| TEWL[1] (median [IQR]) | 4.60 [3.67, 5.34] |
| Cheek | |
| TEWL[1] (median [IQR]) | 13.18 [11.64, 15.27] |
| Sebum (median [IQR]) | 7.00 [2.00, 19.25] |
| **Skin discomfort** | |
| Have itchy skin (%) | 59 (29.5) |
| Have sensitive skin (%) | 69 (34.5) |

[1]TEWL = Transepidermal water loss

**Supplementary Table 1: Basic characteristics of study participants.** Shown here are summary statistics for demographic data, key skin physiological parameters, and skin discomfort data for the participants enrolled in this study (n=200).

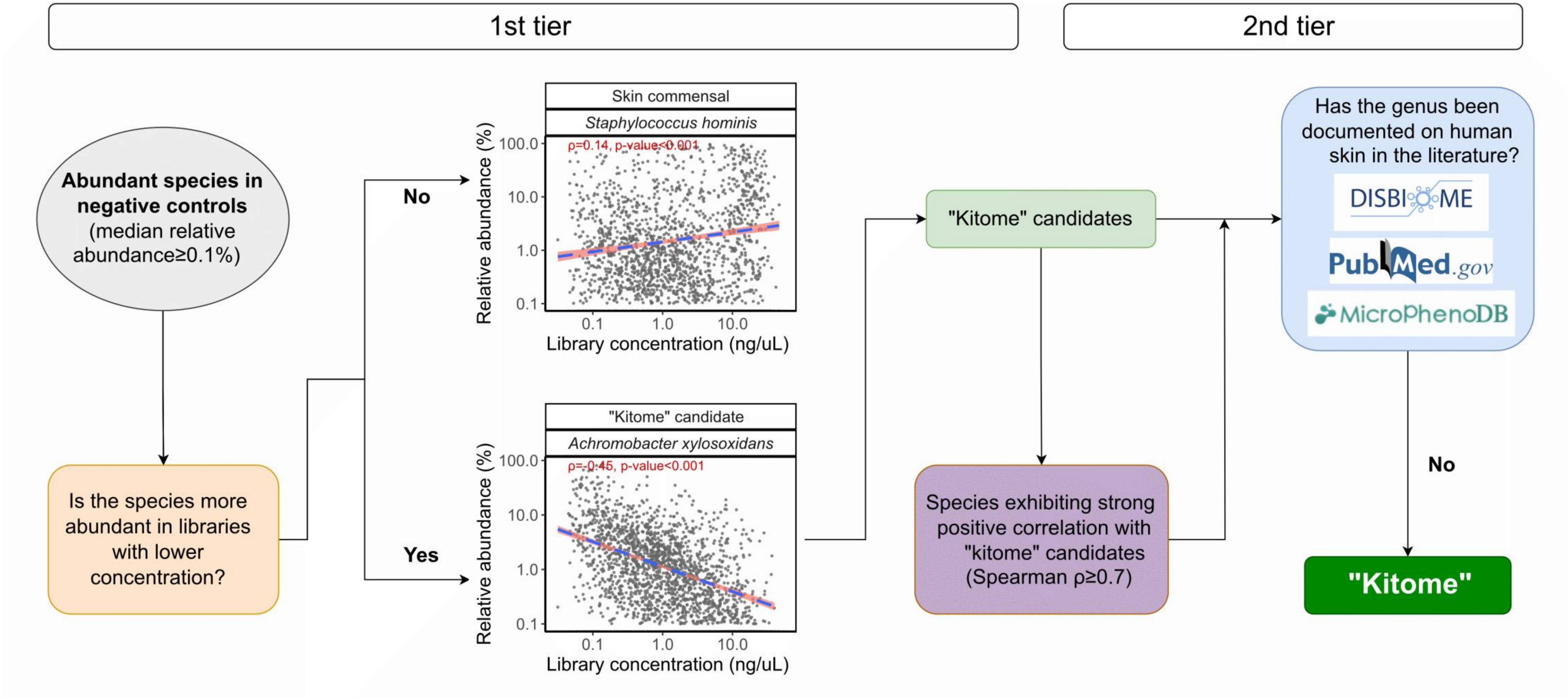


**Supplementary Figure 1: Kitome identification protocol.** Schema depicting the workflow for how potential “kitome” species are identified based on experimental data and prior literature. The first tier uses data from negative controls to identify abundant species and checks if they are more likely to be found in libraries with lower concentrations (negative Spearman correlation, $p$-value<0.05). These are designated as “kitome” candidates, and any species strongly correlated with these candidates are also considered “kitome” candidates (Spearman $\rho \geq 0.7$, $p$-value<0.05). The second tier refines the selection by restricting to genera that have not previously been reported on human skin. The identified “kitome” species were then excluded from subsequent taxonomic analyses.

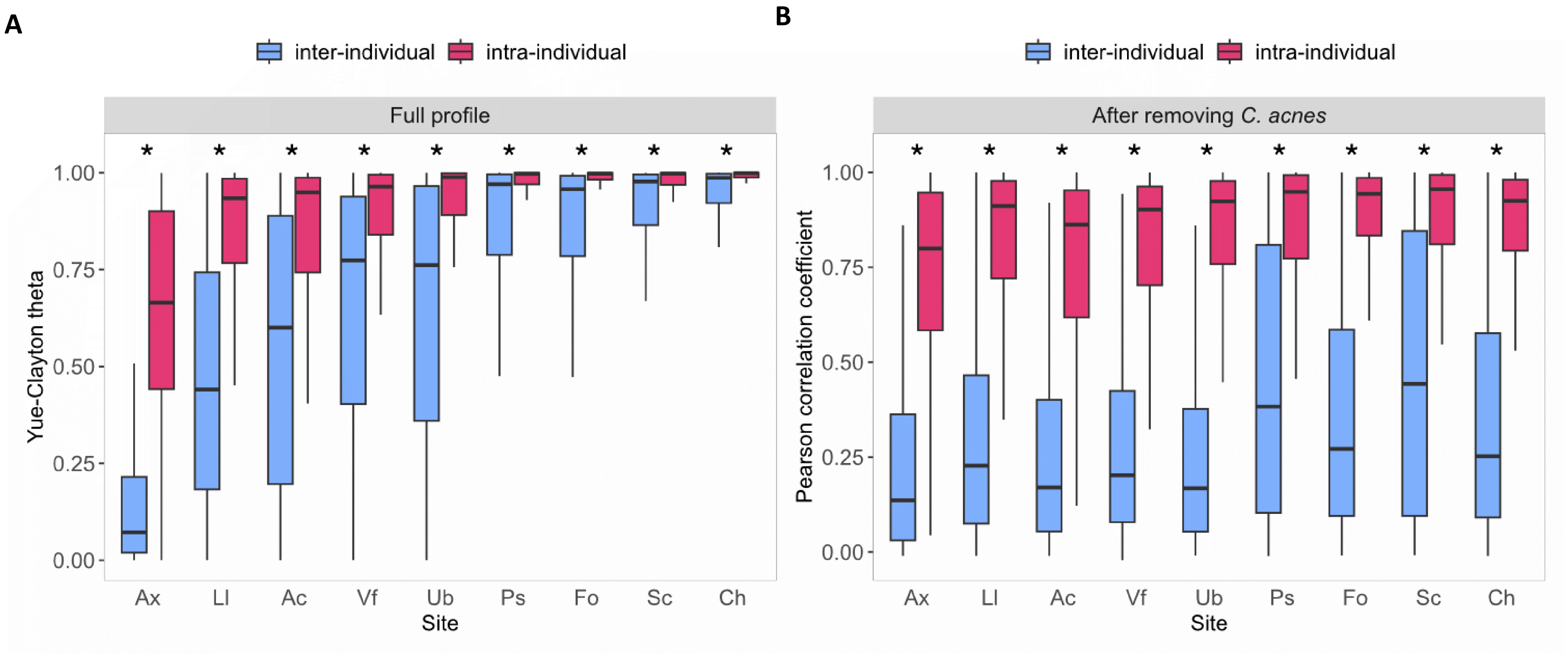


**Supplementary Figure 2: Boxplots quantifying the similarity of the skin microbiome within and between individuals across site types.** Similarity between taxonomic profiles was calculated using two indices, (**A**) Yue-Clayton theta index based on the complete microbial profile, and (**B**) Pearson correlation coefficient after removing the dominant species *Cutibacterium acnes*. All p-values were calculated using the one-sided Wilcoxon rank-sum test (* denotes p-value<0.001). Boxplot whiskers represent 1.5× interquartile range or the maximum/minimum data point within the range. x

**A**

| Site | Concordant | Discordant† |
|---|---|---|
| Ac | 177 (88.9%) | 22 (11.1%) |
| Ax | 189 (94.5%) | 11 (5.5%) |
| Ch | 162 (81.4%) | 37 (18.6%) |
| Fo | 168 (84.4%) | 31 (15.6%) |
| Ll | 174 (87.4%) | 25 (12.6%) |
| Ps | 159 (79.5%) | 41 (20.5%) |
| Sc | 167 (83.9%) | 32 (16.1%) |
| Ub | 167 (83.9%) | 32 (16.1%) |
| Vf | 175 (87.9%) | 24 (12.1%) |

† Pearson correlation coefficient < 25th percentile - 1.5*IQR

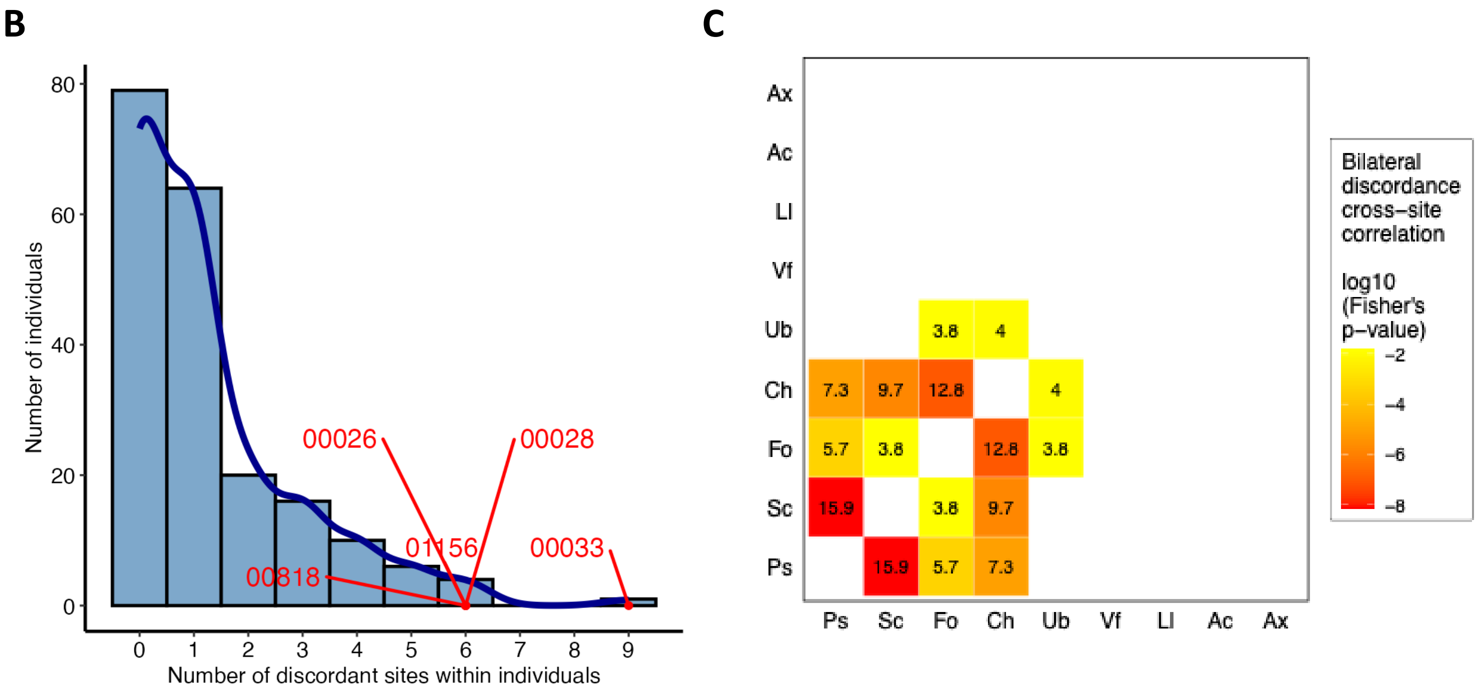


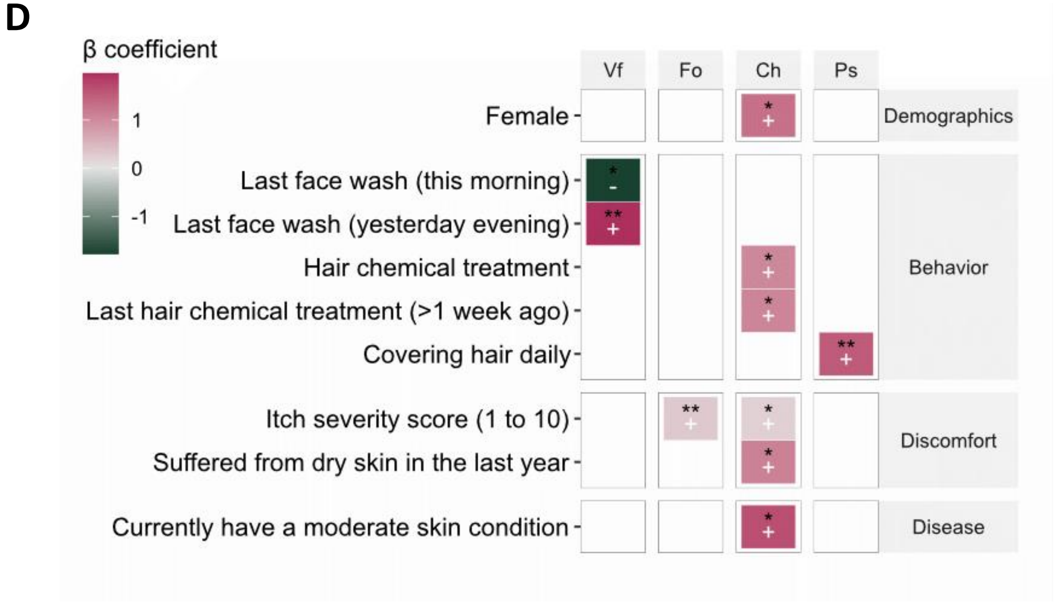


**Supplementary Figure 3: Bilateral discordance in the skin microbiome and association with phenotypic data.** (**A**) Number of discordant individuals per skin site. (**B**) Distribution of the number of discordant sites within individuals, with significant subjects (BH-adjusted binomial test p-value<0.05) highlighted in red. (**C**) The color gradient denotes $\log_{10}$-transformed BH-adjusted p-values from Fisher's exact test, and odds ratios are annotated in the boxes. (**D**) Significant associations between bilateral discordance and various phenotypes (*, ** and *** denote BH-adjusted p-value<0.1, 0.05 and 0.01, respectively).

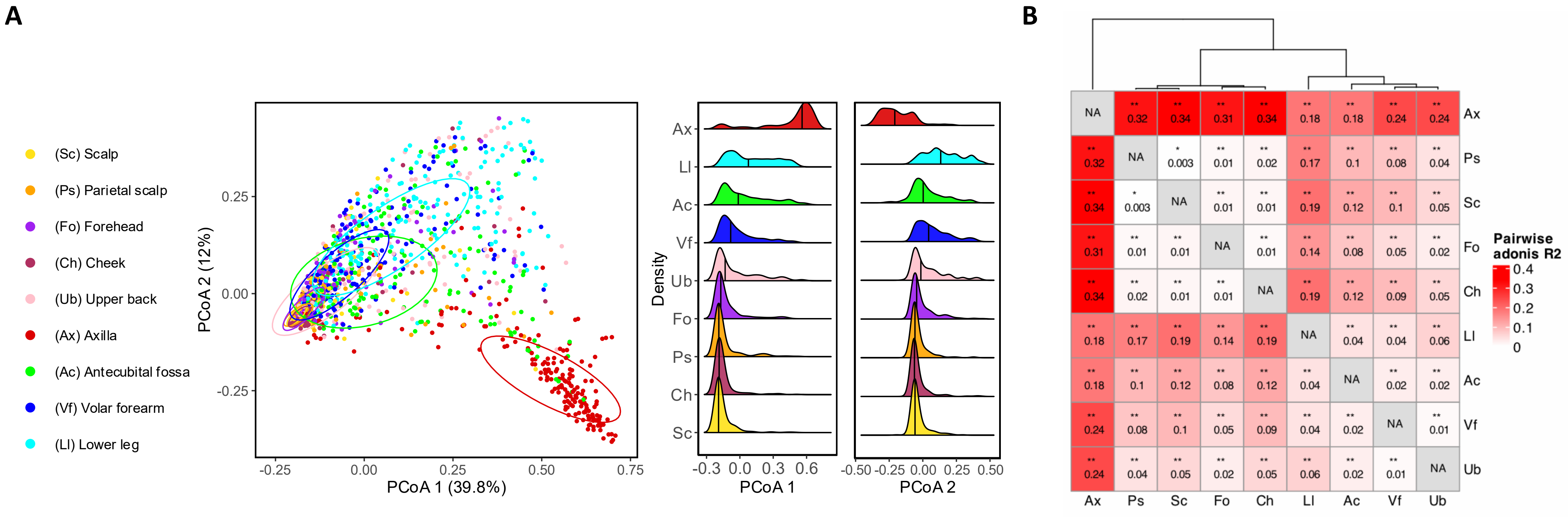


**Supplementary Figure 4: Human skin microbial community differences across sites.** (**A**) Principal coordinates analysis (PCoA) plot based on Bray-Curtis dissimilarity between left-and-right averaged site-specific taxonomic profiles (species level) of all 200 subjects with site-specific color coding. Adjacent density plots delineate sample distributions along the first two PCoAs across sites. (**B**) Heatmap of significant pairwise PERMANOVA (adonis) $R^2$ between sites, calculated based on Bray-Curtis dissimilarity, with dendrogram clustering sites with similar microbial composition (*, ** and *** denote BH-adjusted p-value<0.05, 0.01 and 0.001, respectively).

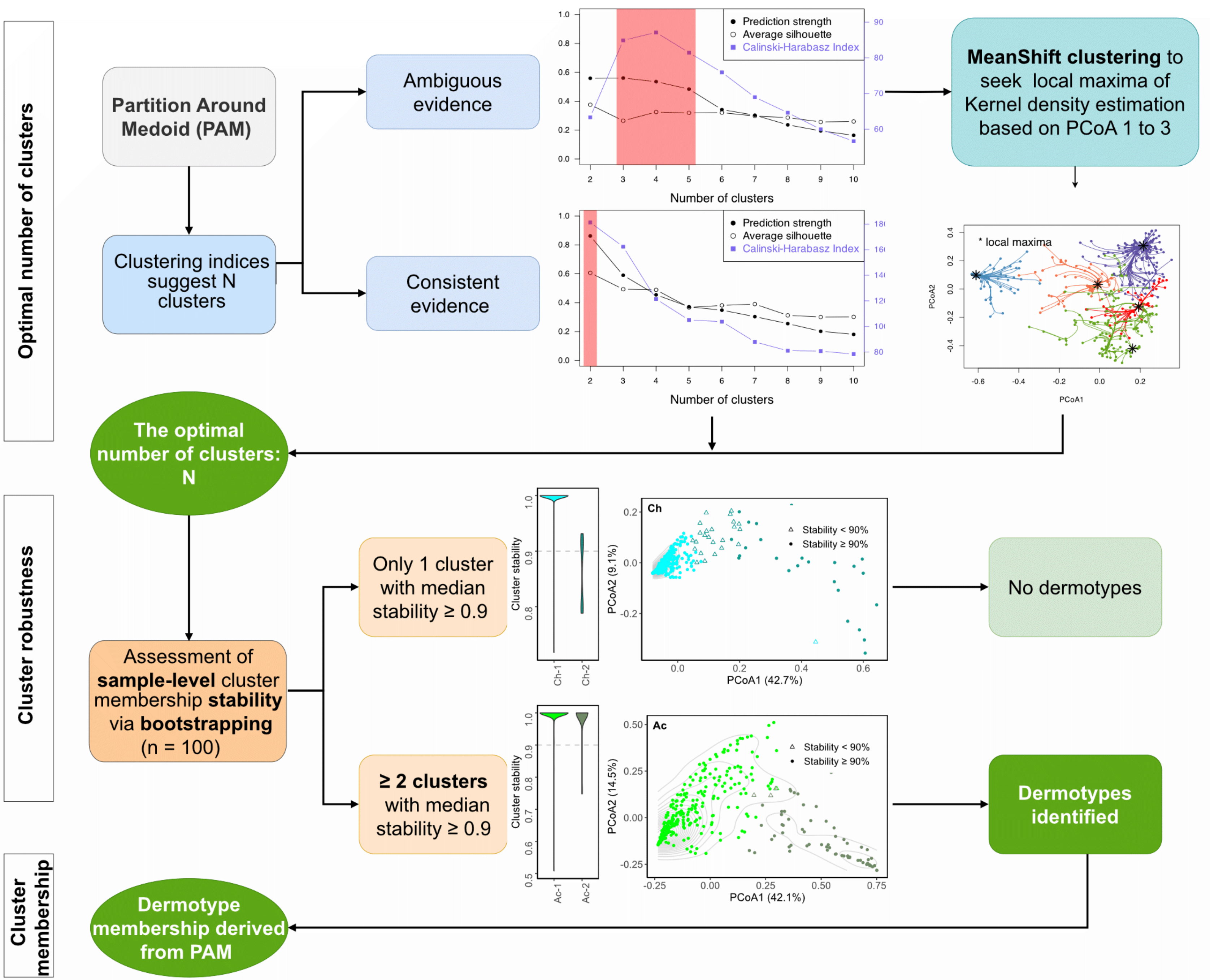


**Supplementary Figure 5: Dermotype identification protocol.** Schema illustrating the dermotype identification process and stability assessment. Firstly, Partitioning Around Medoid (PAM) clustering was performed using the Bray-Curtis dissimilarity for each site; the optimal number of clusters was determined using key indices (prediction strength, silhouette index, and Calinski-Harabasz index). In cases of ambiguity (i.e., disagreement across various indices), we employed Mean-Shift clustering (bandwidths determined by cross-validation) to search the local maxima of kernel density estimated in PCoA space (PCoA 1-3). Next, we assessed each sample’s cluster membership stability via bootstrapping (n=100). Clusters where at least 50% of samples consistently clustered together in over 90% of the iterations were considered robust (median stability≥0.9). Only sites with at least 2 robust clusters were concluded to contain dermotypes and considered for subsequent analyses. Finally, we relied on PAM to infer cluster memberships for robust dermotypes.

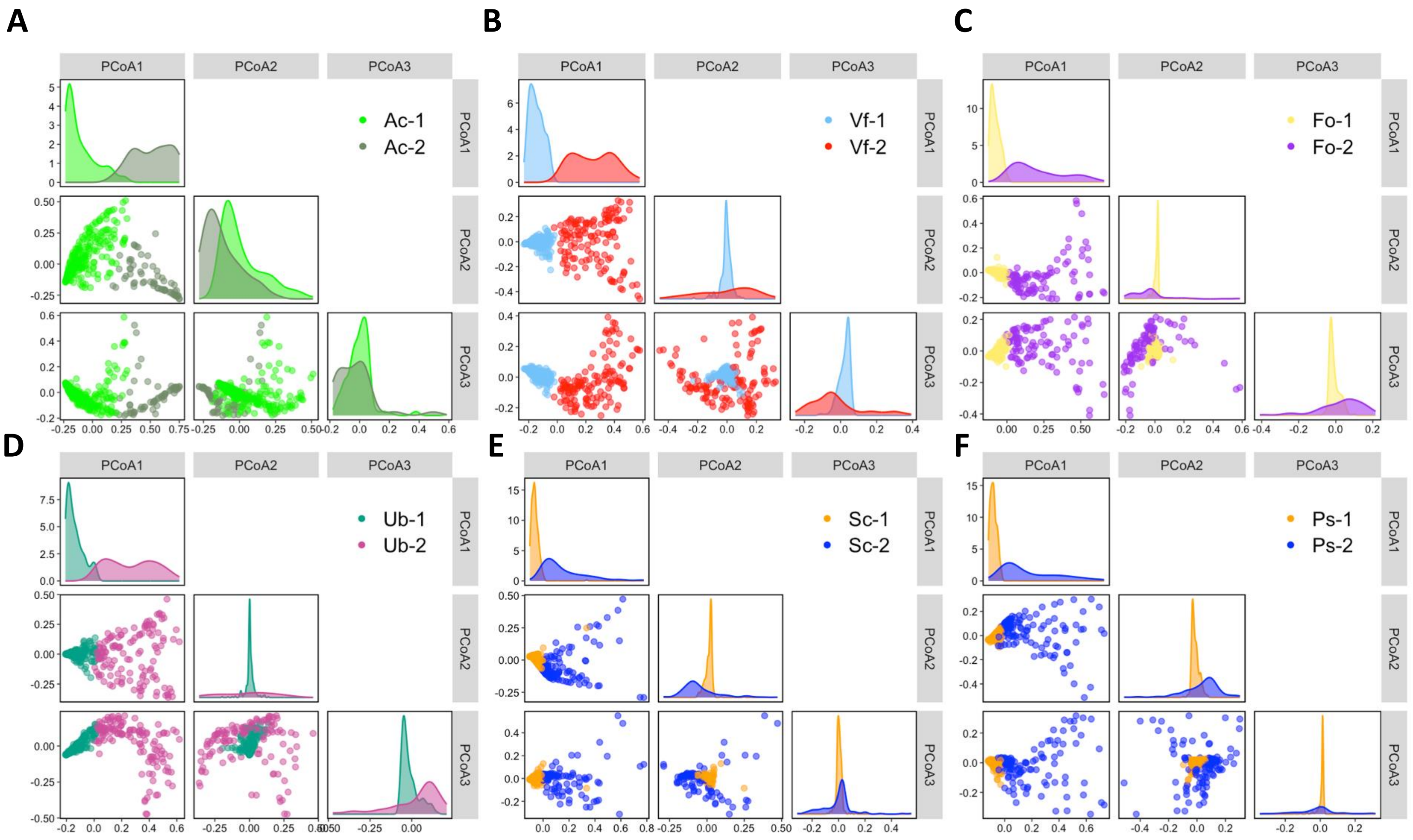


**Supplementary Figure 6: Dermotypes identified at various sites**. Site-specific principal coordinates analysis (PCoA) plots based on Bray-Curtis dissimilarity, and density plots delineating sample distributions along the first three PCoAs, reveal two dermotypes found at (**A**) elbow crease, (**B**) volar forearm, (**C**) forehead, (**D**) upper back, (**E**) scalp, and (**F**) parietal scalp, respectively.

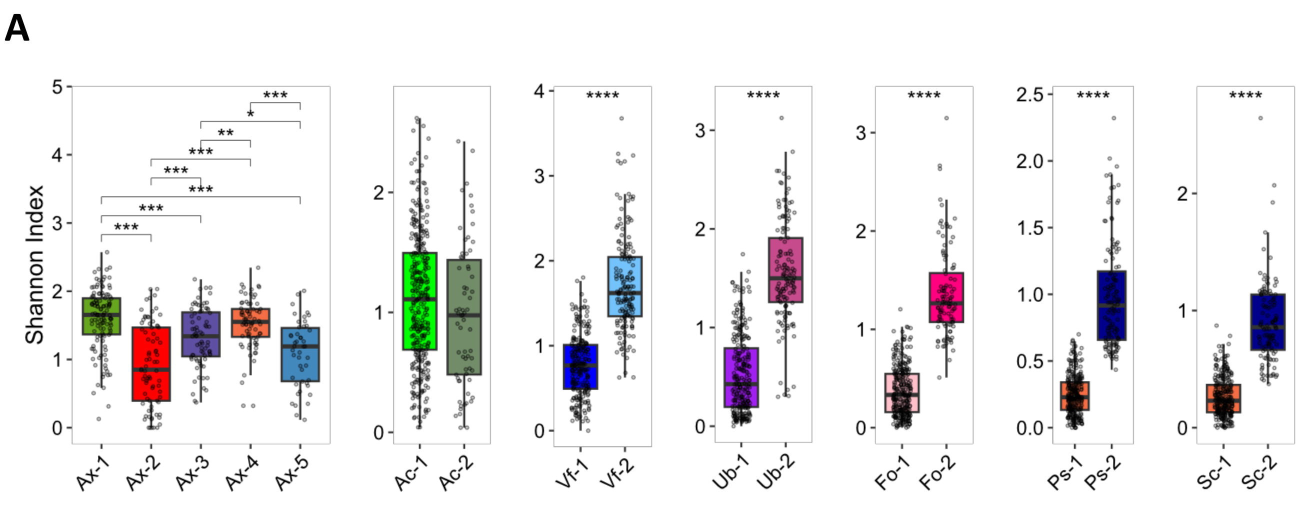


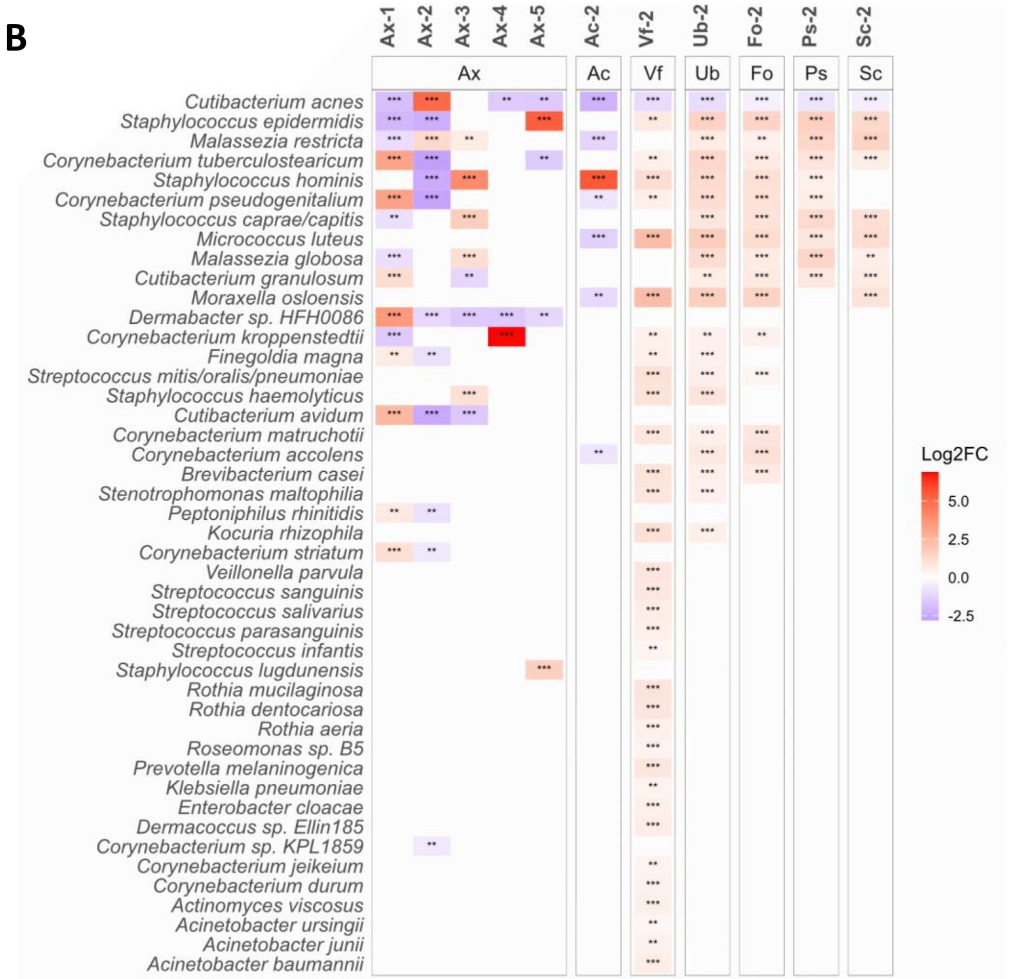


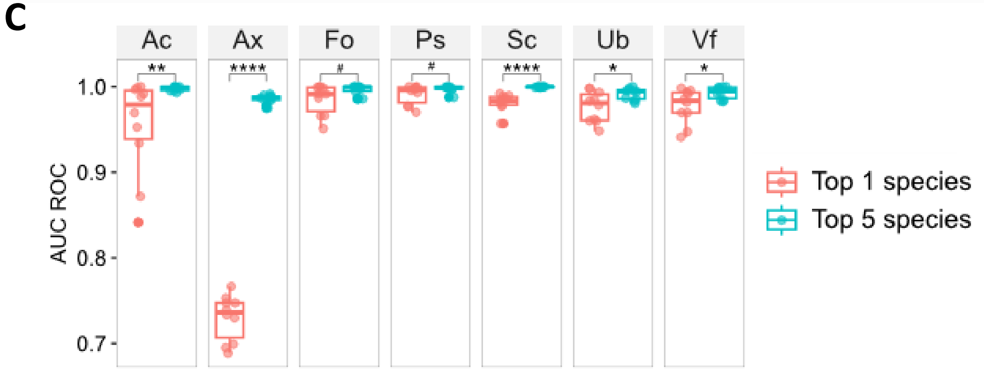


**Supplementary Figure 7: Diversity and taxonomic differences between dermotypes**. (**A**) Boxplots depicting alpha diversity comparisons between dermotypes. P-values were calculated using the two-sided Wilcoxon rank-sum test and BH-adjusted. (**B**) Differential abundant species between dermotypes, identified using one-versus-the-rest approach (prevalence≥10%, BH-adjusted p-value<0.05). Low stability samples (stability<90%) were excluded. $Log_2$ fold changes were calculated with MaAsLin2 generalized linear model with the following formula: $Species \sim Dermotype + Age + Gender + Ethnicity + (1|Subject)$. (**C**) Boxplots depicting AUC-ROC values for dermotype classification using random forest models trained on top five versus one discriminatory species. P-values were calculated using the one-sided Wilcoxon rank-sum test and BH-adjusted. Statistical significance: $^{\#}p<0.1$, $^{*}p<0.05$, $^{**}p<0.01$, $^{***}p<0.001$, $^{****}p<0.0001$.

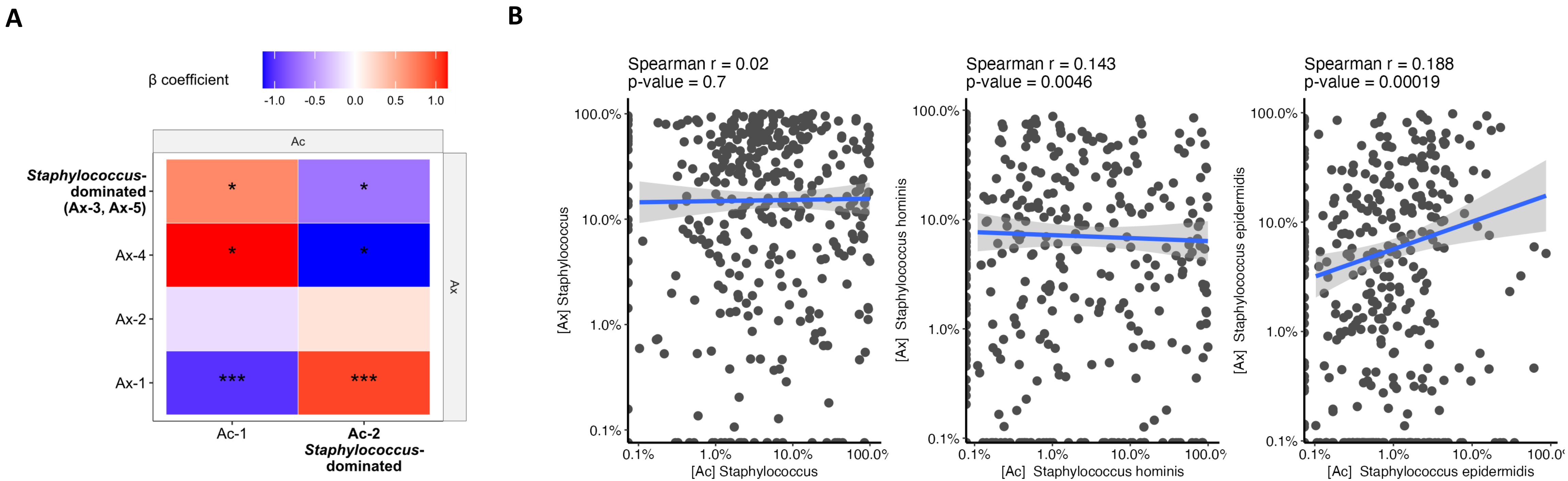


**Supplementary Figure 8: Dermotype correlations between elbow crease (Ac) and axilla (Ax).** (**A**) Heatmap displaying correlations between Ac and Ax dermotypes, with color gradients representing the estimated association strength, derived using a logistic GLM (∗, ∗∗ and ∗∗∗ denote p−value<0.05, 0.01 and 0.001, respectively). (**B**) Correlation analysis of *Staphylococcus* abundance between Ac and Ax microbiomes.

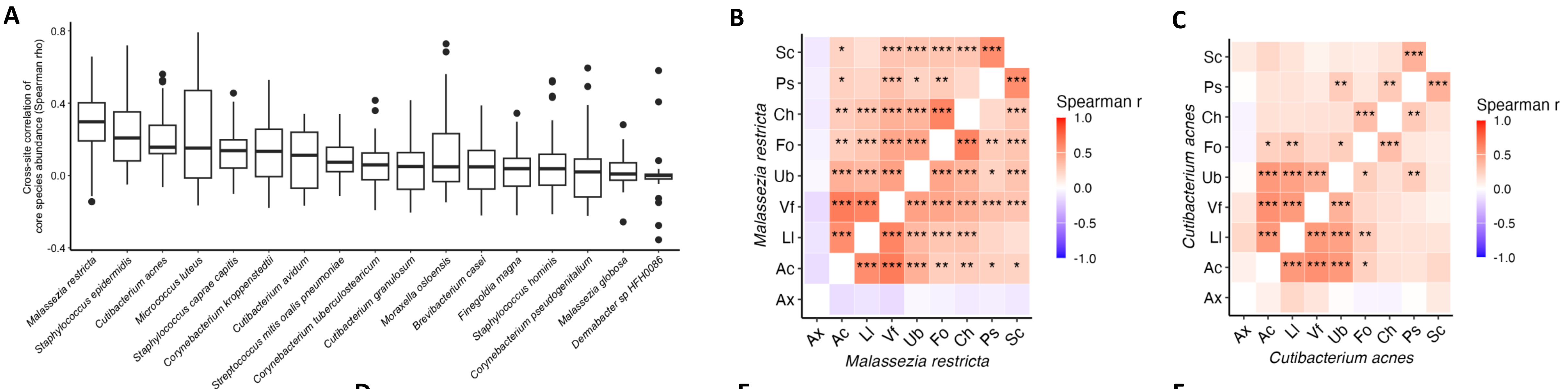


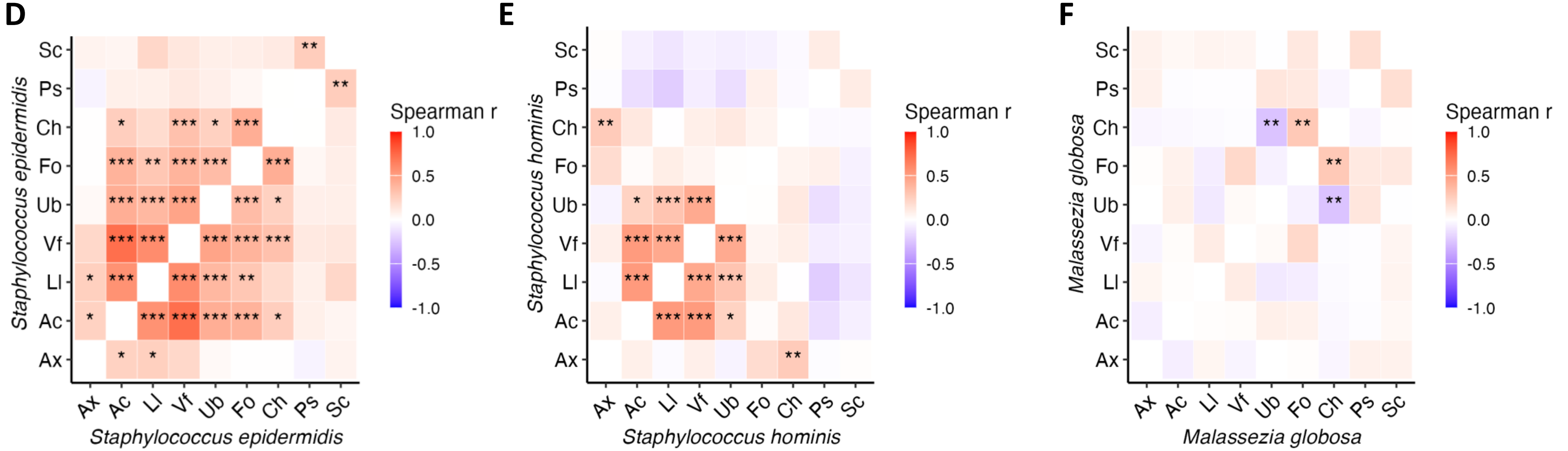


**Supplementary Figure 9: Intra-individual cross-site correlations of core species abundance.** Taxonomic profiles from bilateral sites were averaged before applying centered log-ratio (CLR) transformation. Spearman correlation was used to assess correlation strength. (**A**) Boxplots that show the distribution of core species correlation strengths across all pairs of skin sites. (**B–F**) Heatmaps illustrating cross-site correlation patterns for selected core species, with color gradients representing correlation strength (∗, ∗∗ and ∗∗∗ denote BH−adjusted p−value< 0.05, 0.01 and 0.001, respectively).

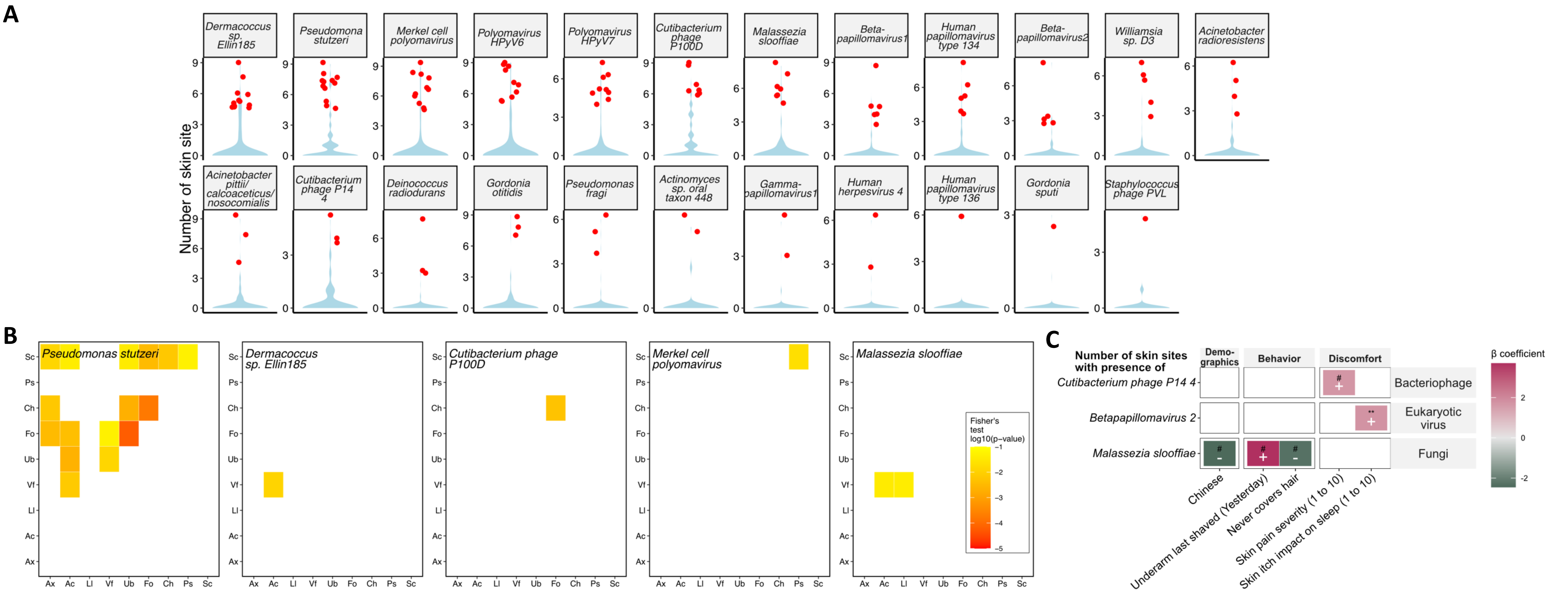


**Supplementary Figure 10: Enrichment of ancillary taxa across multiple skin sites of many individuals.** (**A**) Violin plot showing the distribution of the number of skin sites each ancillary taxa is present in within a given individual. Statistical significance was assessed using a Poisson binomial test, where the expected probability for each trial was the prevalence of the taxon across the population and per site. Red dots indicate individuals with significant enrichment of ancillary species across multiple body sites, defined as a BH-adjusted p-value<0.05. (**B**) Correlation analysis of ancillary species co-occurrence across skin sites. Analysis was restricted to species present in at least 10 individuals per site. The color gradient represents $\log_{10}$-transformed BH-adjusted p-values from Fisher's exact test. (**C**) Heatmap illustrating associations between ancillary species multi-site carriage and host phenotypes. Ancillary species detected in fewer than 10 individuals or with fewer than 2 individuals carrying the species at multiple sites were excluded. Color gradients represent estimated beta coefficients from linear models. Statistical significance: #p<0.1 , *p<0.05, **p<0.01.

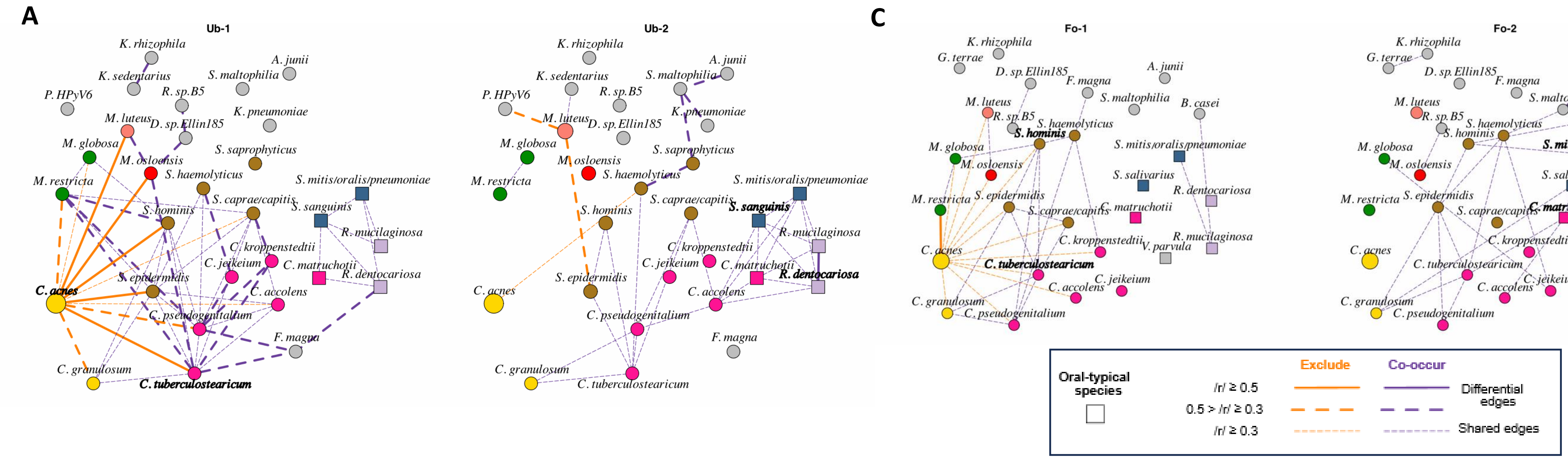


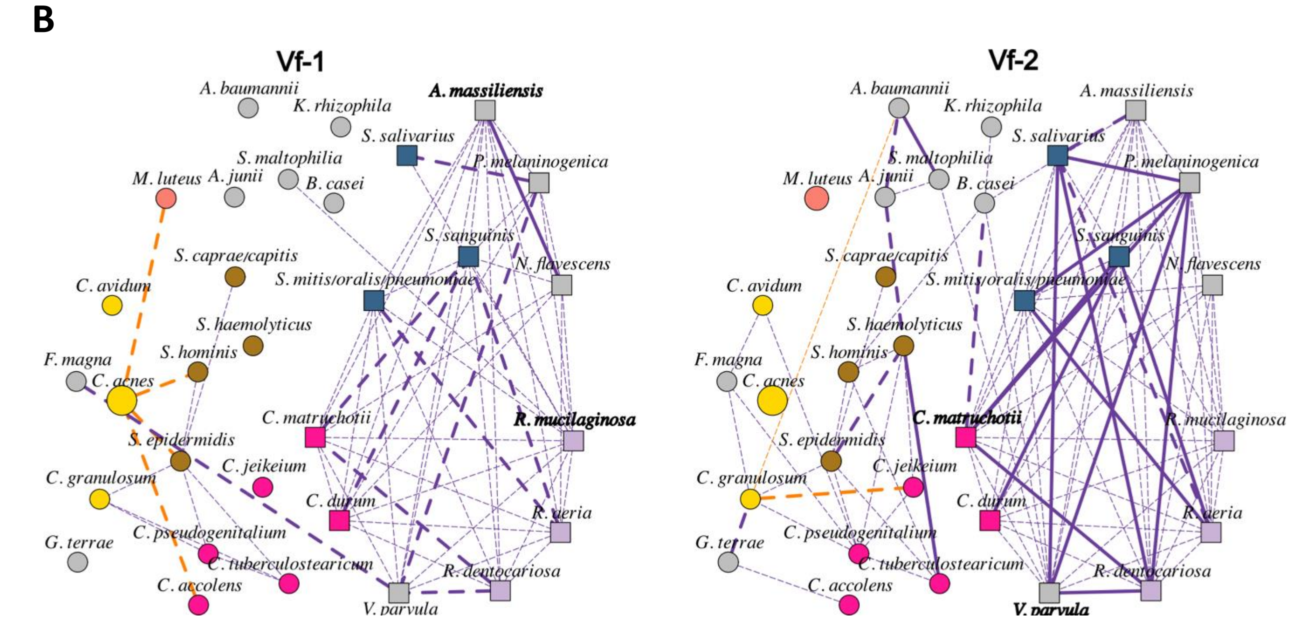


**Supplementary Figure 11: Co-abundance networks across dermotypes for selected sites, including (A) upper back, (B) volar forearm, and (C) forehead.** Each network was constructed with taxonomic profiles of corresponding samples. Nodes in the networks represent microbial species, and edges indicate significant compositionally-corrected Spearman correlations between the relative abundances of species (CCREPE, |ρ|≥0.3, p-value<0.05). Purple and orange edges denote positive and negative correlations, respectively. Solid and thick dashed lines represent significant differential edges between the two networks, while thin dashed lines represent shared edges. Bolded species are hubs identified in each network. Full results, including Spearman correlation values and corresponding p-values for each species pair in each dermotype, are available in **Supplementary File 7**.

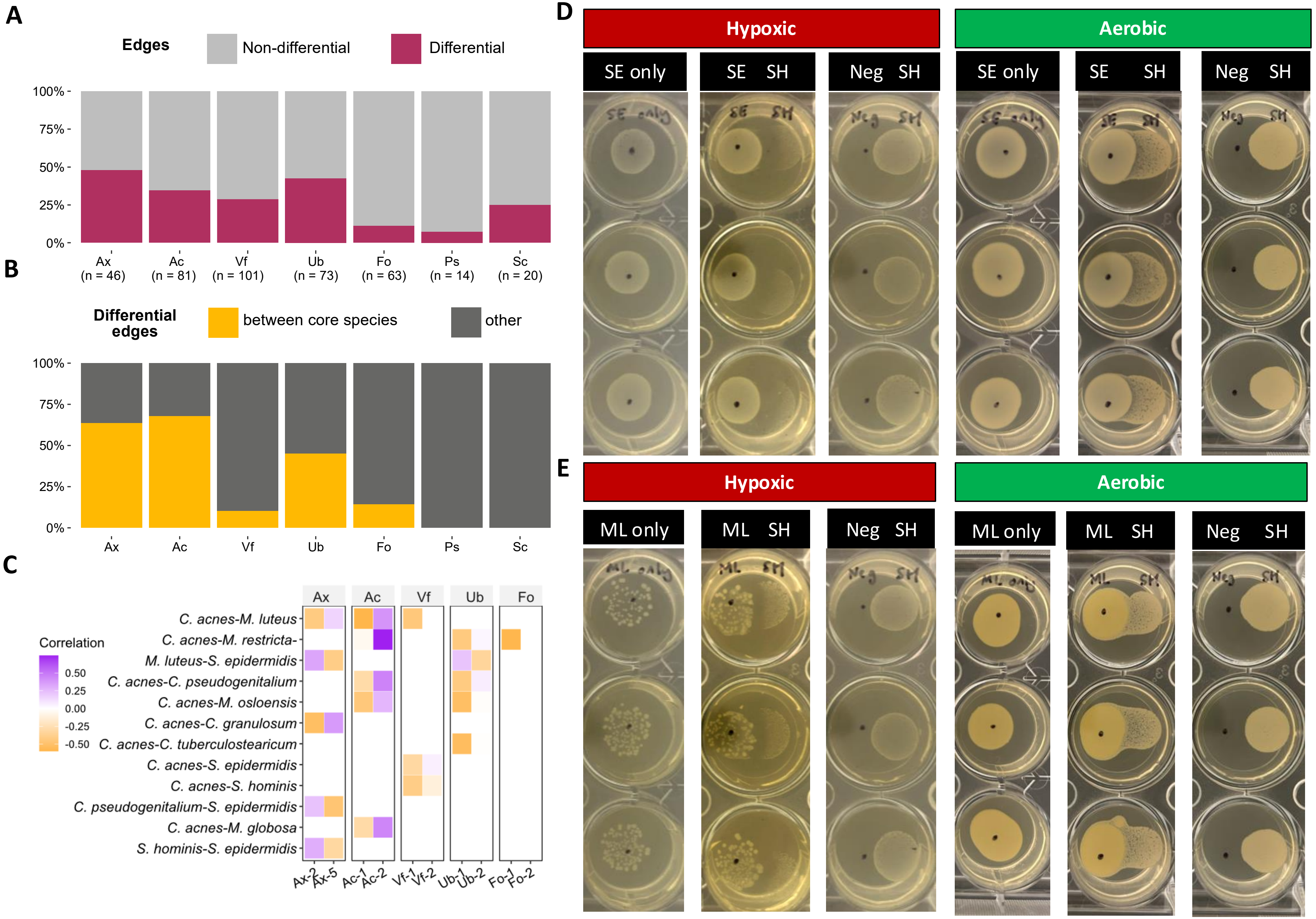


**Supplementary Figure 12: Co-abundance networks and microbial interactions under different conditions.** (**A**) Proportion of network edges that are differentially present in a site between dermotypes. (**B**) Proportion of differentially present edges that connect core species in the dermotypes for a site. (**C**) Top five differential co-abundance relationships between core species at each dermotype, and their corresponding correlation coefficients. (**D-E**) Full set of agar spot antagonism assay results showing *S. epidermidis* (**D**) and *M. luteus* (**E**) interactions with *S. hominis* under hypoxic and aerobic conditions.

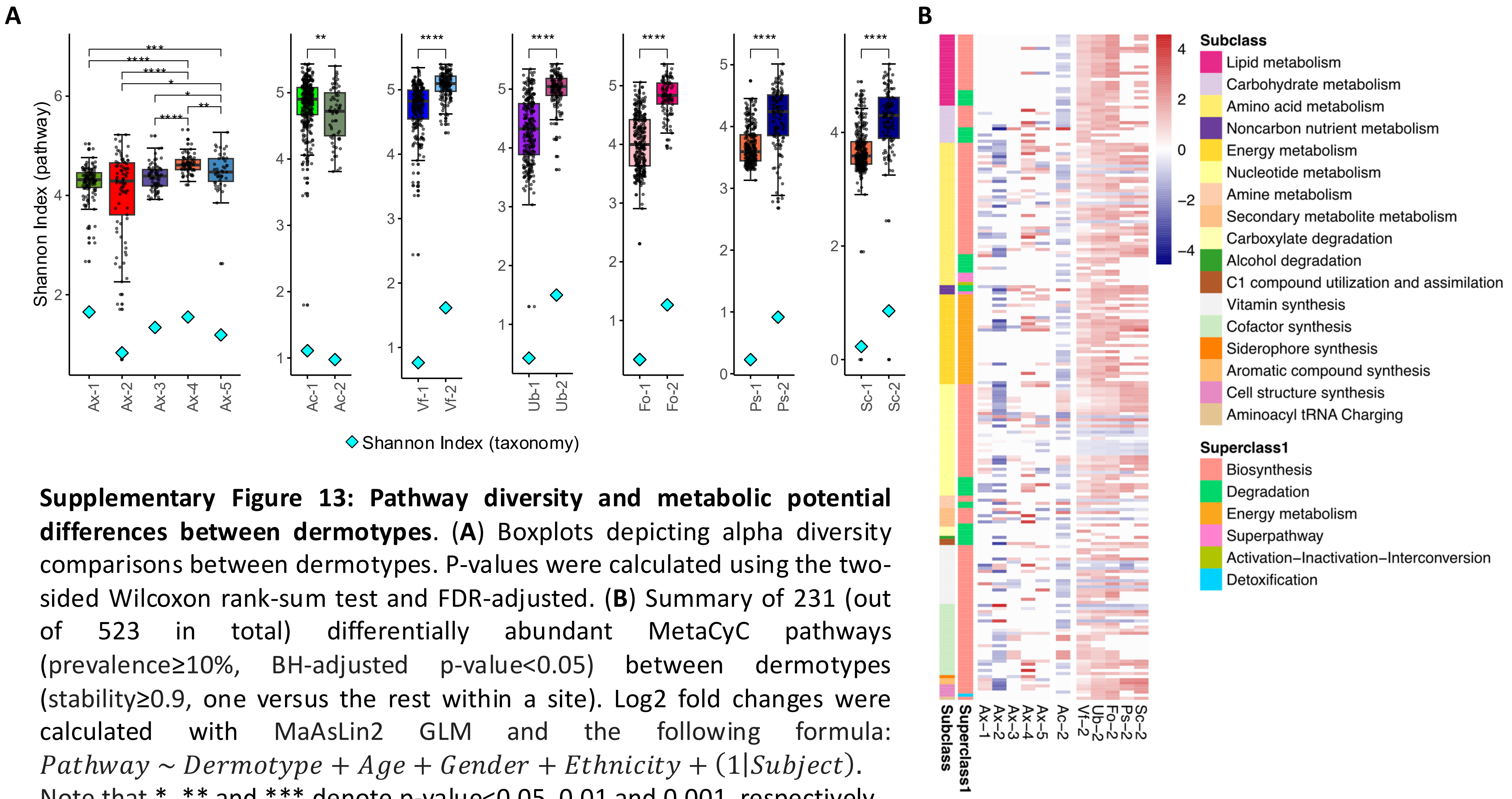


**Supplementary Figure 13: Pathway diversity and metabolic potential differences between dermotypes**. (**A**) Boxplots depicting alpha diversity comparisons between dermotypes. P-values were calculated using the two-sided Wilcoxon rank-sum test and FDR-adjusted. (**B**) Summary of 231 (out of 523 in total) differentially abundant MetaCyC pathways (prevalence≥10%, BH-adjusted p-value<0.05) between dermotypes (stability≥0.9, one versus the rest within a site). Log2 fold changes were calculated with MaAsLin2 GLM and the following formula: $Pathway \sim Dermotype + Age + Gender + Ethnicity + (1|Subject)$. Note that *, ** and *** denote p-value<0.05, 0.01 and 0.001, respectively.

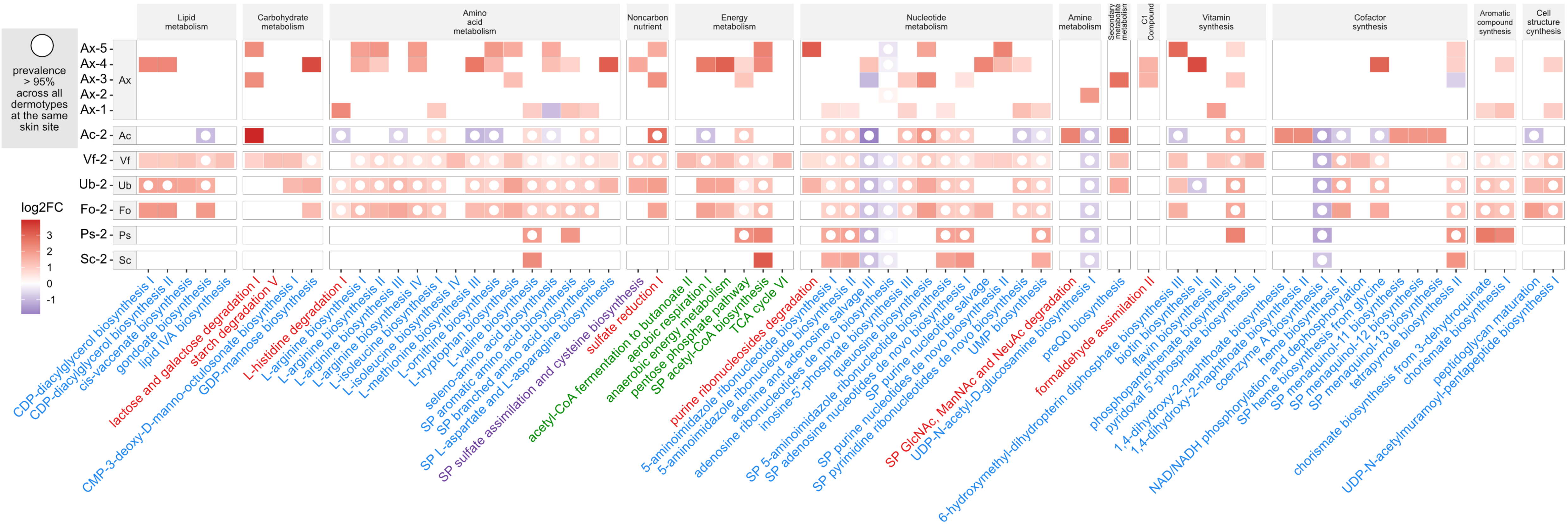


**Supplementary Figure 14: Heatmap showing the top 10 differentially abundant MetaCyC pathways across dermotypes.** A pathway is included if it is prevalent in over 95% of samples in at least one dermotype and ranks among the top five based on |log2FC|. Hollow circles indicate pathways present in over 95% of samples across all dermotypes at the same skin site.

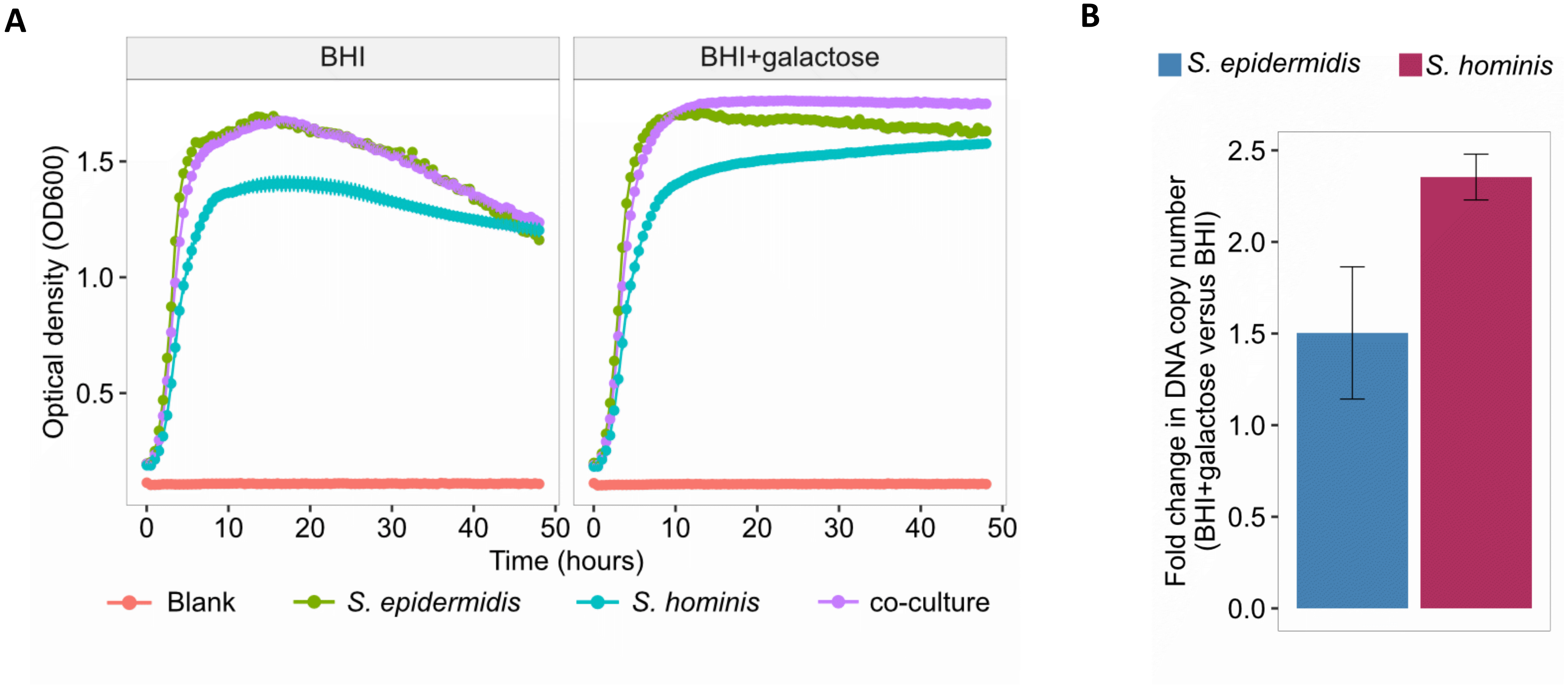


**Supplementary Figure 15: Effect of galactose supplementation on the growth of *Staphylococcus* species.** (**A**) Growth curves for *S. epidermidis*, *S. hominis*, and their co-culture in liquid medium (BHI) without and with 5% galactose, measured by optical density at 600nm ($OD_{600}$). (**B**) Bar plots showing the log2 fold change in DNA copy number of *S. hominis* and *S. epidermidis*, as quantified by qPCR, under co-culture conditions in BHI supplemented with 5% galactose versus BHI medium (48h).

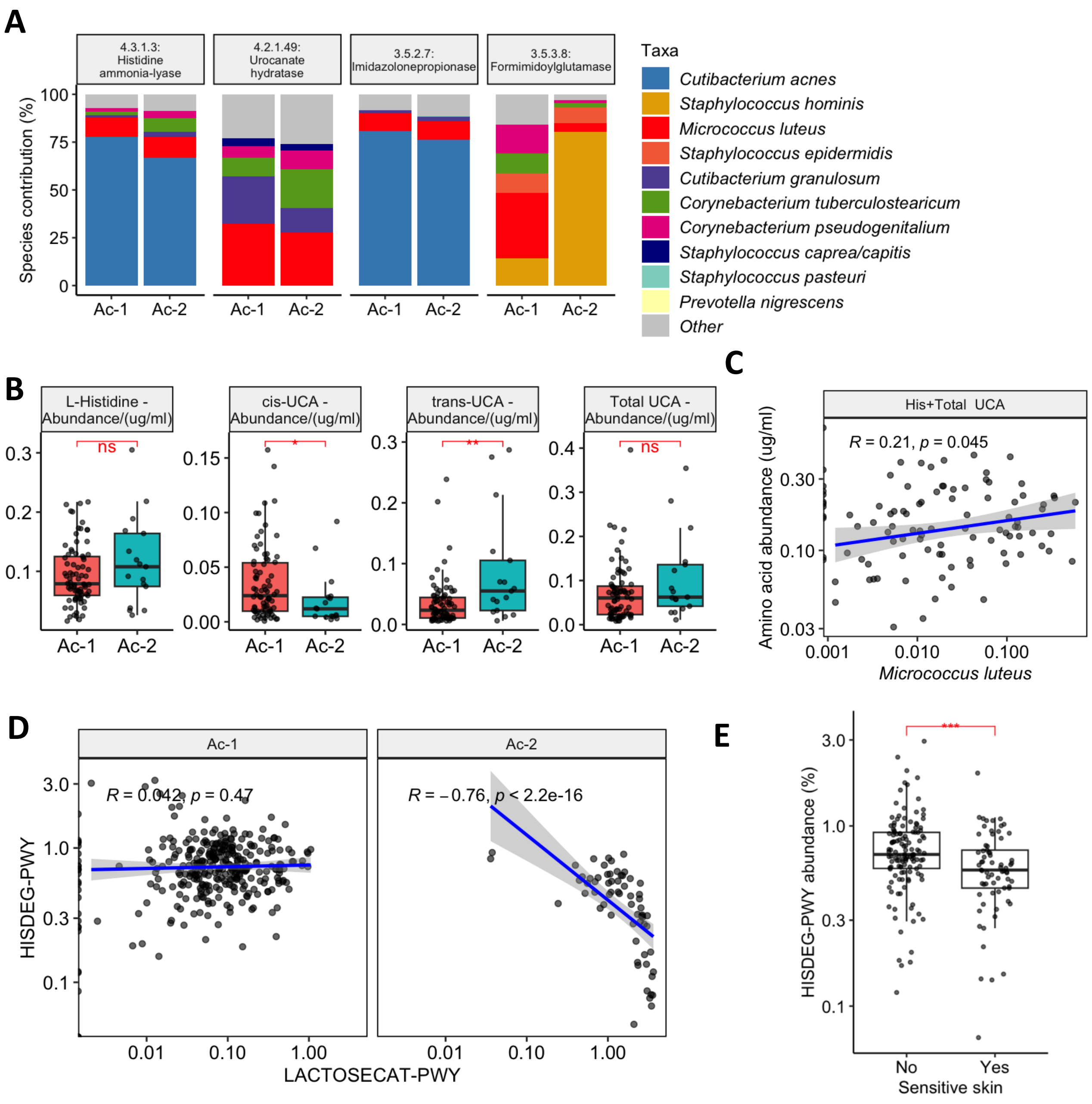


**Supplementary Figure 16: Multi-omic analysis of *Micrococcus luteus* and histidine-related amino acids in the elbow crease.** (**A**) Stacked barcharts depicting HUMAnN2-derived species contributions to enzymes involved in consecutive steps of the L-histidine degradation pathway I (HISDEG-PWY). (**B**) Boxplots depicting levels of L-histidine and its derivatives in the elbow crease of Ac-1 and Ac-2 subjects. Data was from an independent cohort (n=100). (**C**) Scatterplot showing positive correlation between *M. luteus* abundance and levels of L-histidine and its derivatives in the elbow crease. Spearman rank correlation coefficients and p-values are indicated. (**D**) Scatterplot showing the correlation between abundances of the L-histidine degradation I pathway and lactose and galactose degradation I pathway in Ac-1 and Ac-2 subjects. (**E**) Boxplots depicting the differences in abundance of the L-histidine degradation I pathway in subjects with and without sensitive skin. UCA: urocanic acid. Statistical significance: *p<0.05, **p<0.01, ***p<0.001, ****p<0.0001.

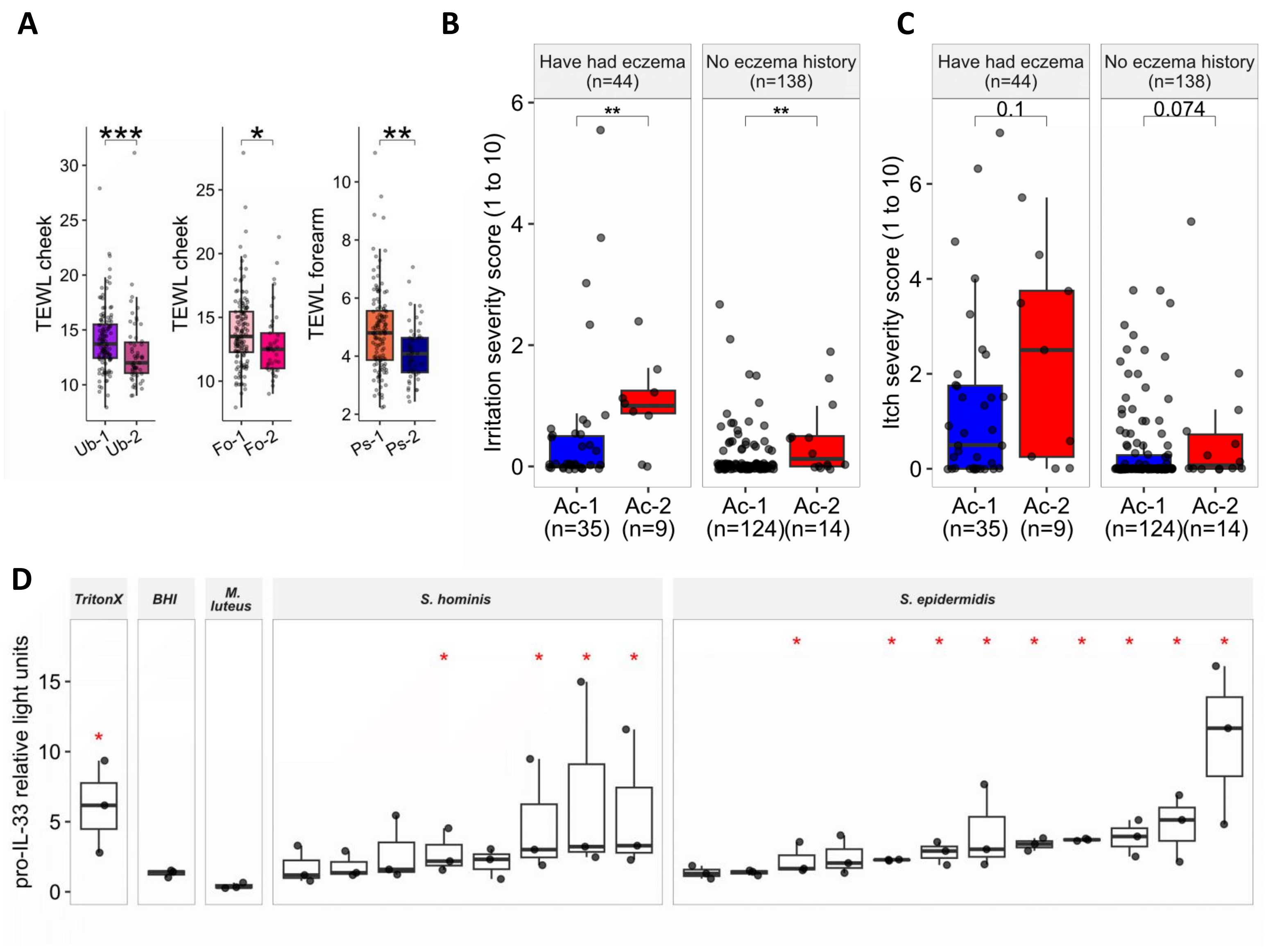


**Supplementary Figure 17: Dermotype-associated variations in skin physiology, irritation, itch, and an itch-related immune marker.** (**A**) Boxplots showing significant differences in cheek and forearm trans-epidermal water loss (TEWL) across upper back (Ub), forehead (Fo), and parietal scalp (Ps) dermotypes. P-values were calculated using the one-sided Wilcoxon test. (**B-C**) Boxplots depicting skin irritation severity (B) and itch severity (C) across Ac dermotypes, stratified by eczema history. P-values were calculated using the one-sided Wilcoxon test. (**D**) Boxplots showing levels of pro-IL-33 relative light unit (RLU) in human keratinocytes (measured by HiBiT) following exposure to supernatants from *Micrococcus luteus*, *Staphylococcus hominis*, and *Staphylococcus epidermidis* strains. Data was from three biological repeats, each compromising three technical replicates. One-sided Wilcoxon tests were used to compare pro-IL-33 RLU levels to negative controls (treated with BHI and treated with *M. luteus* supernatant) and BH-adjusted p-values are shown. Statistical significance: *p<0.05, **p<0.01, ***p<0.001.

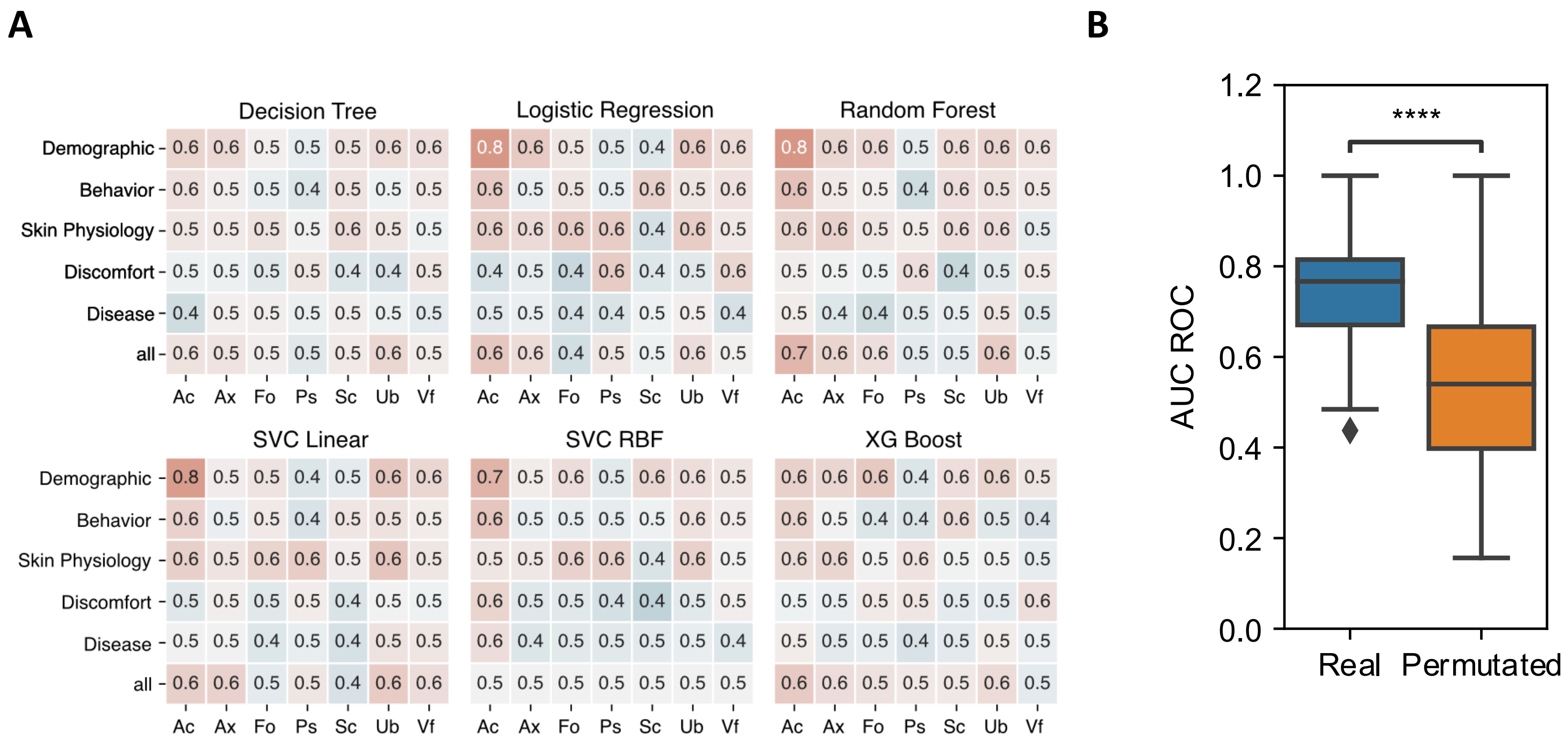


**Supplementary Figure 18: Dermotype classification with host attributes as features.** (**A**) Average ROC-AUC values of six different dermotype classifiers across seven skin sites, trained using distinct host attribute domains and all domains combined. (**B**) Distribution of AUC-ROC values for a random forest Ac dermotype classifier, trained on ethnicity attribute, using either real or permutated labels, with five iterations. The p-value was calculated using one-sided Wilcoxon rank-sum test (**** denotes p-value<0.0001).

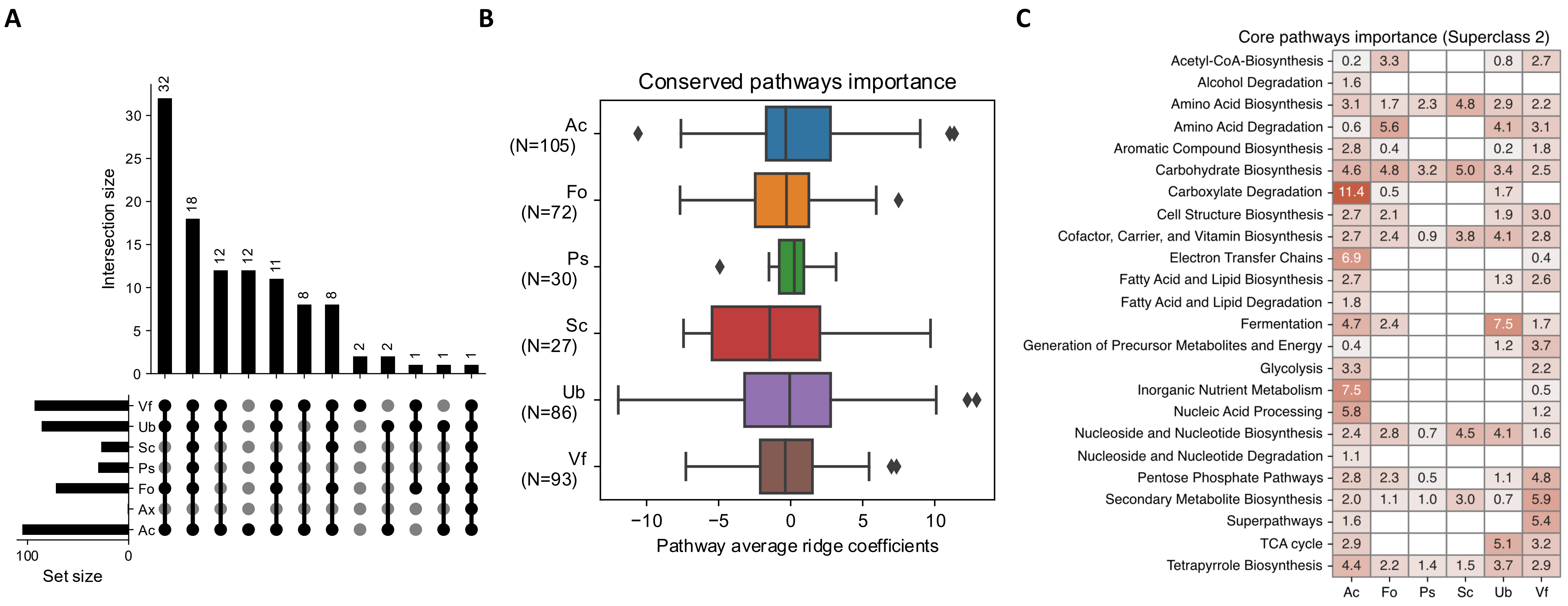


**Supplementary Figure 19: Dermotype classifiers trained on key metabolic pathways.** (**A**) Upset graph representing the overlap of conserved pathways across sites. Ax was excluded from further analysis due to a low number of conserved pathways. (**B**) Average ridge coefficients for all conserved pathways for each site. (**C**) Average feature importance (mean of absolute values of ridge coefficients) for each conserved pathway superclass across skin sites.

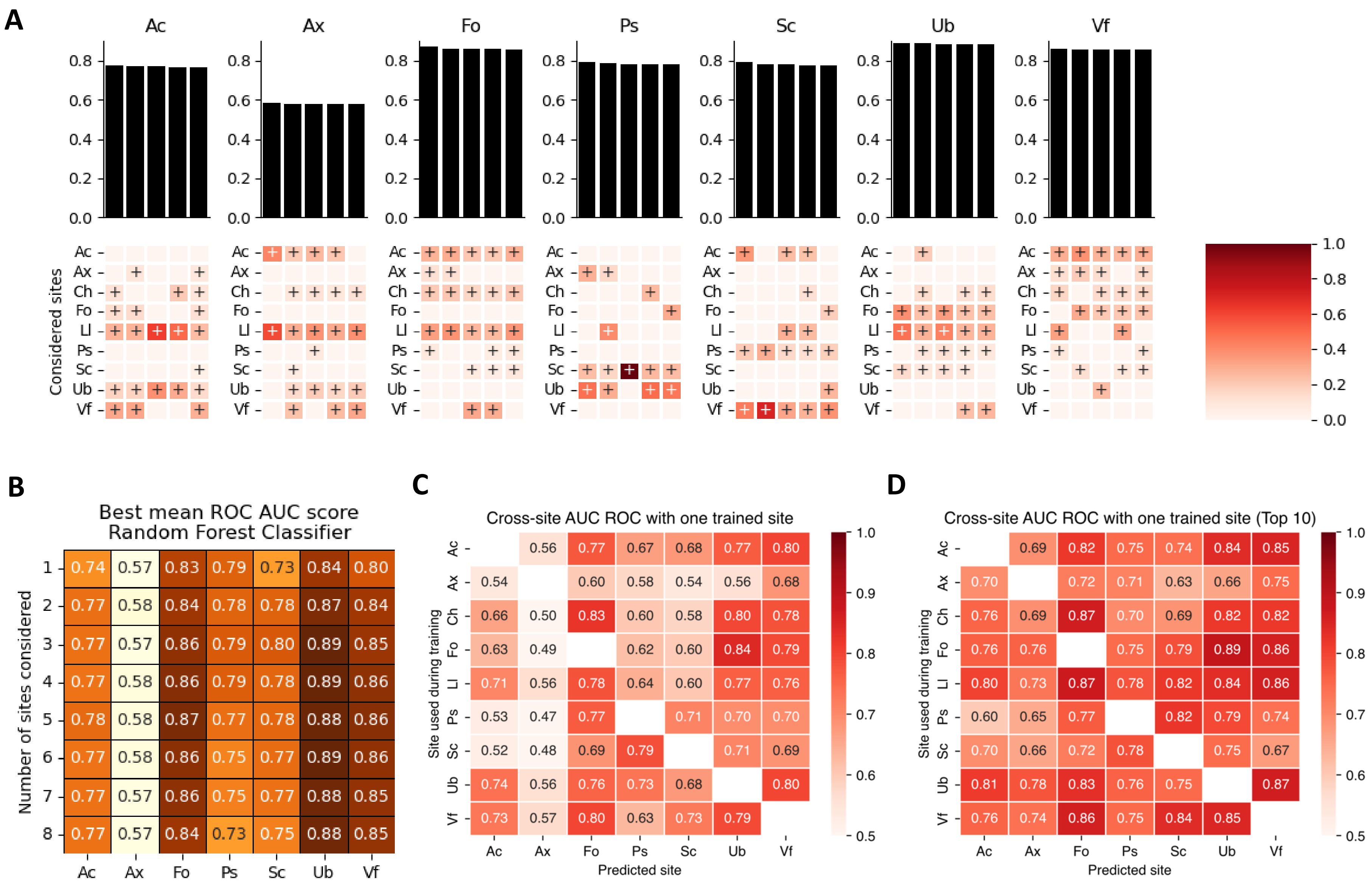


**Supplementary Figure 20: Random forest dermotype classifiers trained on cross-site taxonomic information.** (**A**) Top 5 combinations of cross-sites features for each skin site. Each column represent a training combination, with AUC-ROC values displayed in the top bar plot. The bottom heatmap indicates the cumulative importance of features from their corresponding sites, with a "+" symbol highlighting the sites included in each training combination. (**B**) AUC-ROC values obtained with an increasing number of sites included in the training. (**C-D**) Dermotype classification using cross-site taxonomic information for a single site, with models trained on (**C**) all species abundances or (**D**) just the top 10 most important species for each pair.

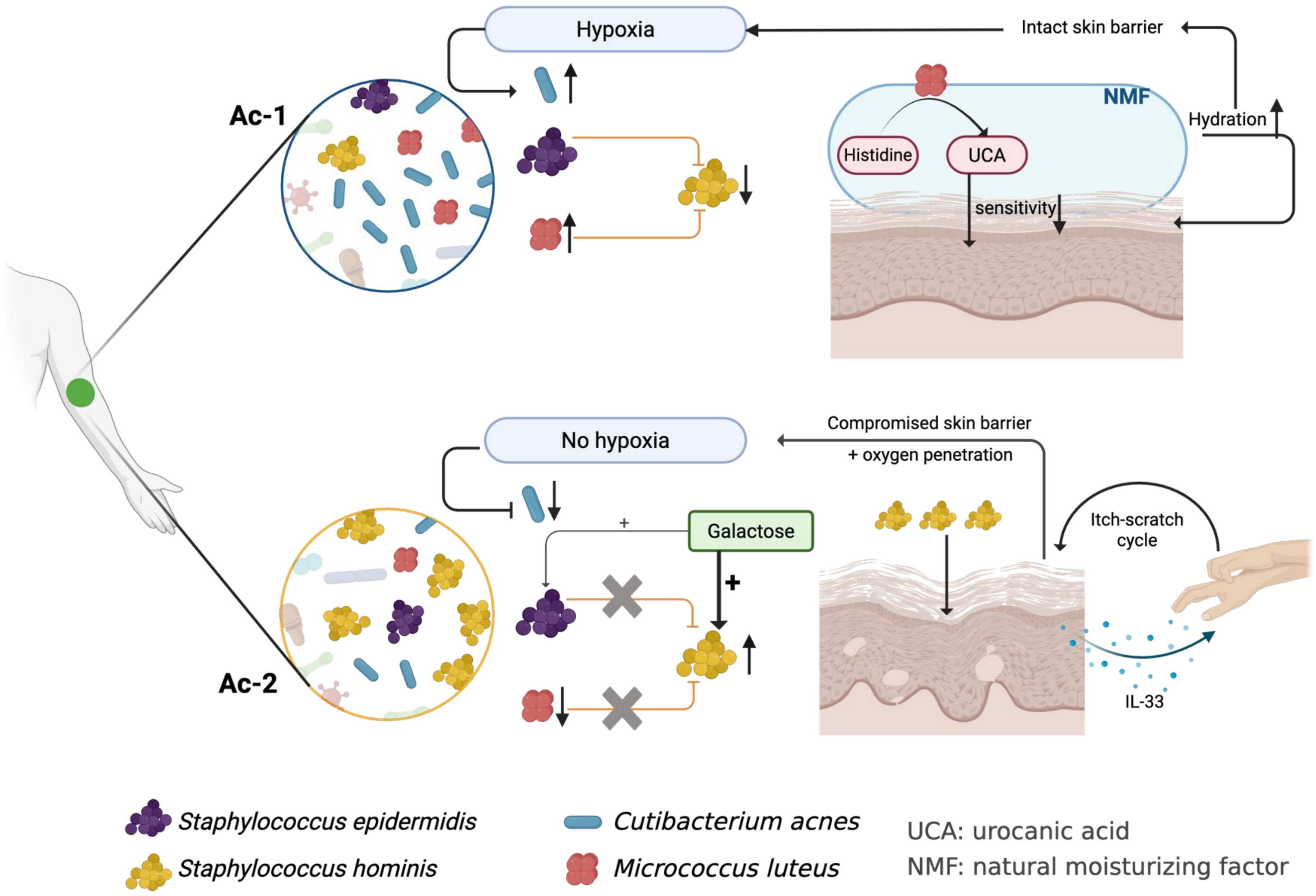


**Supplementary Figure 21: Schematic illustrating key species interactions that could impact dermotype structure, metabolic function and skin health in the elbow crease (Ac).**
